# Carbon isotope fractionation by an ancestral rubisco suggests biological proxies for CO_2_ through geologic time should be re-evaluated

**DOI:** 10.1101/2022.06.22.497258

**Authors:** Renée Z. Wang, Robert J. Nichols, Albert K. Liu, Avi I. Flamholz, Juliana Artier, Doug M. Banda, David F. Savage, John M. Eiler, Patrick M. Shih, Woodward W. Fischer

**Author notes:** **Corresponding author:** Renée Z. Wang, **Email:**.

## Abstract

The history of Earth’s carbon cycle reflects trends in atmospheric composition convolved with the evolution of photosynthesis. Fortunately, key parts of the carbon cycle have been recorded in the carbon isotope ratios of sedimentary rocks. The dominant model used to interpret this record as a proxy for ancient atmospheric CO_2_ is based on carbon isotope fractionations of modern photoautotrophs, and longstanding questions remain about how their evolution might have impacted the record. We tested the intersection of environment and evolution by measuring both biomass (ε_p_) and enzymatic (ε_Rubisco_) carbon isotope fractionations of a cyanobacterial strain (*Synechococcus elongatus* PCC 7942) solely expressing a putative ancestral Form 1B rubisco dating to ≫1 Ga. This strain, nicknamed ANC, grows in ambient pCO_2_ and displays larger ε_p_ values than WT, despite having a much smaller ε_Rubisco_ (17.23 ± 0.61‰ vs. 25.18 ± 0.31‰ respectively). Measuring both enzymatic and biomass fractionation revealed a surprising result—ANC ε_p_ exceeded ANC ε_Rubisco_ in all conditions tested, contradicting prevailing models of cyanobacterial carbon isotope fractionation. However, these models were corrected by accounting for cyanobacterial physiology, notably the CO_2_ concentrating mechanism (CCM). Our model suggested that additional fractionating processes like powered inorganic carbon uptake systems contribute to ε_p_, and this effect is exacerbated in ANC. Understanding the evolution of rubisco and the CCM is therefore critical for interpreting the carbon isotope record. Large fluctuations in that record may reflect the evolving efficiency of carbon fixing metabolisms in addition to changes in atmospheric CO_2_.

**Significance Statement:** Earth scientists rely on chemical fossils like the carbon isotope record to derive ancient atmospheric CO_2_ concentrations, but interpretation of this record is calibrated using modern organisms. We tested this assumption by measuring the carbon isotope fractionation of a reconstructed ancestral rubisco enzyme (>1 billion years old) *in vivo* and *in vitro*. Our results contradicted prevailing models of carbon flow in Cyanobacteria, but our data could be rationalized if light-driven uptake of CO_2_ is taken into account. Our study showed that the carbon isotope record tracks both the evolution of photosynthesis physiology as well as changes in atmospheric CO_2_, highlighting the value of considering both evolution and physiology for comparative biological approaches to understanding Earth’s history.

## Introduction

Photoautotrophs have evolved over geologic time to harness energy from the sun in order to ‘fix’ external, inorganic carbon (C_i_) into reduced, organic carbon (C_o_), thereby creating biomass for growth. Today, and likely for much of Earth’s history (1), the most widespread strategy for carbon fixation is the Calvin-Benson-Bassham (CBB) Cycle, where the key carbon fixation step is catalyzed by ribulose-1,5-bisphosphate (RuBP) carboxylase/oxygenase (rubisco) (2, 3). But rubisco’s central role in the CBB cycle and oxygenic photosynthesis poses a conundrum because it is usually considered to be a non-specific and slow enzyme. The first issue concerns rubisco’s dual carboxylase and oxygenase activities: the RuBP intermediate (enediolate) is susceptible to both O_2_ and CO_2_ attack (4). Consequently, instead of fixing a CO_2_ molecule during photosynthesis, rubisco can instead assimilate O_2_ to yield 2-phosphoglycolate (2-PG), which is not part of the CBB cycle and therefore must be salvaged through photorespiratory pathways that consume ATP, reducing power, and carbon (5). The second issue concerns rubisco’s maximum carboxylation rate (*V_C_*), which is ≈7-10 times slower than other central metabolic enzymes (6), and displays very limited variation across large phylogenetic distances (7).

Both issues—its dual carboxylase / oxygenase activity and limited maximum carboxylation rate—are typically rationalized by considering its evolutionary history in the context of long-term changes in environmental CO_2_ and O_2_ concentrations. Rubisco is thought to have evolved at a time when there was trace O_2_ and much higher CO_2_ concentrations in the atmosphere, in contrast to the modern atmosphere where O_2_ is roughly 20% while CO_2_ is only about 0.04% by partial pressure (1). In addition, rubisco is thought to have been the primary carboxylating enzyme of global photosynthesis since the Great Oxygenation Event, and potentially far prior (1).

Likely in response to these changing environmental concentrations, many aquatic photoautotrophs have evolved CO_2_ concentrating mechanisms (CCMs) that concentrate CO_2_ around rubisco in order to enhance carboxylation and suppress oxygenation. Even with CCMs, the effective *in vivo* rates of extant rubiscos are estimated to be much lower (≈1% for terrestrial and ≈15% for marine rubiscos) than their maximal catalytic rates observed in lab at 25°C, likely due to rubisco not working at night and the lower temperature of marine environments (2). However, all known Cyanobacteria today have CCMs, as do many bacterial chemolithoautotrophs, many aquatic algae, and some plants (8). The bacterial CCM has two main components: i) C_i_ pumps producing high cytosolic HCO_3_^-^ concentrations, and ii) co-encapsulation of carbonic anhydrase (CA) and rubisco inside proteinaceous organelles known as carboxysomes (Figure 1A) (9–11). These powered C_i_ pumps include BCT1 (ATP-dependent powered HCO_3_^-^ transporter), SbtA (Na^+^/HCO_3_^-^ symporters), BicA (Na^+^-dependent HCO_3_^-^ transporter), NDH-1MS and NDH-1MS’ (NADPH-dependent powered CO_2_ uptake; see (12) for review). There are competing arguments in the literature for when the CCM evolved, ranging from the Proterozoic to the Phanerozoic Eon (8, 13). Therefore, for up to half of Earth’s history, bacterial rubiscos have functioned in concert with a system that pumps C_i_ into and around the cell.

**Figure 1:**
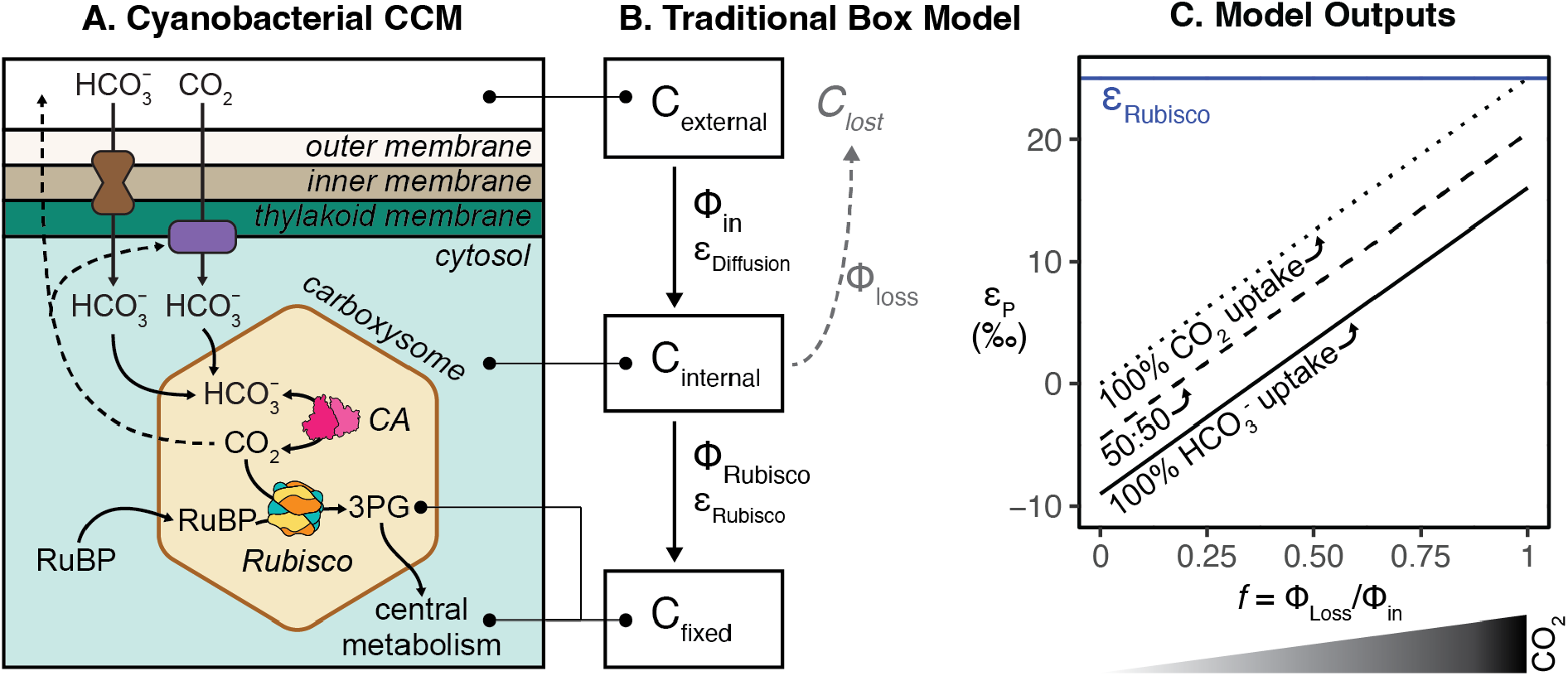
Comparing the Cyanobacterial CO_2_ Concentrating Mechanism (CCM) to the traditional box model of photosynthetic C isotope discrimination. A) Cyanobacterial CCMs rely on i) active C_i_ uptake into the cell, and ii) co-encapsulation of carbonic anhydrase (CA) and rubisco within the carboxysome. Independent, powered transporters for HCO_3_^-^ and CO_2_ are shown in brown and purple; both work to increase cytosolic concentrations of HCO_3_^-^ (see (12) for review). All CCM components work to produce a high carboxysomal CO_2_ concentration that enhances CO_2_ fixation by rubisco and suppresses oxygenation. Limited CO_2_ escapes from the carboxysome – some is scavenged by CO_2_ pumps while the rest leaves the cell. B) Architecture of the traditional box model based on (16–19); see Supplemental for full discussion of this model. Boxes denote carbon pools of interest, and fluxes between boxes are denoted by Φ. Each flux has its own isotopic fractionation denoted by ε; no fractionation is assumed for Φ_loss_. Model assumes an infinitely large external carbon pool, that carbon not fixed by rubisco (C_lost_) returns to this pool, and that fluxes are at steady state. Note that this architecture does not include a box for the carboxysome. C) Model solution for the traditional model is ε_P_ = *a*\*ε_equil_ + *f*\*ε_Rubisco_, where where ε_P_ is defined as the difference in *δ*^13^C of C_external_ and C_fixed_, *f* is defined as the fraction of C_i_ lost (Φ_loss_/Φ_in_), and *a* is the fractional contribution of HCO_3_^-^ to total C_i_ uptake. When *a* = 0, all C_i_ uptake is as CO_2_ (dotted line); when *a* = 1, all C_i_ uptake is as HCO_3_^-^ (solid line). This model is presented in (20), which is a generalization of (21) that accounts for the fact that C_i_ uptake (Φ_in_ in Panel B) ranges in composition between CO_2_ and HCO_3_^-^ based on which C_i_ uptake system is used. Values of ε_Rubisco_ = 25‰ and ε_equil_ = −9‰ were used for this illustration (22). Model outputs indicate that at high external CO_2_ concentrations (dark wedge under graph) there is excess CO_2_ that rubisco cannot use, causing net C_i_ leakage (larger *f* values) from the cell.

However, rationalization of rubisco’s evolutionary history is highly dependent on our understanding of past environments, e.g. atmospheric CO_2_ and O_2_ concentrations. For the vast majority of Earth’s history, we must rely on chemical fossils like the carbon isotope record for this purpose. The carbon isotope record is composed of measurements of the relative ratios of ^13^C to ^12^C isotopes in C-bearing phases of sedimentary rocks over time. Contemporaneous C_i_ pools are preserved as carbonate salts (like limestones and dolomites), while contemporaneous biomass and Co pools are preserved in the organic-rich components (typically kerogen) of many different lithologies and are measured as rock total organic carbon (TOC) (14). The difference in C-isotope ratios between organic samples and an inorganic reference is typically reported using delta notation (*δ*^13^C) and expressed in per mil (‰, see Methods). Currently, carbon isotope data has been assembled globally from myriad environments to cover ≈3.8 billion years (Ga) of Earth’s 4.5 Ga history; this data is viewed as a record of both inorganic and organic carbon cycle processes over geologic time (15).

The carbon isotope record is particularly important for constraining ancient atmospheric pCO_2_ (23, 24) because direct observations of the past atmosphere from ice cores only extends back ≈1 million years (25). One notable feature of the record from ≈3.8 Ga to the present is that the *δ*^13^C of C_o_ is depleted in ^13^C by ≈25‰ compared to C_i_ (14, 15, 26), and this offset roughly matches the carbon isotopic fractionation of known carbon-fixing metabolisms and enzymes in the modern. (Note that the convention for reporting ε in this field is the opposite of other geochemistry fields – here, a negative value indicates a relative ^13^C enrichment, in contrast to other fields where negative values mean ^13^C depletion.) Rubisco displays a kinetic isotope effect (KIE) where it preferentially fixes ^12^CO_2_ over ^13^CO_2_ due in part to the *V_C_* being slightly faster for ^12^CO_2_ than ^13^CO_2_ (27), which causes the reaction product, 3-phosphoglycerate (3-PG), to be relatively depleted in ^13^C by several percent (tens of ‰) relative to the isotopic composition of the initial CO_2_ substrate. The difference in *δ*^13^C of the CO_2_ substrate and the 3-PG product is typically reported as ε_Rubisco_ and varies between 18-30‰ for several extant rubiscos (26, 28), with the exception of the coccolithophore *Emiliania huxleyi* with ε_Rubisco_ ≈ 11‰ (29). Because all biomass is synthesized from 3-PG in autotrophs utilizing the CBB cycle, biomass is depleted in ^13^C compared to external C_i_ pools and the magnitude of this difference is called ε_P_. There is an additional fractionation factor associated with the preservation of biomass and C_i_ as rocks, so the magnitude of fractionation between C_i_ and C_o_ pools measured from the rock record is termed ε_TOC_ and varies slightly from ε_P_ (30). Therefore, if one can accurately derive ε_P_ from the rock record (ε_TOC_) and pair it with some model relating ε_P_ to pCO_2_, one could learn about both the evolution of photosynthetic physiology and abiotic changes in the carbon cycle over geologic time.

The dominant model used today for this purpose (“C Isotope Record Model,” Figure S7) is:

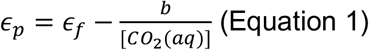

ε_f_ is the maximum possible isotopic fractionation of photosynthetic carbon fixation and is typically set to ε_Rubisco_. The term *b* (‰ kg μM^-1^) is empirically fit from pure culture experiments of eukaryotic and bacterial algae, and encompasses all physiologic effects that may affect isotopic fractionation like the CCM, growth rate, cell size and geometry, membrane permeability, growth media composition (e.g. pH, salinity, limiting nutrient), strain genetics and physiological state (31–35). [CO_2_(aq)] is the concentration of dissolved CO_2_ in solution around the cells, and in the limit of high [CO_2_(aq)], the term *b*/[CO_2_(aq)] goes to zero and ε_P_ = ε_f_, which is assumed to be ε_Rubisco_. Therefore, the maximum value of ε_P_ is ε_Rubisco_, and the term *b* sets how quickly ε_P_ approaches the limit of ε_Rubisco_.

The C Isotope Record model has such a limit because the laboratory studies of plants and algae that it is based on (“Traditional model,” Figure 1) shows such a limit. The traditional model was originally based on studies of carbon isotope fractionation in plants (dotted line in “Traditional model” in Figure 1C; all C_i_ uptake is as CO_2_ for plants) and was later adapted to eukaryotic and bacterial algae. The primary architecture of this model stems from a seminal study by Park and Epstein (18) who proposed a “two step model” to explain ε_P_ of tomato plants grown in varied CO_2_ concentrations and light levels. In this model, carbon can be viewed as residing in one of three pools, or “boxes” (Figure 1B) - C_i_ outside the cell (C_ext_), C_i_ inside the cell (C_internal_), or C_o_ as biomass (C_fixed_). A “leakiness” term, *f*, is defined as the ratio of fluxes (Φ) of C_i_ exiting or entering the plant, where all of the C_i_ that entered the cell is lost when *f*=1. In this simplified model, ε_p_ is determined by the isotopic effect of two distinct steps: i) the diffusion of CO_2_ into the plant (ε_Diffusion_; <1‰ across a diaphragm cell in water at 25°C (36)); and ii) the carbon fixation step catalyzed by rubisco (ε_Rubisco_; ≈18-30‰). Notably, Park and Epstein proposed that the isotopic fractionations of these two steps are not additive *in vivo* (i.e. ε_p_ ≠ ε_Diffusion_ + ε_Rubisco_) but instead reflects the process by which photosynthesis is limited, either diffusion of CO_2_ into the cell (ε_p_ = ε_Diffusion_) or CO_2_ fixation by rubisco (ε_p_ = ε_Rubisco_) (18).

This physiological interpretation results from the model solution, which is usually solved by assuming steady state and results in a linear relationship between ε_p_ and *f* where the minimum and maximum ε_p_ values are ε_Diffusion_ and ε_Rubisco_ respectively (Figure 1C). This allows experimentally measured values of ε_p_ to then be used to solve for CO_2_ leakage (*f*, Figure 1C). Therefore, the corresponding physiological interpretations at the minimum and maximum model limits are when ε_p_ ≈ ε_Diffusion_, nearly all carbon entering the cell is used and with this mass balance constraint rubisco’s ^12^C preference is not “expressed”; conversely, when ε_p_ ≈ ε_Rubisco_, very little of the carbon entering the cell is fixed (f ≈ 1, i.e. nearly all of the carbon leaks from the cell) and rubisco can “choose” between ^12^C and ^13^C substrates so that rubisco’s KIE is fully expressed. Farquhar et al. (19) later derived a relationship between ε_p_ and the ratio of external vs. intracellular CO_2_ partial pressures, allowing CO_2_ concentrations at the site of rubisco to be estimated from ε_p_. Therefore, given the assumption that C_i_ is taken up passively, it is possible to derive an increasing relationship between Cext and ε_P_ from this model, where large ε_P_ suggests that high external CO_2_ concentrations are creating excess CO_2_ at rubisco and ultimately causing more CO_2_ to leak out of the cell than can be fixed (see Supplemental and (17)).

This model was later adapted to algae to account for CCMs – mainly active uptake of C_i_ as HCO_3_^-^ and/or CO_2_ – and physiological parameters including growth rate and cell geometry (21, 31, 32, 37, 38). These studies grew eukaryotic and bacterial algae in a range of pCO_2_ and culturing conditions to test if the linear relationship between ε_p_ and [CO_2_] observed in plants still holds. Interestingly, cyanobacterial ε_p_ was found to be roughly constant independent of environmental pCO_2_ and growth rate (31). In addition, because measured cyanobacterial ε_p_ values were less than known cyanobacterial ε_Rubisco_ values, additional isotopic fractionation factors were not needed to explain ε_p_, even though some active C_i_ transport processes, which may fractionate carbon isotopes, were known in Cyanobacteria at the time (39–41). Therefore, though different versions of this model exist, all variations essentially modified the plant model by shifting the y-intercept of Fig. 1C to account for uptake of HCO_3_^-^ in addition to CO_2_. If C_i_ entering the cell is primarily CO_2_, the model effectively represents plants (dotted line in Fig. 1C). If C_i_ entering the cell is primarily HCO_3_^-^ through active uptake, as in many algae, all values are shifted to lower ε_p_ values (solid line in Fig. 1C) because of the equilibrium isotopic effect (ε_equil_) between CO_2_ and HCO_3_^-^ (≈ −9‰ (22)). In Figure 1C, we plot the traditional model as derived in Eichner et al. (20), which is an adaptation of (21):

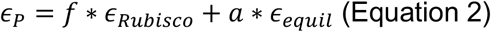

Here ε_equil_ is the equilibrium isotope effect, and *a* is the fraction of C_i_ entering the cell as CO_2_ (*a*=0) or HCO_3_^-^ (*a*=1); the diffusion isotope effect (ε_Diffusion_) is considered negligible. See Supplementary Information for further discussion of the “traditional” model.

Overall, current models relating pCO_2_ and autotrophic carbon isotope fractionation have a limit where ε_p_ cannot exceed ε_Rubisco_ (Figure 1C). Yet, the largest ε_P_ values observed in the Archaean Eon exceed 30‰ (14, 15) and also exceed all current measurements of ε_Rubsico_ (for recent compilation see (26)). In addition, recent studies in dinoflagellates have shown that ε_p_ can regularly exceed ε_Rubsico_ under certain growth conditions (for review see (28)), and detailed studies of Cyanobacteria imply that leakage estimates derived from ε_p_ are not physiologically possible (20). These studies have therefore motivated updated models of carbon isotope fractionation in algae that account for the isotopic fractionations associated with different C_i_ uptake mechanisms (20, 28).

In addition to taking modern physiology into account, it is also important to understand how the evolution of rubisco and the CCM may have affected the carbon isotope composition of biomass and therefore *δ*^13^C values of C_o_ preserved in the rock record. Recent studies have addressed this issue directly by testing model organisms that may better resemble an ancestral counterpart, including a cyanobacterial strain lacking a CCM (42), a cyanobacterial strain that overexpresses rubisco (43), and a cyanobacterial strain expressing an inferred ancestral rubisco dating from ≈1-3 Ga (44, 45).

Here, we measured the ε_p_ of a control strain of *Synechococcus elongatus* PCC 7942 expressing the wild-type rubisco (NS2-KanR, referred to as ‘WT’, see Methods), as well as a strain, ‘ANC’, engineered to express an inferred ancestral Form 1B enzyme (dating to >1 Ga) as its sole rubisco (46) in varied CO_2_ and light conditions. This putative ancestral rubisco was previously purified and its kinetics were characterized *in vitro*. Its sequence was then inserted into the genome of a modern cyanobacterium, though the genome of the strain in that study contained both extant and ancestral rubisco sequences (46). Here we study a strain where the extant sequence has been fully deleted and replaced with the reconstructed ancestral sequence. In addition, we measured ε_Rubsico_ of the present-day and ancestral rubiscos *in vitro*. We observed that: i) biomass ε_p_ is greater for ANC than for its WT counterpart for all conditions tested, even though ANC ε_Rubsico_ (17.23 ± 0.61‰) is considerably less than WT ε_Rubsico_ (25.18 ± 0.31‰); ii) ANC ε_p_ exceeds ε_Rubsico_ in all tested conditions even though the traditional model sets the maximum possible ε_p_ = ε_Rubsico_; iii) ANC ε_p_ increases with light levels while WT ε_p_ increases with CO_2_; iv) ANC displays a growth defect at ambient pCO_2_ that is rescued at high pCO_2_; and v) ANC growth is severely inhibited in high light. Consistent with recent studies of eukaryotic algae (20, 28), ANC ε_p_ exceeding ε_Rubsico_ implies that the traditional box model is incomplete and additional isotopic fractionations are needed. In addition, modulation of ε_p_ with light suggests that aspects of cyanobacterial physiology beyond the CBB cycle must be taken into account to explain how ε_p_ can vary independently of CO_2_. We posit additional factors related to C_i_ uptake that might explain fractionation measurements that deviate from box model predictions in both extant and ancient organisms.

## Results & Discussion

### Ancestral rubisco enzyme fractionates less than WT rubisco enzyme

We measured the carbon isotope fractionations of WT and ANC rubiscos *in vitro* using the substrate depletion method ((47–50); see Methods). The kinetics of this putative ancestral rubisco were previously characterized *in vitro* and are summarized in Table 1 (46). Previous work on rubisco isotope discrimination predicted that ε_Rubsico_ should correlate positively with specificity (S_C/O_), a unitless measure of the relative preference for CO_2_ over O_2_ (51). We therefore expected ANC and WT ε_Rubsico_ values to be the same within uncertainty because of their similar S_C/O_ values, but found that ANC ε_Rubsico_ (17.23 ± 0.61‰) was about 8‰ less than WT ε_Rubsico_ (25.18 ± 0.31‰, Table 1).

**Table 1:** Rubisco characteristics. Starred values (*) for the modern Form 1B were measured in rubiscos purified from *Synechococcus* sp. PCC 6301, which has the same small and large subunit (*RbcS, RbcL*) sequences as our working WT strain, *Synechococcus* sp. PCC 7942 (46). Kinetic isotope effect (ε_Rubisco_, avg. ± s.e.) was measured in this study using the substrate depletion method (47–50). Carboxylation turnover rate under substrate-saturated conditions (V_C_); Michaelis constant for CO_2_ in ambient levels of O_2_ (K_C_^Air^); the catalytic efficiency towards CO_2_ in ambient air (V_C_/K_C_^Air^); and specificity, a unitless measure of the relative preference for CO_2_ over O_2_; (S_C/O_) are from (46).

| Rubisco | $\epsilon_{\text{Rubisco}}$ (‰) | $V_c$ (s<sup>-1</sup>) | $K_c^{\text{Air}}$ (μM) | $V_c/K_c^{\text{Air}}$ (s<sup>-1</sup>mM<sup>-1</sup>) | $S_{\text{C/O}}$ |
| --- | --- | --- | --- | --- | --- |
| Ancestral Form 1B | $17.23 \pm 0.61$ | $4.72 \pm 0.14$ | 168.7 | 28 | $49.6 \pm 1.8$ |
| Modern Form 1B | $25.18 \pm 0.31^*$ | $9.78 \pm 0.48^*$ | 184.1* | 53.1* | $50.3 \pm 2.0^*$ |

### The ANC strain fractionates more than WT

Working in *S.elongatus* PCC 7942, we produced a mutant strain lacking the native Form 1B rubisco and expressing instead an ancestral Form 1B rubisco produced by computational ancestral sequence reconstruction (46) as its sole rubisco enzyme. We then grew this strain, termed ANC, and a control strain, termed wild-type or ‘WT’ (see Methods), in a variety of light and CO_2_ levels: i) A reference condition (ambient pCO_2_ of 0.04% v/v, standard light flux (120 μE)); ii) High CO_2_ (5% pCO_2_, 120 μE); iii) High light (0.04% pCO_2_, 500 μE). The CO_2_ gas at ambient and high CO_2_ conditions had *δ*^13^C values of −12.46‰ and −36.84‰ respectively.

Counter to expectations based on ε_Rubisco_ (Table 1), ANC ε_P_ was as large or larger than WT ε_p_ in all conditions tested (Figure 2). This was consistent with recent results from a similar ancestral analog, where the ancestral analog’s ε_p_ values exceeded WT in ambient and elevated CO_2_ levels (44). In this study, the highest ANC ε_p_ values were observed for cultures grown in high light, where growth was comparatively slow (doubling time ≈ 50 hours, Figure 3 and Table S3). ANC ε_p_ values were also modulated by light and CO_2_ differently than WT. Compared to the reference condition, WT ε_P_ values were indifferent to high light and only increased in high CO_2_ (Figure 2A). In contrast, ANC ε_P_ values did not increase in high CO_2_ and only increased in high light (Figure 2B). This result contrasted with the ancestral analogue in (44) where ε_p_ values increased by ≈10% at 2% CO_2_.

**Figure 2:**
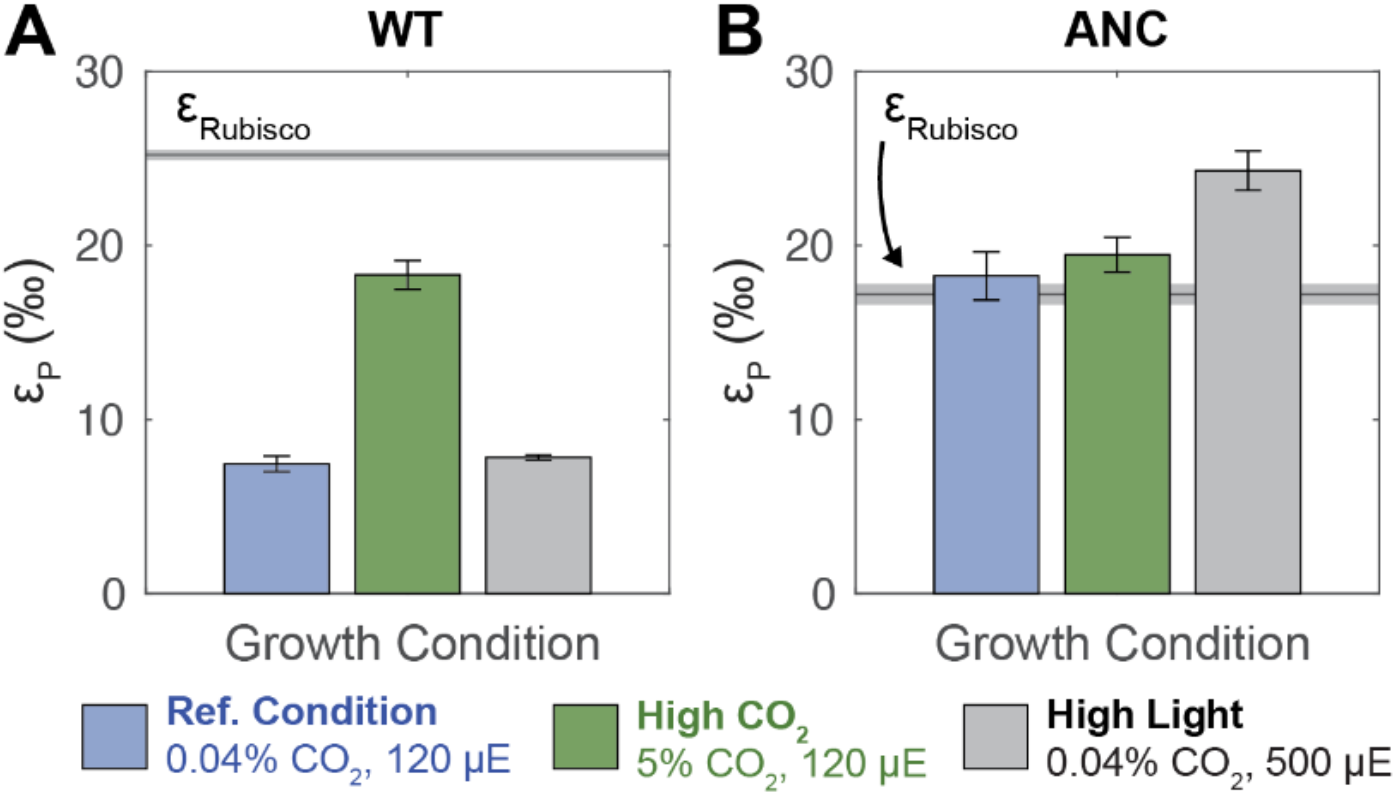
Whole cell carbon isotope fractionation by WT and ANC strains. ε_P_ (‰) values (avg. ± s.e.) for A) WT and B) ANC strains across growth conditions.. For each strain, the maximum ε_P_ possible based on the traditional model (ε_P_ = ε_Rubisco_) is shown as a gray line (avg. ± s.e.). Most measured ANC ε_P_ values exceed the theoretical limit (ε_P_ > ε_Rubisco_ + s.e.), while WT ε_P_ values do not (ε_P_ < ε_Rubisco_). WT ε_P_ values increase in response to elevated CO_2_ concentrations, while ANC ε_P_ values increase in response to elevated light flux. See Table S3 for full results.

**Figure 3:**
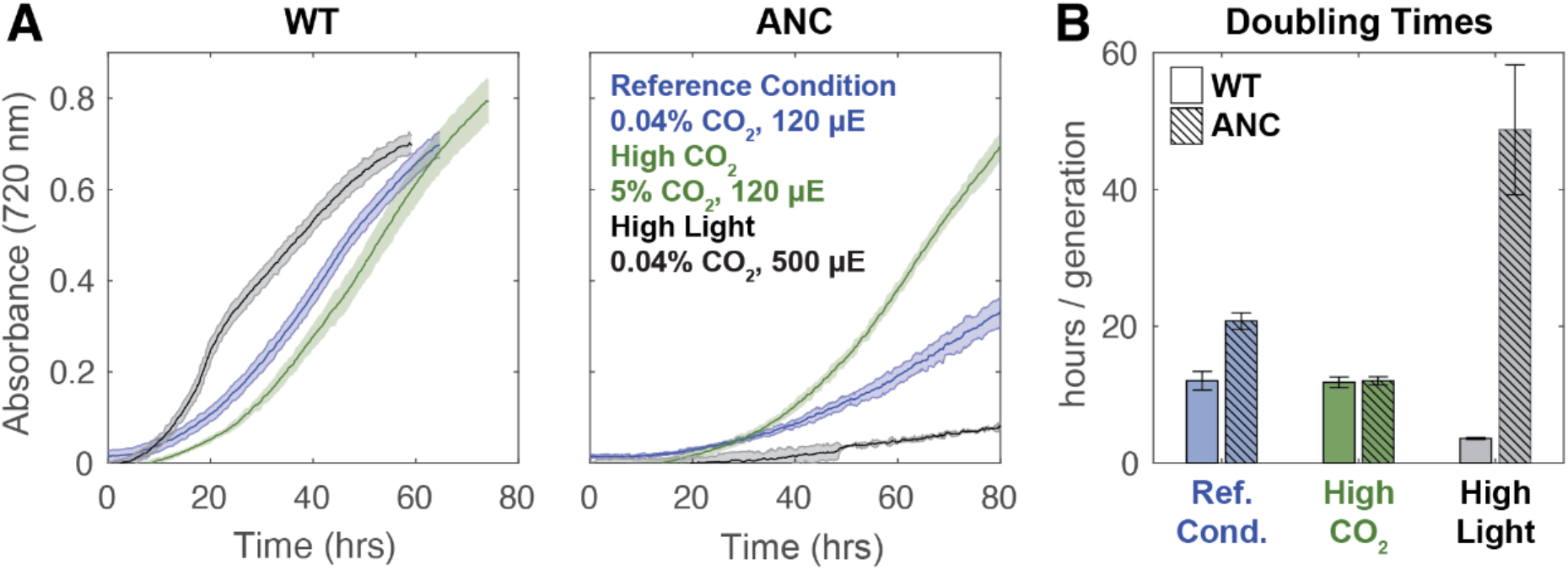
Growth curves for WT and ANC strains across experimental conditions. A) Averaged growth curves shown for WT and ANC strains to 80 hours, colored by growth condition as indicated in figure. Data was smoothed with a rolling median (Methods); see full ANC growth curves in Supplemental Fig. S12. B) Average doubling times with standard deviations. See Supplemental for details of doubling time calculation. ANC displayed a growth defect relative to the WT at the reference condition, which was rescued by high CO_2_. ANC grew slowest in high light, while WT grew fastest in that condition.

In addition, the traditional box model described above cannot accommodate ε_p_ values in excess of εRubsico (Figure 1C). However, average ANC ε_P_ values exceeded ANC εRubsico in all growth conditions (Figure 2), particularly under high light conditions where the largest difference was seen (ε_p_ = 24.30 ± 0.12‰ vs εRubsico = 17.23 ± 0.61‰). The traditional box model also states that ε_P_ values are solely modulated by changing external pCO_2_ concentrations, which cannot accommodate the ANC ε_P_ observations.

### Ancestral rubisco strain grows at ambient CO_2_ concentrations

Remarkably, the ANC strain managed to grow in ambient pCO_2_ and standard light conditions (Figure 3A), even though the ancestral rubisco has a *V_C_* roughly half that of WT (Table 1). This implies that its rubisco enzyme is properly encapsulated in the carboxysome, since it is well established that improper carboxysome formation greatly inhibits growth in ambient pCO_2_ (52, 53). Mutant strains that are unable to form carboxysomes cannot grow in ambient air (53). Indeed, electron micrographs of WT and ANC cells grown in ambient CO_2_ and light conditions (Methods) show multiple carboxysomes per cell in both strains (Figure 4 and Figure S13). Rubisco density can be seen within some of the carboxysomes (Figure 4C). In addition, the rubisco residues necessary for protein interactions mediating ***β***-carboxysome encapsulation were recently identified (54). We aligned WT and ANC rubisco sequences and found that fourteen of the sixteen residues are conserved in the ancestral sequence (Tables S8-9, Figure S14). In addition, WT and ANC strains harvested during exponential growth in the reference condition exhibit similar photosystem stoichiometry, as indicated by absorbance spectra (Figure S15). Taken together, these data indicated that carboxysomes form in ANC and the ancestral rubisco is encapsulated within these structures. Further strengthening our inference that the ancestral sequence is compatible with *β*-carboxysome formation, a similar ancestral analogue was also found to grow in ambient air (44).

**Figure 4:**
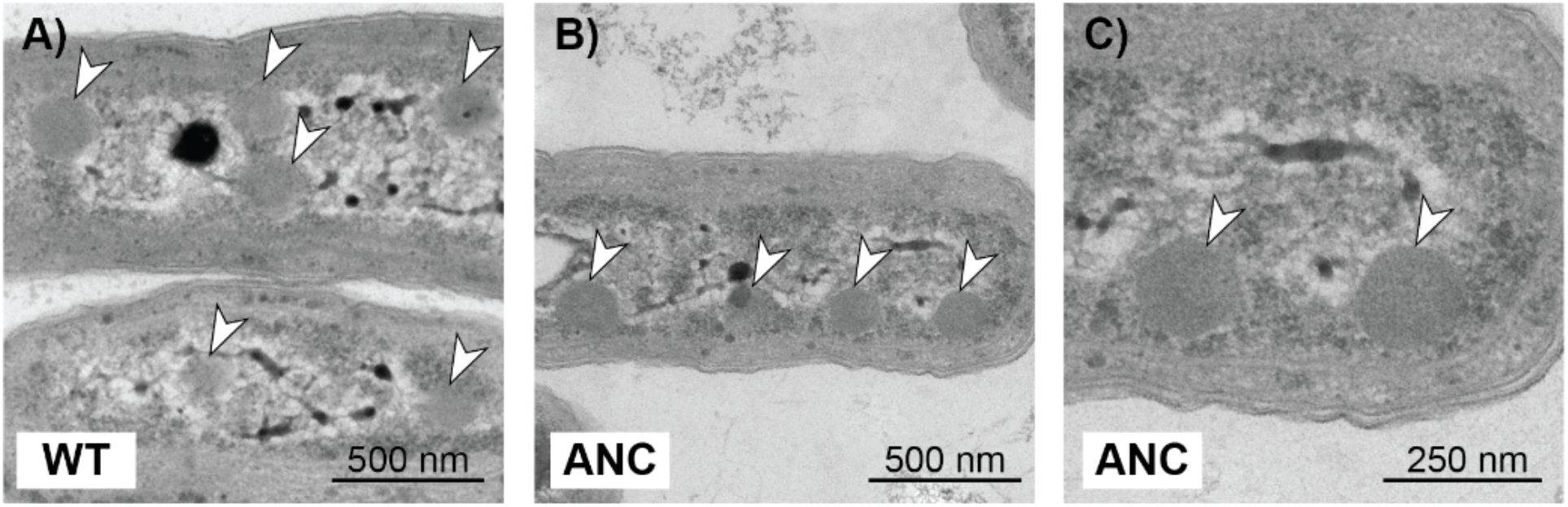
WT and ANC strains both produce carboxysomes at ambient pCO_2_. Transmission electron micrographs of WT (A) and ANC (B,C) strains that were harvested during exponential growth in the reference condition (ambient pCO_2_, standard light flux). Both strains show multiple carboxysomes per cell, as indicated by white arrows, and carboxysomes exhibit the typical hexagonal shape (53). C) is the same image as in B) but magnified to show that rubisco density seen can be within the carboxysomes of ANC. The dark internal body in A) is likely a polyphosphate body (55). See Figure S13 for additional images.

In addition, the difference in *V_C_* between the ancestral and modern rubiscos was mirrored in the doubling times of WT and ANC strains (Figure 3B, Table S3), where ANC doubling times were roughly twice that of WT in the reference condition (20.8 ± 1.2 vs. 12.0 ± 1.4 hours respectively). In addition, the carboxylation efficiency in ambient air (V_C_/K_C_^Air^) for the ancestral Form 1B rubisco, which measures the enzyme’s ability to carboxylate in conditions with low CO_2_ and relatively high O_2_, is roughly half that of the modern Form 1B rubisco as well (Table 1). This suggested that ANC’s growth was limited by its ability to fix CO_2_ from ambient air. This growth defect was ameliorated by high pCO_2_, where doubling times for both strains were the same within uncertainty (WT 11.8 ± 0.8 hours; ANC 12.0 ± 0.6 hours), though we observed a longer lag phase for ANC. WT doubling times were the same within uncertainty for the reference and high CO_2_ conditions (12.0 ± 1.4 vs. 11.8 ± 0.8 hours respectively), consistent with previous studies where increased pCO_2_ did not affect growth rate (56). In contrast, elevated CO_2_ greatly accelerated the growth of ANC, reducing its doubling time from ≈21 to ≈12 hrs, supporting our inference that CO_2_ availability is limiting the growth of ANC and implicating the CCM in its growth defect. Consistent with our results, a similar ancestral Form 1B analogue displayed total carboxylase activity roughly half that of the modern Form 1B (57).

We observed the greatest differences in doubling times between ANC and WT when the strains were grown at high light (Figure 3, Table S3). In these conditions, WT cultures were a dark, blue-green color typical of healthy cyanobacterial cells while ANC cultures were yellow-green (Fig. S11), suggesting degradation of phycobilisomes via a known starvation pathway to reduce the cell’s capacity for light harvesting and photochemical electron transport (58). We therefore infer that ANC could not fix CO_2_ at a rate matching its light harvesting capability, and hence invoked this regulatory pathway to decrease light harvesting capacity. WT, in contrast, grew rapidly in the high light condition.

### Proposed influence of a light-powered, vectoral carbonic-anhydrase

As discussed above, recent studies in extant bacterial and eukaryotic algae have shown that ε_p_ can regularly exceed ε_Rubsico_ under certain growth conditions (for review see (28)), motivating updated models of carbon isotope fractionation in both eukaryotic and bacterial algae (20, 28, 59). These models suggested that ε_P_ values could only be explained if another enzyme acting as a CA catalyzing an energy-coupled vectoral hydration of intracellular CO_2_ to HCO_3_^-^ was taken into account, since this reaction is calculated to have a large isotopic effect and would therefore allow ε_p_ to exceed ε_Rubisco_ (20, 28, 59). Though the cell does not “know” where CO_2_ in the cytosol came from, these models primarily invoke such an enzyme for internal C_i_ recycling, where CO_2_ lost from the carboxysome could be converted to HCO_3_^-^ so that it could remain in the cell (20, 28, 59). However, energy-coupled CAs can also serve as CO_2_ uptake systems by converting extracellular CO_2_ that passively translocates the membrane to intracellular HCO_3_^-^ (Figure 1A), which is advantageous in conditions (e.g. acidic pH) where CO_2_ is the dominant form of extracellular C_i_ (10, 60, 61).

In general, Cyanobacteria have been shown to have two modes of active C_i_ uptake: uptake of hydrated C_i_ (predominantly H_2_CO_3_ and HCO_3_^-^) and uptake of CO_2_ (61). In order for the CCM to function, either mode would need to produce a high, non-equilibrium concentration of HCO_3_^-^ in the cytoplasm (8, 10). This is thought to be achieved by coupling CA to an energy source (e.g. light or an ion gradient) that drives the vectoral hydration of CO_2_ to HCO_3_^-^ in the cytoplasm (62). There is now excellent data supporting this hypothesis in Cyanobacteria, where accessory proteins that bind to the NDH complex, the cyanobacterial homolog of the respiratory Complex I NADH-dehydrogenase, are known to mediate CO_2_ uptake specifically (63–65). Additionally, one of these accessory proteins, CupA/B, is reminiscent of a CA and contains a telltale zinc active site situated near a proton channel in a membrane subunit (66). The prevailing understanding of these data is, therefore, that these complexes couple inorganic carbon uptake to energy supplied by photochemical electron transport. Moreover, a similar protein complex has been described in proteobacterial chemoautotrophs, suggesting that energy-coupled CO_2_ hydration is widespread (60).

A vectoral CA would affect ε_p_ for two reasons. First, CO_2_ and HCO_3_^-^ are isotopically distinct. At equilibrium in standard conditions, HCO_3_^-^ is ≈8L more enriched in ^13^C than CO_2_ (67, 68). Therefore, if a cyanobacterium is predominantly taking up CO_2_, the internal C_i_ pool from which biomass is formed would be isotopically lighter (^13^C-depleted) than if HCO_3_^-^ is the dominant source of C_i_. Second, unidirectional CO_2_ hydration is expected to impart a substantial isotope effect, with calculated values ranging from ≈19 to 32L (67, 69–72). Therefore, there are two mechanistic reasons that ε_P_ could exceed ε_Rubisco_ in conditions where energized CO_2_ uptake and hydration is active. Indeed, a recent model of C-isotope fractionation in Cyanobacteria specifically invoked the NDH complex to rationalize ε_p_ values that exceed ε_Rubisco_ (20).

Because this energy-coupled CO_2_ uptake and hydration by the NDH complex is driven by light energy, e.g. via cyclic electron flow around photosystem I (66), and because the vectoral hydration of CO_2_ to HCO_3_^-^ is thought to have a large carbon isotope fractionation, we hypothesized that ε_p_ would increase with light intensity. Indeed, we observed the largest ANC ε_P_ values, far exceeding ANC ε_Rubisco_, in the high light condition and found that ANC ε_P_ varies primarily with light and not CO_2_ (Figure 3). This observation is counter to the traditional model which proposes ε_P_ as a direct correlate of external pCO_2_ (16, 17). Furthermore, on short timescales (≈minutes) cyanobacterial C_i_ uptake can be modulated by light intensity alone, fully independent of external C_i_ concentrations (73), and CO_2_ uptake can occur in the absence of carbon fixation (74, 75). Based on these physiological and isotopic observations, our study also supports the hypothesis that a powered, vectoral CA like the NDH complex is likely active in Cyanobacteria, and is likely responsible for ε_p_ > ε_Rubisco_ in ANC.

### Proposed model for carbon isotope fractionation in Cyanobacteria

As discussed above, the traditional box model cannot produce ε_p_ > ε_rubisco_ (Figure 1C). In this model, the C_i_ leakage term (*f*) is fit from measured ε_p_ values and *f* = 1 implies that all carbon uptake leaks out of the cell. Though the traditional box model can accommodate both CO_2_ and HCO_3_^-^ uptake, which differ in their equilibrium isotopic composition, even modeling 100% CO_2_ uptake gave *f* > 1 for ANC in all conditions (Figure 5A and S8). Yet, ANC grew reproducibly in all conditions tested (Figure 3). We also encountered challenges using the traditional model to rationalize WT data: fitting the model gave *f* < 1 in ambient pCO_2_ conditions, but high CO_2_ conditions yielded *f* > 1 unless all C_i_ uptake was assumed to be as HCO_3_^-^ (see Figure S8 and Supplemental Information for discussion). Therefore, to rationalize our results for both WT and ANC, we developed a box model that represents a small modification of the traditional model.

**Figure 5:**
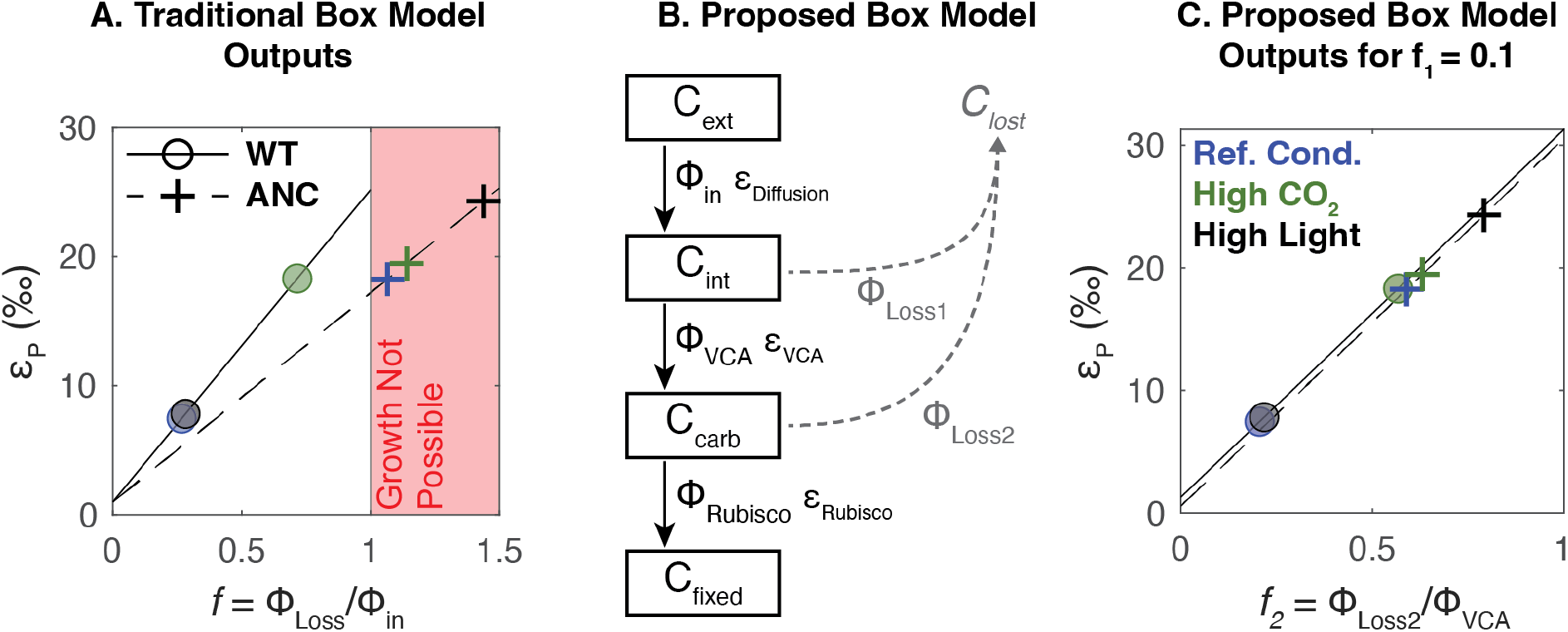
Proposed box model based on experimental results. A) Experimental results (circles and crosses) plotted onto traditional box model outputs (dashed and solid lines) for WT and ANC respectively if C_i_ uptake is all CO_2_. Uncertainties are smaller than data points. Colors indicate growth conditions: blue = reference condition (0.05% pCO_2_ (v/v), 120 μE); green = high CO_2_ (5% pCO_2_ (v/v), 120 μE); black = high light (0.05% pCO_2_ (v/v), 500 μE). *f* is as defined in Figure 1; region where *f* > 1 is shaded in red. B) Proposed box model architecture, with main carbon pools of interest in boxes. Subscripts indicate external (*ext*), internal (*int*), carboxysome (*carb*), and fixed (*fixed*) carbon pools. Fluxes are denoted by Φ where subscripts indicate fluxes into the cell (*in*), out of the cell (*Loss1, Loss2*), into the carboxysome (*VCA* for *V*ectoral *C*arbonic *A*nhydrase), and into fixed biomass (*Rubisco*), each with a corresponding isotopic fractionation denoted with ε. Loss fluxes were assumed to have no isotopic fractionation. In this proposed model, *f_1_* is defined as Φ_Loss1_/Φ_in_, and *f_2_* is defined as Φ_Loss2_/Φ_VCA_. See text for model assumptions. C) Experimental results plotted onto proposed box model outputs for *f_1_* = 0.1; colors and symbols are the same as Panel A. ε_P_ is defined as the difference in *δ*^13^C between C_ext_ and C_fixed_. See Supplemental Figure S10 for full results; only results for *f_1_* = 0.1 are shown.

We modified the traditional model (Figure 1B,C) by adding an additional isotopic fractionation so that ε_p_ can exceed ε_rubisco_. As discussed above, we hypothesize that this additional fractionation is due to a vectoral, powered CA like the NDH complex. In the modified model, we distinguish between carbon in the cytosol (C_int_) and carbon in the carboxysome (C_carb_), and allow a flux for carbon to be lost from the carboxysome (Φ_Loss2_, Figure 5B). Therefore, external C_i_ enters the cell (flux Φ_in_) where it can either leak out (flux Φ_Loss1_) or undergo active hydration (flux Φ_VCA_, where *VCA* denotes *V*ectoral *C*arbonic *A*nhydrase). Intracellular C_i_ can then enter the carboxysome, where it is either fixed (flux Φ_Rubisco_) or ultimately leaks out of the cell (flux Φ_Loss2_).

We made similar simplifying assumptions as the traditional box model: i) an infinite supply of external carbon, ii) no isotopic fractionation for carbon lost from the cell, iii) Φ_in_ has the isotopic fractionation associated with ε_Diffusion_, and iv) the system is at steady state. We did not add an explicit term for light energy used to power C_i_ uptake. Instead, the model included an energized CA (denoted VCA) and its associated isotopic fractionation as free parameters. In modeling each strain, we used the appropriate ε_Rubisco_ measurements (Table 1). We do not know the true value for ε_VCA_, but used a value of 30‰ similar to a recent model that explicitly invoked the NDH complex in Cyanobacteria (20). For comparison with the traditional model, we plotted Figure 5C with *f_1_* = 0.1 so that it could be represented in two dimensions; see Figure S10 for full model outputs. In this updated model, each value of ε_p_ corresponds to a set of feasible *f_1_* and *f_2_* values that fall along a line (Figure S10). Therefore, our model constrains but does not uniquely determine *f_1_* and *f_2_*, nor does it allow for estimation of external C_i_ levels because many pairs of *f_1_* and *f_2_* values can produce the same ε_p_. In addition, we focused only on C_i_ uptake as CO_2_ because we are interested in a model that could achieve large ε_P_ values (indicating ^13^C-depleted biomass) to account for at least an additional ~8‰ of fractionation in ε_p_ (maximum of ~25‰ in the high light condition) *greater* than εRubsico (~17‰) in ANC. HCO_3_^-^ uptake through bicarbonate pumps would not achieve this effect because it would shift all ε_p_ values to be 9‰ more negative (^13^C-enriched biomass, see Figure 1C).

With the addition of a powered, vectoral CA and an additional loss term, the model was able to rationalize our experimental data of ε_p_ > ε_Rubisco_ with leakage values compatible with cell growth (*f_2_* < 1, Figure 5C). In addition, it helped us understand why the high light condition gave such varied results between WT and ANC. Our model results implied that ANC lost more carbon than WT at the branch point before rubisco (Φ_Loss2_); i.e. even though carbon was present in the cell, it could not be fixed by the ancestral Form 1B rubisco, perhaps because of its lower *V_C_* (Table 1). Excess CO_2_ allowed rubisco’s KIE (ε_Rubisco_) to be expressed in ε_P_. These results implied that, in high light, the powered CA was delivering high amounts of CO_2_ to both the WT and ANC rubiscos. The faster WT rubisco was able to match this flux, which was reflected in its fast growth rate (Figure 3) and no change in ε_p_ vs. the reference condition (Figure 2). However, the slower ANC rubisco was not, which led to its slowest growth rate (Figure 2), and highest ε_p_ values across all conditions. Conditions where ε_p_ exceeded ε_Rubisco_ in ANC suggested that, in addition to Φ_Loss2_ being large (allowing ε_Rubisco_ to be expressed), Φ_Loss1_ was high as well, which allows ε_VCA_ to be expressed. That is, the slower ANC rubisco created a “backup” where leakage increased all along the carbon fixing pathway, and this effect was exaggerated at high light.

In addition, we fit our data to other models that are aware of active C_i_ uptake as part of general algal (21) or cyanobacterial (20, 59) CCMs (Figure S9). Cyanobacterial models that incorporated an explicitly one-way, “CA-like” enzyme (59) or the NDH complex specifically (20) were mostly able to rationalize our data as well. The poorest model fits for our data were when C_i_ uptake was mostly as HCO_3_^-^ (Figure S9). Overall, our model and theirs (20, 59) show that adding an additional carbon isotope fractionation step produces a model capable of rationalizing our data by enabling ε_p_ > ε_Rubisco_ with leakage values less than 1.

We also note that our use of the term “vectoral” CO_2_ hydration connotes a net flux that is dominantly in the direction of CO_2_ hydration, rather than implying that the flux of HCO_3_^-^ dehydration is zero. As such, there is likely some bidirectional activity of the NDH complex. It is difficult to experimentally measure the isotope effect associated with the hydration reaction (CO_2_ ⇒ HCO_3_^-^), but transition state theory and quantum chemical modeling (67, 68, 71) suggest that the value is large (roughly 25‰, see (28) for review). HCO_3_^-^ dehydration, and equilibration in general, would tend to reduce the isotopic fractionation (67). Our results here do not require a larger isotopic effect, however. Rather, a smaller value of ≈10‰ would have allowed us to rationalize our measurements, as we need only account for an additional ≈8‰ of fractionation in ε_P_ (maximum of ≈25‰) above ε_Rubsico_ (≈17‰) in ANC.

Overall, our measurements and analyses indicated that, in addition to rubisco, processes relevant to the CCM can play an important role in setting ε_p_ values. While our model is highly idealized and relies on a minimum set of fractionating processes associated with carbon fixation in Cyanobacteria, i.e. adding only one additional fractionation factor and one additional leakage point, the results demonstrated that a simple addition to the traditional model accounting for a known mode of energized CO_2_ uptake can explain our experimental results. Moreover, one useful implication of this model is that carbon isotope values may measure the efficiency of the CCM and carbon fixation in Cyanobacteria, in addition to ambient environmental CO_2_ concentrations.

### Consequences for understanding the evolution of carbon-fixing metabolisms

The goal of this study was to test if prevailing models of carbon fixation and isotopic fractionation held up in an ancestral analogue strain that may be more relevant to understanding the carbon cycle over geologic time. We did so by measuring the isotopic fractionation of a reconstructed ancestral rubisco both inside and outside a living Cyanobacterium. We emphasize that ANC is not a true ancestral Cyanobacteria; rather it is a chimeric construct—a modern strain saddled with a predicted Precambrian enzyme. This reconstructed ancestral rubisco is characterized by slower carboxylation kinetics (46) and a much lower ε_rubisco_ than the modern strain’s native enzyme (Table 1).

Recent studies in extant bacterial (20) and eukaryotic algae (for review see (28)) have already motivated updated models of C isotope fractionation in cells; these models address observations that: i) ε_p_ can exceed ε_Rubisco_ in certain conditions; ii) factors other than pCO_2_ can modulate ε_p_. We observed similar phenomena in our ANC strain, where ε_p_ exceeded ε_Rubisco_ in all conditions tested, and increased light intensity led to greater ε_p_ values. To date, such anomalous ε_p_ values have been observed during relatively slow growth; in (76) ε_p_ > ε_Rubisco_ occurred early in the growth curve as cells were acclimating to fresh culture media, in (28) ε_p_ > ε_Rubisco_ occurred during nitrogen and phosphorus limitation, and in this study ε_p_ > ε_Rubisco_ was observed in a mutant strain growing slowly while expressing a reconstructed ancestral rubisco. These observations indicated that growth physiology affects isotopic fractionation by photosynthetic algae and, in all cases, motivated a rethinking of the traditional box model (Figure 1B,C) to include more physiological detail relating to the presence of a CCM.

Here we observed ε_p_ > ε_Rubisco_ in all growth conditions for ANC, and especially in high light (Figure 2). As high light consistently slowed growth, induced chlorosis (yellowing of cultures, Figure S11) and increased ε_p_, we were motivated to consider the effects of light-related physiology on ε_p_. The yellowing of ANC cultures in high light was consistent with the well-described phycobilisome degradation pathway, which is typically induced in nutrient starvation conditions and taken to indicate that light levels exceeded the downstream capacity for CO_2_ fixation (58, 77). We interpreted these observations as indicating that the replacement of the native rubisco with a reconstructed ancestor decreased the cellular capacity for CO_2_ fixation, potentially due to the inferior carboxylation rate of the ancestral enzyme (Table 1).

Low CO_2_ fixation capacity would not, on its own, explain anomalously high ε_p_ values, however. An additional fractionating process is required to explain ε_p_ values in excess of ε_Rubisco_, which we assumed is due to light-coupled vectoral hydration of CO_2_, which has a large calculated isotope effect (67, 69–72). Cyanobacteria have been shown to take up CO_2_ independently of HCO_3_^-^ (61). In model Cyanobacteria, this activity is due to the Cup proteins (CupAS/B, also known as Chp proteins), which bind to the NDH complex of Cyanobacteria (66, 78). The NDH complex is involved in light energy capture via photosynthetic electron transport and cyclic electron flow around photosystem I (66) and, moreover, CO_2_ uptake is stimulated by light alone and abrogated by inhibitors of photochemical electron transport (73). Not only has CupA been shown to carry a key Zn^2+^ in a domain resembling a carbonic anhydrase (66), but the *cupA* gene is induced under low CO_2_ conditions (78). In order for CO_2_ uptake to drive the CCM and promote CO_2_ fixation, it would need to produce a high, non-equilibrium HCO_3_^-^ concentration in the cytoplasm (8, 10). We and others therefore assumed that the complex of NDH-1 and CupAS/B couples light energy to the one-way hydration of CO_2_ to HCO_3_^-^ at a carbonic anhydrase-like active site (66).

It is readily apparent from our experiments that ε_Rubisco_ does not set an upper bound on ε_p_, nor does it predict which strains will have larger ε_p_ values *in vivo* (Figure 2). This inference was only possible because we measured the isotope fractionation due to the ancestral rubisco (ε_Rubisco_) and compared it to ANC strain biomass (ε_p_), in contrast with the study of (57), which measured ε_p_ but not ε_Rubisco_. While our ANC ε_p_ values (≈18-24L) fell within the range of ε_P_ values derived from the carbon isotope record (42), they exceeded its measured ε_Rubisco_ (Figure 2). As such, the relative consistency of ANC ε_p_ values with extant ε_p_ values does not indicate that the traditional box model is applicable across geologic time as claimed in (57). Rather, a model including some additional fractionating process is required to explain our observation that ε_p_ > ε_Rubisco_ in ANC. Attention has been paid to outliers in the carbon isotope record where ε_p_ exceeds ε_Rubisco_ precisely because they violate the assumptions underlying the dominant model (Equation 1) used to interpret the record (28). In addition, ANC εRubsico (17.23 ± 0.61‰) is anomalously low; not only is it ≈8% less than WT εRubsico (25.18 ± 0.31‰) but it is among the lowest measured rubisco KIEs. However, only thirteen unique rubisco KIEs have been measured thus far (for recent review see (26)) while ≈300 distinct rubiscos have been kinetically characterized (7, 79), suggesting that measuring the isotopic effects of several well-chosen rubisco variants is worthwhile.

Our study in an ancestral analogue strain suggests that the carbon isotopic fractionations observed in both modern environments and in the geological record reflect not just the environmental abundance of CO_2_ and/or the rubisco present, but also the operation of C_i_ uptake processes, like the NDH complex, that operate as part of the CCM. Both these processes can lead to a highly variable range of carbon isotope fractionations. Our study supports the conclusion of prior studies that a carbon isotope model that engages more fully with photosynthetic physiology, like the CCM, is required to describe ε_p_ values and more accurately constrain environmental CO_2_ concentrations from environmental context (e.g. light and nutrient levels) and physiological parameters (e.g. ε_Rubisco_, photosynthetic capacity, growth rate). The model proposed here was written to mathematically validate a hypothesis – that ε_p_ can only exceed ε_Rubisco_ if another fractionating process was considered. In addition, it represents only a first step in this direction as it substantially simplifies the bacterial CCM (10); a similar statement applies to box models of Eukaryotic algae which also express complex CCMs (28, 80). Future work on carbon isotope fractionation by Cyanobacteria should grapple in more detail with photosynthetic physiology, including uptake of CO_2_ and HCO_3_^-^ by independent systems, integration of both light and dark reactions, and effects of nutrient limitation on growth rate. As mechanistic biochemical understanding of cyanobacterial C_i_ uptake improves (66), it may also become feasible to directly measure or better constrain the isotopic fractionation associated with these processes. Coupling such a model with experiments in natural and engineered organisms will help validate these models and improve our ability to understand environmental and evolutionary changes of the carbon cycle over Earth history.

In addition, this study and other recent work (42, 57) have raised a greater question for the Earth Sciences: what is uniformitarianism for biology? Earth scientists often apply uniformitarian assumptions – assuming that physical and chemical processes behave the same now as they did billions of years ago – in order to reason about the past. This approach is powerful, but these assumptions are challenged by biological processes that undergo substantial evolution on geologic timescales. Ongoing discoveries of novel metabolisms have supported some principles like ‘the principle of microbial infallibility’ – that microbes will always find a way to take advantage of available energy sources (for recent review see 81) – but it is not clear what principles apply to the details of metabolism. Take rubisco, for example – most extant autotrophs use rubisco to fix carbon, but rubisco sits within a variety of broader metabolisms (i.e. C3, C4, CAM in plants) that temper the effect of ε_Rubisco_ on ε_p_ (for recent review see (26)). We are far from having a clear answer to this question, but recent work at the interface of molecular biology and isotope geochemistry show that these ideas can be tested in the lab. Here and in other recent papers (42, 57, 82), we used synthetic biology to construct organisms with ancestral components so that specific aspects of ancient organisms can be isolated and tested. These “ancestral-like” organisms helped sharpen our understanding of the physiological and environmental factors determining growth (82) and isotopic fractionation (this work) in both ancient and modern autotrophs, and showed that models rigidly based on modern taxa are likely not universally applicable across geologic timescales.

Overall, carbon fixation was a fundamental challenge that autotrophs overcame early in the history of Earth’s biosphere (1). These early processes were recorded in some fashion in the carbon isotope record, but robust interpretation of this record must grapple with the fact that the carbon cycle is an amalgam of both environmental changes and evolutionary processes, mediated by physiology. We now have synthetic biological approaches that offer a way to probe these long timescale co-evolutionary problems by producing ancient process analogs of carbon fixation in the laboratory.

## Materials and Methods

### Ancestral enzyme reconstruction

Ancestral Rubisco enzyme sequences were previously reported and characterized by Shih et al. (2016) (46). Briefly, for both the large subunit (LSU) and small subunit (SSU) of Rubisco, encoded by *rbcL* and *rbcS* respectively, the most recent common ancestor (MRCA) for Form 1A (α), 1B (β), and 1A/B (α/ β) clades were predicted from independently derived phylogenetic trees for RbcL and RbcS containing a broad diversity of Form 1A and 1B Rubisco (>100 sequences). Maximum-likelihood algorithms were used to reconstruct the most probable ancestral sequence for each clade. Ancestral sequences were then expressed in *Escherichia coli* and purified, and enzyme kinetics were measured.

### ANC strain generation

The ‘ANC’ strain studied here was generated by replacing the native large and small Rubisco subunits (cbbL and cbbS respectively) of the parent strain (*Synechococcus elongatus* PCC 7942) with the reconstructed β ancestral cbbL and cbbS sequences. The NS2-KanR (‘WT’ strain) was generated by inserting a KanR cassette into neutral site 2 (NS2) (GenBank: U44761.1). This was done as a control for having the KanR in the neutral site. *Synechococcus elongatus* PCC 7942 were transformed from the WT strain using the approach of Golden and Sherman (1984) (83). Briefly, cultures were grown to OD750nm = 0.5. Cultures were centrifuged at 18,000 x *g* for 2 minutes. Pellets were washed with 100 mM CaCl2 and spun again at 18,000 x *g* for 2 minutes. Pellets were resuspended in BG-11 media followed by addition of plasmid and grown for 16 hours in the dark at 30°C. Transformants were then plated onto BG-11 + KAN100 agar plates and placed under 100 μE of light at 30°C. Single colonies were then genotyped by PCR amplification of the Rubisco locus followed by sequencing. Table S1 lists plasmids and primers used in this study.

### Growth conditions

For ambient CO_2_ growth, NS2-KanR and β Ancestral Rubisco-KanR strains were grown in quadruplicate in a photobioreactor (Photon Systems Instruments - MC 1000) at the University of California, Berkeley (UC Berkeley) for four biological replicates total. Cultures were grown in buffered BG-11 media with 50mM HEPES at pH 8. Cultures were inoculated at a starting OD720nm = 0.015 and cultivated at 120 μE, 30°C, and bubbled with ambient air. High CO_2_ growth was performed using the same conditions as ambient growth with the exception of placing the photobioreactor in a 5% CO_2_ chamber (Percival AR22L) and bubbling in air from the chamber. High light growth was performed using the ambient conditions above with the exception of using 500 μE for light intensity. Cells were harvested by centrifugation at 6000 x *g* for 20 minutes at 4°C. Decanted pellets were then flash frozen with liquid N2 and lyophilized overnight with the Millrock Technology Model BT85A freeze dryer. Doubling time was calculated by fitting the exponential phase of growth (*k*) using a Markov Chain Monte Carlo (MCMC) approach, using the generic model y = a*EXP(k*x)+b. Growth curves displayed in Figure 3 were smoothed with a rolling median (*n* = 12) to remove errant readings caused by bubbles advected in front of the detector. See Supplemental for more information.

### Carbon isotope analysis

Carbon isotope data is reported using delta notation (*δ*^13^C) in units of per mille (‰) where *δ*^13^C = [(^13^C/^12^C)_sa_/(^13^C/^12^C)_ref_-1]*1000, where the subscripts ‘sa’ and ‘ref’ denote sample and reference respectively. The reference used is the Vienna Pee Dee Belemnite (VPDB). *δ*^13^C values of cyanobacterial cells were measured on an EA-IRMS (Elemental Analyzer Isotope Ratio Mass Spectrometer; Costech Thermo Delta-V) at the California Institute of Technology (Caltech) in Pasadena, CA. Each biological replicate was run four times with two different isotope standards – urea (−27.8‰) and sucrose (−10.45‰). A suite of urea and sucrose standards were run at the beginning, middle, and end of run for sample bracketing and to assess drift throughout the run. An average *δ*^13^C and standard error were calculated and reported for each biological replicate (see Supplemental for more information). The *δ*^13^C of the starting CO_2_ gas was measured on the Thermo Mat 253 Ultra at Caltech; the CALT-2049C standard was used, which has a *δ*^13^C_VPOB_ value of −3.62‰. CO_2_ gas from high pCO_2_ experiments was sourced from a CO_2_ tank, while the CO_2_ gas in ambient pCO_2_ experiments was distilled from ambient lab air through cryogenic distillation at Caltech. ε_p_, the carbon isotope fractionation between CO_2_ gas and bulk cyanobacterial cells, was calculated as (α_CO2/bio_ - 1)*1000, where α_CO2/bio_ = ^13^R_CO2_/^13^R_bio_, where ^13^R is the ratio of ^13^C to ^12^C in the analyte. We note this in contrast to other isotope literature where ε_p_ is calculated as α_bio/CO2_ - 1)*1000, which would cause the positive values in this study to be negative. In this study, more positive ε_p_ values indicate more ^13^C-depleted; see Supplemental for more detail.

### Rubisco KIE assay

*Syn*6301 and β-MRCA Rubisco were purified according to previous methodologies (84, 85) at University of California, Davis and then shipped on dry ice to Caltech. Clarified lysate from a BL21 DE3 Star *E. coli* culture expressing Rubisco was subjected to ammonium sulfate precipitation, at the 30-40% cut for *Syn*6301 and at the 40-50% cut for β-MRCA, followed by anion exchange chromatography and size exclusion chromatography. We then used the substrate depletion method to measure the KIE of the *Syn*6301 and β-MRCA Rubiscos (ε_Rubisco_), as used previously in similar studies (47–50). Briefly, an assay mix of HCO_3_^-^, bovine carbonic anhydrase, rubisco, ribulose 1,5-bisphosphate (RuBP), MgCl_2_, bicine, and dithiothreitol (DTT) was prepared. As the reaction progressed to completion, aliquots of that assay mix were injected into pre-filled exetainers containing phosphoric acid that both stopped the reaction and converted all inorganic carbon species to gaseous CO_2_. The *δ*^13^C of these CO_2_ aliquots was then measured on a Delta-V Advantage with Gas Bench and Costech elemental analyzer at Caltech. Here, instead of RuBP being given in excess, CO_2_ was given in excess. In addition, instead of determining the fraction of CO_2_ (*f*) consumed independently to create a Rayleigh plot, we fit the curvature of the *δ*^13^C results to find *f* before converting to a Rayleigh plot to calculate ε_Rubisco_, similar to previous studies (48). See Supplemental for more information.

### Transmission Electron Microscopy (TEM) Imaging of Whole Cells

WT and ANC strains were grown in the reference condition as stated above (buffered BG-11 media, shaking at 250 rpm, with white cool fluorescent light at 120 μE, 30°C, and bubbled with ambient air (0.04% CO_2_ (v/v)). WT and ANC cells were collected at mid-log (40 and 80 h, respectively) at OD730=0.4 and pelleted by centrifugation (10,000 x *g* for 10 min). Pelleted cells were then resuspended in 1 mL of cold solution 2.5% Glutaraldehyde in 0.1M Sodium Cacodylate Buffer, pH 7.4 (Electron Microscopy Sciences) and stored in the fixative solution at 4°C until imaging. Sample preparation and sectioning were performed in the Electron Microscope Laboratory core facility at the University of California Berkeley. Briefly, samples were stabilized in 1% low melting-point agarose, cut into small cubes, and then washed at room temperature with 0.1 M sodium cacodylate buffer, pH 7. Samples were then mixed with 1% osmium tetroxide, 1.6% potassium ferricyanide and 0.1 M cacodylate buffer pH 7.2 for an hour in the dark with rotation. These were washed again with a cacodylate buffer pH 7.2, then DI water, and subjected to an 1 h incubation with uranyl acetate 0.5% solution. After a new wash with DI water, samples were dehydrated by an ascending series of acetone concentration (35%, 50%, 75%, 80%, 90%, 100%, 100%). Later, samples were progressively infiltrated in resin (Epon solution: Eponate 12, DDSA NMA and BDMA (Electron Microscopy Sciences) with rotation, followed by a final step at 60°C until polymerized. Thin sections (70 nm) were cut using a Reichert Ultracut E (Leica Microsystems) and collected on 100 mesh formvar coated copper grids. Sections were post-stained using 2% uranyl acetate in 70% methanol and followed with Reynold’s lead citrate. The sections were imaged using a FEI Tecnai 12 transmission electron microscope operated at 120 kV (FEI). Images were collected using UltraScan 1000 digital micrograph software (Gatan Inc).

## Acknowledgments

We thank Newton Nguyen for valuable guidance in the MCMC model used to calculate doubling times from growth curve data. We thank Victoria Orphan and Alex Sessions for access to lab space and analytical instruments, as well as lab managers Stephanie A. Connon, Fenfang Wu, and Nami Kitchen for assistance. This research was supported by the David and Lucille Packard Foundation (12540178), Simons Foundation (554187), NASA Exobiology (00010652), and the Schwartz-Reisman Collaborative Science Program (12520057). R.Z.W. was supported by a National Science Foundation Graduate Research Fellowship. Work in the lab of D.F.S. was supported by the US Department of Energy (DE-SC00016240). Work in the lab of P.M.S. was supported by a Society in Science–Branco Weiss fellowship from ETH Zürich and a Packard Fellowship from the David Lucile Packard Foundation. We thank Danielle Jorgens and Reena Zalpuri at the University of California Berkeley Electron Microscope Laboratory for advice and assistance in electron microscopy sample preparation and data collection.

