## Supplemental Text, Figures, and Tables for "Carbon isotope fractionation by an ancestral rubisco suggests biological proxies for CO_2_ through geologic time should be re-evaluated"

**Supplementary Information Text for Wang et al. (2022) "Carbon isotope fractionation by an ancestral rubisco enzyme *in vitro* and *in vivo* suggests biological proxies for CO2 through geologic time should be re-evaluated"**

**Table of Contents**

1. Calculating Doubling Times from Growth Curves
2. Carbon Isotope Measurements
    a. Delta Notation ($\delta^{13}C$)
    b. $CO_2$ Substrate
    c. Bulk Cyanobacterial Cells
    d. Error on reported $\delta^{13}C$ values
    e. Calculating $\varepsilon_P$ ($CO_2$/bio) and its error
3. Kinetic Isotope Effect of Rubisco
    a. Explanation of Rubisco Assay
    b. Assay Preparation and Execution
    c. Isotopic Measurement
    d. Calculating $\varepsilon_{rubisco}$ and its error
4. Cyanobacterial Box Models
    a. Traditional Box Model
    b. Proposed Box Model
    c. Fitting our data with other models
5. Emplacement of Rubisco into the Carboxysome
    a. Additional TEM Images
    b. Reconstructed ancestral rubisco residue analysis
    c. Spectroscopy

### 1. Calculating Doubling Times from Growth Curves

Growth constants, *k* ($hr^{-1}$), were fit using a custom Markov Chain Monte Carlo (MCMC) approach, written using MATLAB and Statistics Toolbox (vR2020b). Code can be found at https://github.com/reneezwang/ancestral-rubisco-cyano. We used this approach to limit human-based error on assessing when the exponential phase ended, and therefore left this as a free parameter for the MCMC.

To fit the exponential phase of growth, we created a model with five free parameters, and used an MCMC approach to find the best-fit values for each parameter. The model we fit follows an equation for exponential growth:

| | |
|---|---|
| $\boldsymbol{y = a * e^{k*x} + b}$ | (Eqn. S1) |

We then fit this model between a left-bound, *L*, and a right-bound, *R*, around the phase of exponential growth, so that only the exponential phase is fit. These bounds were left unconstrained so that they could be optimized by the MCMC. In total, we fit five parameters: 1) *a,* the pre-exponential factor (units of absorbance at 750 nm); 2) *k*, the growth constant (units of 1/hr); 3) *b*, the offset (units of absorbance at 750 nm); 4) *L*, the left bound (percentage of the length of data for each curve); 5) *R*, the right bound (percentage of the length of data for each curve). The MCMC found the best parameter by minimizing the $\chi^2$ value, and 100,000 to 1,000,000 steps were run for each curve. Fig. S1 shows the best-fit model for one growth curve. In black is the best-fit curve for the exponential phase of growth, with 1 sigma error shown in the black dotted lines. The best-fit left-bound, *L*, is shown in blue, with its 1 sigma error in blue dotted lines. The best-fit right-bound, *R*, is shown in red, with its 1 sigma error shown in red dotted lines. The corresponding parameter outputs are shown in Fig. S2. $\chi^2$ is quickly minimized, and a Gaussian curve is fit to each parameter to find the best fit value and 1 sigma error. The fitted *k* constants for each growth curve are summarized in Table S2. The doubling time was then calculated as ln(2)/*k*. Full growth curves are shown in Fig. S12.

### 2. Carbon Isotope Measurements

#### 2a. Delta Notation ($\delta^{13}C$)

Carbon isotope data were reported using delta notation ($\delta^{13}C$) in units of per mille (‰) where $\delta^{13}C = [(^{13}C/^{12}C)_{sa}/(^{13}C/^{12}C)_{ref}-1]*1000$, where the subscripts 'sa' and 'ref' denote sample and reference respectively. All values in this study were reported relative to the Vienna Pee Dee Belemnite (VPDB) reference.

#### 2b. $CO_2$ Substrate

Two different $CO_2$ substrates were used. For strains grown at ambient $CO_2$ concentrations (Reference Condition and High Light condition), ambient air was bubbled into the photobioreactor. Ambient air from the Savage lab at UC Berkeley was sampled into two 500 mL pre-evacuated glass bottles. Bottles were delivered by car to Caltech, where the contents were distilled on a vacuum line to separate and concentrate $CO_2$. Ambient air was cycled repeatedly as follows: 1) Sample was run over two traps filled with 3 mm diameter glass beads and immersed in liquid nitrogen in order to condensate $H_2O$ and $CO_2$; 2) $H_2O$ was then removed using a dry ice / ethanol slurry. For the High $CO_2$ condition, the $CO_2$ was sourced from a $CO_2$ tank so an aliquot was taken. The purified $CO_2$ from ambient air and the tank $CO_2$ were then both analyzed in triplicate on a Thermo MAT 253 at Caltech to measure its $\delta^{13}C$ value. The CALT-2049C standard, which has a $\delta^{13}C_{VPDB}$ value of -3.62‰, was used to correct measured lab values to the international Vienna Pee Dee Belemnite (VPDB) carbon isotope standard. Measured values can be found in Table S3.

#### 2c. Bulk Cyanobacterial Cells

As stated in the main text, cells were grown in a photobioreactor in the Savage Lab at UC Berkeley in each condition. Cells were then harvested by centrifugation at 6000 x *g* for 20 minutes at 4°C. Decanted pellets were then flash frozen with liquid $N_2$ and lyophilized overnight with the Millrock Technology freeze dryer (Model BT85A). Pelleted cells were then shipped on dry ice overnight to Caltech, where they were measured on a Delta-V Advantage with Gas Bench and Costech Elemental Analyzer (EA) at the California Institute of Technology. Each sample was measured 4 times on the EA. Each biological replicate was run four times with two different isotope standards – urea (-27.8‰) and sucrose (-10.45‰), so that carbon isotope values could be reported relative to VPDB. The uncertainties from correcting samples to this standard curve were smaller than the analytical replicate uncertainties, and so were ignored moving forward. A suite of urea and sucrose standards were run at the beginning, middle, and end of run for sample bracketing and to assess drift throughout the run. See Table S3 for finalized, drift-corrected values reported relative to VPDB.

#### 2d. Error on reported $\delta^{13}C$ values

For each condition, multiple biological replicates were grown (see Table S3 for number of replicates). Each biological replicate was then analyzed 4 times on the EA. The average of 4 technical replicates was taken to represent each biological replicate. The standard deviation (s.d.) was calculated from these values, and the standard error (s.e.) was calculated as s.d./($n$^0.5), where $n$ is the number of technical replicates.

#### 2e. Calculating $\varepsilon_P$ ($CO_2$/bio) and its error

$\varepsilon_P$, the vectorial isotopic fractionation between the inorganic carbon pool ($CO_2$) and bulk biomass (bio) can be calculated in one of two ways: i) From $CO_2$ to bulk biomass, or ii) From bulk biomass to $CO_2$. We calculated this value to be consistent with existing literature (i.e. (1)) in the fashion that follows. In this notation, a more positive $\varepsilon_P$ value means reaction products were more depleted in $^{13}C$.

We first calculated isotope fractionations as alpha values ($\alpha_{CO2/bio}$). $\alpha_{CO2/bio}$ is the relative difference between the $^{13}C/^{12}C$ ratios of two materials. This first requires converting $\delta^{13}C$ values to ratios of $^{13}C/^{12}C$ relative to the VPDB standard ($^{13}R_{VPDB}$; R denotes 'ratio'):

| | |
|---|---|
| $$^{13}R_{sa(VPDB)} = \left(\frac{\delta^{13}C_{sa(VPDB)}}{1000} + 1\right) \times \ ^{13}R_{std(VPDB)}$$ | (Eqn. S2) |

Where $^{13}R_{sa(VPDB)}$ or $^{13}R_{std(VPDB)}$ is the $^{13}R$ ratio of the sample or standard vs. the VPDB international scale, and $^{13}R_{std(VPDB)}$ = 0.01107828 as reported in Meija et al.(2) $\alpha_{CO2/bio}$ is then calculated as:

| | |
|---|---|
| $$\alpha_{CO_2/bio} = \frac{13R_{CO_2(VPDB)}}{13R_{cells(VPDB)}}$$ | (Eqn. S3) |

Then, alpha values were converted to $\varepsilon_{CO2/bio}$ values as:

| | |
|---|---|
| $$\varepsilon_{CO_2/bio} = (\alpha_{CO_2/bio} - 1) \times 1000$$ | (Eqn. S4) |

This $\varepsilon_{CO2/bio}$ value is the $\varepsilon_P$ value referred to in the text. A summary of all the calculated $\varepsilon_{CO2/bio}$ values are shown in Table S3, and the values used in the text are in Table S4.

**3. Kinetic Isotope Effect of Rubisco**

**3a. Rubisco Assay**

We used the substrate depletion method to measure the kinetic isotope effect catalyzed by Rubisco ($\varepsilon_{Rubisco}$), as used previously in similar studies (2–5). In this method, instead of directly measuring the difference in $\delta^{13}C$ of the reactants (i.e. 1 mol $CO_2$ and 1 mol RuBP) and products (i.e. 2 mol 3PGA), the $\delta^{13}C$ of one of the reactants ($CO_2$) is measured over time as the reaction goes to completion. One of the reactants is given in excess while the other is limited so that the $\delta^{13}C$ of the reactant pool eventually asymptotes to a final number as the reaction completes. In previous experiments, RuBP was given in excess while the $CO_2$ was limited. Finally, $\varepsilon_{Rubisco}$ is calculated by fitting the curvature of the results. This is often done by converting the results to a log-log plot, called a Rayleigh Plot, for ease of fitting. The curvature of this line, or its steepness in log-log space, is proportional to $\varepsilon_{Rubisco}$ - i.e. a rubisco with a large $\varepsilon_{Rubisco}$ will have a high degree of curvature and a larger slope in log-log space, and vice versa (Fig. S3).

The assay mix we used is based on previous similar studies. In this set-up, $CO_2$ is supplied in the form of $HCO_3^-$ which is converted to $CO_2$ by a carbonic anhydrase, typically derived from bovines. At equilibrium, this would cause the $CO_2$ pool to be lighter in $\delta^{13}C$ than the $HCO_3^-$ pool (Fig. S3). $CO_2$ and RuBP is then catalyzed by Rubisco to create 3PGA. Therefore, our reaction mixture contains carbonic anhydrase, rubisco, $HCO_3^-$, and RuBP to create the full reaction, and additional reagents such as: i) $MgCl_2$ to ensure the Rubisco active site is fully and correctly metallated, ii) bicine as a buffer, iii) dithiothreitol (DTT) to prevent oxidizing conditions that can inhibit rubisco activity and stimulate its degradation (6).

In our experiment, instead of limiting $CO_2$, we limited the other reactant, RuBP. In addition, *f* (the proportion of $CO_2$ remaining) must be known from an external measurement. Previous experiments have generally done so by taking a separate aliquot to measure the concentration of $CO_2$ directly (2, 5). In our experiment, we converted sampling time to *f* by fitting to the model y = a*EXP(-b*x)+c, based on the fact that the $\delta^{13}C$ of the reactant pool with increase during the reaction and then asymptote to a fixed value as the reaction ceases (i.e. no further carbon isotope discrimination can occur because Rubisco can no longer pull from the $CO_2$ pool as RuBP runs out). In essence, we are purely looking at the curvature of this line, similar to previous rubisco assays where the $\delta^{13}C$ of the reaction vessel headspace was monitored continually on a membrane inlet mass spectrometer (4) instead of traditional methods where discrete aliquots are taken (2). See Section 3d for further explanation.

The rubiscos used here were purified by the Shih lab according to previous methodologies (7, 8), and had their kinetics characterized previously (9). Briefly, as stated in the main text, clarified lysate from a BL21 DE3 Star *E. coli* culture expressing Rubisco (either the WT *Syn*6301 or β-MRCA) was subjected to ammonium sulfate precipitation, at the 30-40% cut for *Syn*6301 and at the 40-50% cut for β-MRCA, followed by anion exchange chromatography and size exclusion chromatography. The enzyme was then shipped on dry ice to Caltech, where the rubisco kinetic isotope effect (KIE) assay was performed.

**3b. Assay Preparation and Execution**

Prior to running the Rubisco KIE assay, the activity of bovine erythrocytes carbonic anhydrase (CA) ordered from Sigma Aldrich (C3934) was checked following the Sigma protocol titled "Enzymatic Assay of Carbonic Anhydrase for Wilbur-Anderson [W-A] Units (EC 4.2.1.1)" (10). We found a value of 3,368 W-A units / mg protein, which exceeded the product stated value of ≥2,000 W-A units / mg protein and proceeded to use this active CA enzyme prep in the rubisco KIE assay.

First, for the rubisco KIE assay, three external standards were prepared by weighing out Carrara marble standards (CIT_CM2013, $\delta^{13}C$ = +2.0 ± 0.1 (‰VPDB)) into three exetainer vials. Standards were then sealed within each tube, purged with He gas for 5 minutes, and then acidified by needle injection with concentrated phosphoric acid (42% v/v).

Next, three substrate exetainers were prepared. Three exetainer containers were purged with He gas for 5 minutes, and then injected with the substrate ($HCO_3^-$ dissolved in DI water). They were then acidified by needle injection with concentrated phosphoric acid (42% v/v) to convert $HCO_3^-$ to $CO_2$, and placed in a 70°C water bath for at least 20 minutes to help the reaction go to completion.

Then, 22 exetainer sampling vials were prepared for the WT and ANC rubisco assays (11 each). All tubes were first purged with He gas for 5 minutes, and then injected with ~1 mL of phosphoric acid. The phosphoric acid will both stop the reaction, and convert all C species into $CO_2$ for analysis.

Next, the reaction assay for each rubisco was prepared. First, a carbonic anhydrase (CA) stock solution was made by dissolving carbonic anhydrase from bovine erythrocytes from Sigma Aldrich (C3934) into DI water. Next, a ribulose 1,5-bisphosphate (RuBP) stock solution was made by dissolving D-Ribulose 1,5-bisphosphate sodium salt hydrate from Sigma Aldrich (R0878) in DI water. Then, one drop of concentrated hydrochloric acid (38% v/v) was added to 20 mL of autoclaved DI water while it was simultaneously stirred with a stir bar and purged with $N_2$ gas from a tube inserted into the solution. This was all done to remove any residual $HCO_3^-$ or $CO_2$ in the solution. The solution was purged for 10 minutes. Then, while $N_2$ gas was blown over the surface of the solution, reagents were added to create a final concentration of 100 mM bicine, 30 mM $MgCl_2$, and 1 mM dithiothreitol (DTT). $NaHCO_3$ from a pre-prepared stock solution was added, and pH was adjusted to 8.5 with NaOH and HCl. CA from the CA stock was added, and then either the WT or ANC rubisco was added to the solution. The solution was gently bubbled with $N_2$ gas for 10 minutes while rubisco 'activated.'

Next, the syringes used for each WT and ANC assay were cleaned with ethanol and water. We used two separate 25 mL gas-tight syringes with a sample-locking needle from the Hamilton Company for each Rubisco (Ref #86326, Model 1025 SL SYR).

Then, RuBP from the RuBP stock was added to each reaction assay, mixed through pipetting and swirling, and then quickly transferred to their respective gas-tight syringes. The first time point (t=0 min) was immediately taken after transfer. To sample, ~1 mL of the reaction assay was injected into the pre-prepared sampling exetainer vial so that the phosphoric acid in the vial would stop the reaction and convert all remaining $HCO_3^-$ to $CO_2$. Each assay was sampled 11 times over 429 minutes.

A control was run in a separate experiment, where all the assay components were mixed together with the exception of a rubisco enzyme. Its isotopic content was monitored through time. The $\delta^{13}C$ of the measured headspace did not change appreciably during this time period, with $\delta^{13}C = -0.42 \pm 0.03$ (‰VPDB) at t = 0 minutes, and $\delta^{13}C = -0.55 \pm 0.03$ (‰VPDB) at t = 277 minutes. The absolute values of these measurements reflect the $\delta^{13}C$ of the substrate used on that experimental day and cannot be related to the WT and ANC data shown here.

**3c. Isotopic Measurement**

The $\delta^{13}C$ of $CO_2$ in the headspace of each exetainer was measured on a Delta-V Advantage with Gas Bench and Costech elemental analyzer. Before measuring samples, two tests were performed to ensure the instrument was functioning normally: i) An 'on/off' test where an internal $CO_2$ standard was opened and closed to ensure instrument sensitivity and to establish a baseline intensity at a 'zero' $CO_2$ concentration, and ii) A linearity test where the concentration of $CO_2$ was increased linearly within the designated sensitivity range of the instrument to ensure that a linear increase in $CO_2$ concentration corresponds to a linear increase in electrical signal on the collector cups. We measured at three masses (44-46 amu). The instrument was also tuned to ensure that each mass was measured at the center of its mass peak.

The headspace of each sample and standard was measured ten times, with an internal $CO_2$ reference run before and after each suite of measurements. Each sample, with its ten measurement repetitions, were visually inspected to ensure the sample was being measured within the correct sensitivity range of the instrument (i.e. of similar intensity and pressure as the internal $CO_2$ reference). Peaks that did not meet this requirement were to be discarded, though no peaks were discarded for this particular assay. The 'raw' $\delta^{13}C$ values were then corrected relative to VPDB using the three standards run. The results of the WT and ANC rubisco assays can be seen in Table S5 and Fig. S4.

**3d. Calculating $\varepsilon_{Rubisco}$ and its error**

There are two sources of uncertainty that needed to be assessed in the Rayleigh plot; these sources are: 1) The spread in $\delta^{13}C$ or $^{13}R$ in the final few data points of the assay; 2) The $\delta^{13}C$ or $^{13}R$ of the *t* = 0 time point for both assays are different.

The spread in the last few points of our assay may be due to a variety of reasons, including: 1) Ambient $CO_2$ contaminating the exetainer containers as they are left out after the reaction; 2) Re-equilibration of the aqueous and gaseous inorganic carbon pools; 3) Instrument error. Since we expect the points to follow an exponential curve that eventually reaches an asymptote, we would therefore expect the points to fall along a straight line in a log-log plot. So, we converted our points from a linear space to a log-log space, systematically fitted lines through different sets of points in this space, and calculated the resulting error. The R value for these fits consistently decreased for the ANC assay after data point 10, and after data point 8 for the WT assay. Therefore, we proceeded using data points 1-10 for the ANC assay, and data points 1-8 for the WT assay.

The other issue in our data is that the $\delta^{13}C$ or $^{13}R$ of the *t* = 0 time point for both assays are different. We expect them to be similar, since both were given the same inorganic carbon pool to start with. However, the WT assay results are depleted in $\delta^{13}C$ relative to the substrate (Fig. S4) even though the remaining inorganic pool should become heavier as Rubisco preferentially uses $^{12}CO_2$ over $^{13}CO_2$ (so that our assay outputs, which sample this remaining pool, gets heavier). It appears that the initial substrate pool is contaminated with isotopically light $HCO_3^-$ or $CO_2$. Therefore, in order to treat both data sets equally, we did not use the $\delta^{13}C$ values of the $HCO_3^-$ substrate pool, as has been done previously to correct for the fractionation factor between $HCO_3^-$ or $CO_2$ (2) and instead derived the KIE from the curvature of the line (or slope in log-log space) as discussed in Section 3a and as done previously in (4). Therefore, we used *t* = 0 as the initial $R_0$ value for our starting substrate.

We converted time to *f*, the fraction of the inorganic C pool consumed. Since RuBP was the limiting substrate, we could calculate the moles of $CO_2$ consumed if we assume: i) A 1:1 ratio of RuBP to $CO_2$ was utilized by Rubisco, and ii) Full consumption of the RuBP pool. In this experiment, 5.47% of the initial $CO_2$ pool was consumed, or $f$ = 0.9543. We then assume that $f$ = 1 at $t$ = 0, and $f$ = 0.9543 at the upper bound of the fit.

A general model of y = a*EXP(-b*x)+c was used. The model y = a*EXP(-b*(x-d))+c was also tried, but no improvement to the fit occurred so we are only showing the best-fit model to the data. The model was fit three times using non-linear regression using MATLAB's *cftool* interface. The resulting fits and errors of those fits are shown in Table S6.

Time was then converted to $f$ using the equation:

| | |
|---|---|
| $$f = 1 - \left( \frac{R_i - R_1}{R_{upper} - R_1} \times (1 - F) \right)$$ | (Eqn. S5) |

Where $R_1$ is the first measured R value in each set of data, $R_{upper}$ is the fitted value 'c' from the general model y = a*EXP(-b*x)+c, and F = 0.9543, which is calculated from the amount of RuBP added to the assay.

Next, the values were converted to log space so that a Rayleigh plot could be made. We used the equation outlined in Guy et al. (2) to transform the R values:

| | |
|---|---|
| $$y = ln(R/R_0) \times 1000$$ | (Eqn. S6) |

Where $R_0$ is the first R value measured in each series. The $f$ values were transformed by taking the negative natural log. The values were then fit with the model y = m*x + b, and the coefficient 'm' was taken as $\varepsilon_{Rubisco}$. Results and the Rayleigh Plot are shown in Fig. S5 and Table S7. The average and standard deviation was calculated by averaging the three different 'm' coefficients that came from the three different fits. The standard error was calculated by dividing the standard deviation by the square root of $n$. The uncertainty in the 95% confidence interval was less than that of the standard deviation, and was therefore ignored for error propagation.

We found the WT (*Syn*6301 Rubisco) $\varepsilon_{Rubisco}$ value to be 25.18 ± 0.31‰ (avg ± s.e.), which was consistent with a previous measurement of a highly similar Form 1B Rubisco from *Synechococcus elongatus* 6301 by Guy et al. (1993), which found a value of 22.0‰. It is also consistent with other Form IB Rubiscos previously measured: i) 28.2 - 30.3‰ for *Spinacia oleracea* (2, 5, 11), and ii) 27.4‰ for *Nicotiana tabacum* (12). See (3) (13) (14) for excellent review and discussion of all currently known and measured Rubisco KIEs. We then found the ANC $\varepsilon_{Rubisco}$ value to be 17.23 ± 0.61‰ (avg ± s.e.).

### 4. Cyanobacterial Box Models

#### 4a. Traditional Box Model

The "traditional box model" described in the text is a simplified version of the model commonly used to relate $\varepsilon_P$ and $CO_2$ concentrations. We note that this is a dynamic area of research, and that many versions of this model topology exist with minor modifications. In this paper, we present a simplified version that is both accessible to those who are not isotope geochemists, and illustrates the primary relationship of interest – that as $\varepsilon_P$ increases, the external concentrations of $CO_2$ increase as well. The full history of this field cannot be covered here, but we give a brief summary to rationalize the traditional box model presented in the main text, and to give an introductory history to those who are not isotope geochemists.

The history of studying and modeling the carbon isotope fractionation of autotrophs (i.e. plants, algae, Cyanobacteria) tracks the birth and maturation of the field of isotope geochemistry. It began with the creation of the first modern, high-resolution mass spectrometers – the fundamental analytical tool that has enabled the field of modern isotope geochemistry – by the American physicist, Alfred O. Nier. Soon after Nier made the first isotopic measurements on a modern, high-resolution sector mass spectrometer (15) (16), his attention soon turned towards the isotopic composition of the natural world. In a seminal paper, Nier and Gulbransen noted the natural variation in carbon isotope ratios among igneous rocks, limestones, plants (in the form of anthracite coal and a modern pine tree), and "unclassified" samples like the air and a modern clam (17). Because of the advanced instrumentation, Nier and Gulbransen were able to improve upon previous studies by showing that these variations were not due to measurement error. Doing so, Nier and Gulbransen made the critical observation that plants tend to "concentrate the light isotope [$^{12}C$]" in comparison to air.

Later, more systematic measurements of plants and algae were carried out, which resulted in different theories of carbon isotope fractionation by autotrophs, notably a disagreement over if the $CO_2$ the plant was fixing was solely derived from the atmosphere, or potentially also from $CO_2$ originating from soils (either produced by microbial respiration of soil organic matter, or dissolved from limestone substrates) (18, 19).

The model that has come to dominate the field originated from a seminal study by Park and Epstein (20). They measured the carbon isotope ratios of tomato plants at varied $CO_2$ concentrations and light levels, as well as the carbon isotope fractionation associated with the rubisco enzyme itself. This key measurement allowed the construction of a "two step model" that could explain existing plant and algae data. Their model concluded that the first limiting step was "absorption of the $CO_2$ from the atmosphere by the leaf," and the second was the "enzymatic conversion of 'dissolved $CO_2$' in the cytoplasm to carbohydrates." They proposed that the isotopic fractionations of rubisco and diffusion are not additive *in vivo* – instead, they proposed that the net isotopic fractionation *in vivo* (bulk biomass carbon isotope composition) reflects the process by which photosynthesis is being limited. Therefore, if photosynthesis were exclusively limited by diffusion, the bulk biomass fractionation ($\varepsilon_P$) would reflect only the diffusive process ($\varepsilon_P = \varepsilon_{Diffusion}$). And if diffusion did not limit photosynthesis, the bulk fractionation would instead reflect rubisco ($\varepsilon_P = \varepsilon_{Rubisco}$). Finally, they noted that though "[t]he model presented here is necessarily in its simplest form and as such, does not define in detail mechanisms responsible for the $C^{13}/C^{12}$ fractionation in $CO_2$ fixation," they were still able to explain both their experimental & literature data based on the "two step model."

Farquhar et al. 1982 built upon this key assumption from Park and Epstein – that the isotopic fractionations of diffusion and rubisco are *not* additive *in vivo* (21). While Farquhar et al. acknowledged that other factors in addition to diffusion and rubisco may affect isotopic fractionation during photosynthesis, their goal was to reconcile most of the differences between observed and expected fractionations, and to create a model so that "measurements of gas exchange physiology and isotopic fractionation" could be made. Importantly, they derived a relationship between the ratio of the partial pressures of atmospheric vs. intercellular $CO_2$ and the bulk carbon isotope fractionation. This allowed their model to be used to predict changes in plant water use efficiency in photosynthesis & carbon isotope fractionation, since both are tied to opening / closing the stomata (where $CO_2$ diffuses into the plant). It also allows the $CO_2$ concentration at the site of rubisco to be estimated from the measured isotopic fractionation.

Interestingly, it was debated in the literature at the time if the isotopic fractionation of each rubisco enzyme *itself* varied. This would be a way to explain variations in $\varepsilon_p$. Farquhar et al. (1982) does note that Whelan et al. (1973) (22) found that rubisco fractionation changes with temperature, but that Christeller et al. (1976) (23) does not. Farquhar et al. state that "it is likely that much of the variation presently evident in the literature reflects experimental uncertainties rather than intrinsic variations in the capacity of the enzyme to fractionate carbon isotopes" (21). Therefore, the current isotope models built upon & after Farquhar (1982) all make the assumption that the isotopic fractionation of rubisco is constant.

This "two-step model," largely based on Park and Epstein, can be derived for plants as follows. In this model architecture, Fig S6A and Main Text Figure 1B, carbon can be: i) external to the cell ($C_{external}$ or $C_{ext}$), ii) inside the cell ($C_{internal}$ or $C_{int}$), or iii) fixed by the cell into biomass ($C_{fixed}$). Carbon that enters the cell but does not get fixed by Rubisco is assumed to eventually be lost by the cell, and return to the external carbon pool ($C_{lost}$).

We used the classic Hayes isotope flux model system to evaluate our results (24). In this approach, each flux has its own isotopic fractionation ($\varepsilon$), as well as carbon isotope composition ($\delta$). For the carbon pools, this $\delta$ refers to the isotopic composition of the pool. For the fluxes, $\delta$ refers to the instantaneous isotopic composition of that flux (see (24) for a detailed review). We also made a set of simplifying assumptions: i) The system is at steady state, ii) The external carbon pool is infinitely large compared to the cell (i.e. its carbon isotope composition does not change). We first defined the isotopic relationships for each flux in our system:

| | |
|---|---|
| $\delta_{in} = \delta C_{ext} + \epsilon_{in}$ | (Eqn. S7) |

| | |
|---|---|
| $\delta_{loss} = \delta C_{int} + \epsilon_{loss}$ | (Eqn. S8) |

| | |
|---|---|
| $\delta_{Rubisco} = \delta C_{int} + \epsilon_{Rubisco}$ | (Eqn. S9) |

We will also define $\varepsilon_P$ as the difference in $\delta^{13}C$ of the external vs. fixed carbon pools, i.e.:

| | |
|---|---|
| $\varepsilon_P = \delta C_{ext} - \delta C_{fixed}$ | (Eqn. S10) |

Most of these models are solved with the assumption of steady state, which we will assume as well. We can then define the mass balance relationships with φ denoting fluxes; $\varphi_{in}$ is the flux of carbon into the cell, $\varphi_{loss}$ is carbon loss from the cell, and $\varphi_{Rubisco}$ is carbon that is fixed by rubisco:

| | |
|---|---|
| $\boldsymbol{\varphi_{in} = \varphi_{loss} + \varphi_{Rubisco}}$ | (Eqn. S11) |

The traditional model assumes that the amount of carbon entering the cell is inversely proportional to a concentration gradient of $pCO_2$ inside vs. outside of the cell, or that $\Phi_{out}/\Phi_{in}$ = $[C_{int}]/[C_{ext}]$. So, we can then define a loss fraction:

| | |
|---|---|
| $\boldsymbol{f_{\blacksquare} = \frac{\varphi_{loss}}{\varphi_{in}}}$ | (Eqn. S12) |

The isotopic relationships and mass balance equations were combined to create an isotope mass balance equations:

| | |
|---|---|
| $\boldsymbol{\varphi_{in}\delta_{in} = \varphi_{loss}\delta_{loss} + \varphi_{Rubisco}\delta_{Rubisco}}$ | (Eqn. S13) |

These sets of equations can be solved symbolically to arrive at the solution:

| | |
|---|---|
| $\boldsymbol{\varepsilon_P = (1 - f)(\varepsilon_{in}) + f(\varepsilon_{Rubisco})}$ | (Eqn. S14) |

This solution is plotted as the green line in Figure S7A and referred to as the 'plant-based' model.

Much work was done after this to adapt the plant-based model to algae. The main modification done was to account for active $C_i$ uptake in the form of $HCO_3^-$ or $CO_2$ (25). The Sharkey & Berry (1985) model is very similar to the plant-based model in that: 1) A linear relationship exists between $\varepsilon_P$ and inorganic carbon ($C_i$) leakage out of the cell (defined as $F_3/F_1$ in Sharkey & Berry (1985), and defined as $f=\Phi_{loss}/\Phi_{in}$ in this study); 2) $\varepsilon_P$ cannot exceed $\varepsilon_{Rubisco}$. We have plotted the plant-based model vs. the Sharkey & Berry (1985) in Figure S7A below – the slope of both lines is set by $\varepsilon_{Rubisco}$, and the models only differ by their y-intercept. This is because active $C_i$ uptake was a known part of the CCM, and Sharkey & Berry (1985) took this into account by assuming all $C_i$ entering was $HCO_3^-$ (flux $F_1$ in Sharkey & Berry (1985) Figure 4). This causes the $C_i$ pool inside the cell to be ≈8‰ enriched in $^{13}C$, which causes the y-intercept to be more negative (in this community's framework, a positive $\varepsilon_P$ value means $^{13}C$-depletion while a more negative $\varepsilon_P$ value means $^{13}C$-enrichment). This is plotted as the blue line in Figure S7A and referred to as the Sharkey & Berry model.

Popp et al. 1989 (26) and Laws et al. 1995 (27) also made key contributions by extending this plant-based model to algae. Popp et al. worked to account for issues related to growth physiology— specifically growth rate, cell shape and size— to adapt the C3 plant model to unicellular algae. Interestingly, they found cyanobacterial $\varepsilon_p$ to be roughly constant independent of environmental $pCO_2$ and growth rate. (This is in contrast to contemporaneous studies in Cyanobacteria at the time that *did* find cyanobacterial $\varepsilon_p$ varies with $pCO_2$ (Erez et al. 1998).) They hypothesized that this invariance stems from the large surface area to volume ratio (SA/V) of Cyanobacteria, which was taken to imply much faster passive $CO_2$ uptake (scaling with SA) than fixation (scaling with V). Because cyanobacterial $\varepsilon_p$ was constant ≈17‰ and less than known cyanobacterial $\varepsilon_{Rubisco}$ values, additional fractionating factors were not needed to explain $\varepsilon_p$, even though some active transport processes related to light were known in Cyanobacteria at the time (28–30). They note, "Although results of our experiments suggest that $CO_2$(aq) does not cross the plasmalemma by passive diffusion alone, but rather is supplemented by an active transport mechanism, the inescapable conclusion is that $\varepsilon_p$ nonetheless varies as a linear function of growth rate, [$CO_2$(aq)] and the cellular-carbon-to-surface-area ratio under most natural conditions." In other words, the simple linear relationship between $pCO_2$ and $\varepsilon_p$ in C3 plants appeared to hold up in algae and Cyanobacteria as well.

Many versions of this traditional model exist. Eichner et al. (2015) (31) presents a nice version of the traditional model that is stated in their study as a generalization of the Sharkey & Berry (1985) model (Equation 15 in Eichner et al.) that we are citing and presenting as the "traditional" model in Figure 1 in the main text. It relates the plant-based model and the Sharkey & Berry model by introducing the term $a_{cyt}$, which varies the proportion of $CO_2$ vs. $HCO_3^-$ in total $C_i$ uptake (Figure 1; Figure S7B). We use the Eichner et al. $\varepsilon_{db}$ value of -9‰ instead of Sharkey & Berry (1985) $\varepsilon_{db}$ value of -7.9‰. Essentially, in the Eichner et al. version of the Sharkey & Berry model, when $a_{cyt}$ = 0, all $C_i$ uptake is $CO_2$ and you get the plant-based model. When $a_{cyt}$ = 1, all $C_i$ uptake is $HCO_3^-$ and you get the Sharkey & Berry (1985) model solution. We note that all of these models have the key limitation that $\varepsilon_P$ cannot exceed $\varepsilon_{Rubisco}$.

The final step was to extend this model to environments both modern and ancient. Francois et al. 1993 and Rau et al. 1989 both found, from measuring the carbon isotope composition of particulate organic matter (POM) or phytoplankton from ocean surface waters, that concentrations of dissolved $CO_2$ were correlated with $\varepsilon_P$ values (32, 33). These studies were notable because they showed the prior model calibrated in the lab could potentially be extended to the field, and that a model calibrated in plants even seemed to hold in algae. In addition, Hayes formalized the above model into an isotope flux model that is the dominant mathematical form used to model autotrophic carbon isotope fractionation today (34). Hayes also increased the model's detail by predicting the isotopic composition of specific metabolic intermediates, and by extending this model to new metabolic systems like eukaryotic lipid biosynthesis. He also noted that values of $\varepsilon_P$ derived from the carbon isotope record may "provide information about the nature of the primary producer organisms and their environment" like "$CO_2$ paleobarometry."

Popp et al. (26) had previously determined the isotopic compositions of sedimentary porphyrins, but did not estimate paleo-$CO_2$ levels because their model had empirically fit parameters (i.e. "*b*") that they could not determine for ancient environments and materials. This *b* term is an empirically fit slope that "quantifies the rate at which $\varepsilon_P$ decreases as concentrations of $CO_2$ become smaller," and is related to $\varepsilon_P$ by the relationship $\varepsilon_P = \varepsilon_f - b/C_e$, where $\varepsilon_f$ is the isotopic fractionation of all carbon-fixation reactions active in the cell but mainly rubisco, and $C_e$ is the concentration of dissolved $CO_2$ (35). The term *b* effectively sets how quickly $\varepsilon_P$ approaches the limit of $\varepsilon_{Rubisco}$ (Figure S7D). Freeman and Hayes (36) subsequently showed that, indeed, they could calculate ancient $CO_2$ levels up to 100 Ma after calibrating the empirical "*b*" value from Popp et al. (26). Much work continues today empirically calibrating this model so that it can be applied to geologic time (37). Common values used for *b* are on the order of magnitude ≈100‰

kg μM$^{-1}$ (38). Overall, both the C Isotope Record Model and the Traditional Model have a limit where $\varepsilon_P$ cannot exceed $\varepsilon_{Rubisco}$ (Figure S7C,D).

We refer to this as the "C Isotope Record Model" in the main text. It is derived from work based on model organisms in the lab (i.e. the Traditional Box Model shown in Figure 1 and S7C) because the parameter *b* is derived from bench-top lab experiments.

**4b. Proposed Box Model**

Our proposed model incorporated two more boxes and an additional isotope fractionation factor (Figure S6B). Therefore, the four main reservoirs are: i) Carbon that is external to the cyanobacterial cell ($C_{ext}$); ii) Carbon inside the cell ($C_{int}$); iii) Carbon in the carboxysome ($C_{carb}$); and iv) Carbon that is fixed into biomass ($C_{fixed}$). The three isotope effects are: i) Diffusion into the cell ($\varepsilon_{in}$); ii) Fractionation by the a powered carbonic anhydrase which catalyzes the unidirectional hydration of $CO_2$ to $HCO_3^-$ ($\varepsilon_{PCA}$); iii) Fractionation by rubisco during carbon fixation ($\varepsilon_{Rubisco}$). For $\varepsilon_{in}$ a value of 1(‰VPDB) was used based on the diffusion of $CO_2$ in water (39). For $\varepsilon_{PCA}$, a wide range of values exist in the literature based on both lab experiments and *ab initio* calculations using transition state theory, but they range from 13-39(‰VPDB) as shown in Wilkes and Pearson (2019), which offers an excellent discussion on the topic, and we direct the reader to that paper for further reading (40). We used a value of 30 (‰VPDB) based on a previous study by Eichner et al. (2015), who used this value to model C isotope fractionation by the NDH-1$_4$ complex in their model organism *Trichodesmium erythraeum* IMS101 (31). For $\varepsilon_{Rubisco}$, two different values were used for either the WT or ANC strain, based on *in vivo* measurements done for this paper. $\varepsilon_{Rubisco}$ = 17.23 ± 0.61 (‰VPDB) for ANC, and $\varepsilon_{Rubisco}$ = 25.18 ± 0.31 (‰VPDB) for WT. These values were derived as detailed in Section S3.

Finally, we then permitted two pathways for loss in our system. The first flux is for C that diffuses into the cell, but then exits the cell and does not continue into the carboxysome ($\varphi_{loss1}$). The second flux is for C that enters the carboxysome but is not fixed by Rubisco, and then exits the cell ($\varphi_{loss2}$).

We again use the classic Hayes (2001) isotope model to model our system (24). This model assumes that the system is at steady state. We defined the isotopic relationships for each box and flux in our system:

| | |
|---|---|
| $\delta_{in} = \delta C_{ext} + \varepsilon_{in}$ | (Eqn. S15) |
| $\delta_{loss1} = \delta C_{int} + \varepsilon_{loss1}$ | (Eqn. S16) |
| $\delta_{PCA} = \delta C_{int} + \varepsilon_{PCA}$ | (Eqn. S17) |
| $\delta_{loss2} = \delta C_{carb} + \varepsilon_{loss2}$ | (Eqn. S18) |
| $\delta_{Rubisco} = \delta C_{carb} + \varepsilon_{Rubisco}$ | (Eqn. S19) |

We then defined the bulk cyanobacterial fractionation, $\varepsilon_P$, as:

| | |
|---|---|
| $\varepsilon_P = \delta C_{ext} - \delta C_{fixed}$ | (Eqn. S20) |

Since there is only one path for the last flux into the $C_{fixed}$ box,

| | |
|---|---|
| $\delta_{Rubisco} = \delta C_{fixed}$ | (Eqn. S21) |

So:

| | |
|---|---|
| $\varepsilon_P = \delta C_{ext} - \delta_{Rubisco}$ | (Eqn. S22) |

As in the prior section, we then defined the mass balance relationships with φ denoting fluxes; $\varphi_{in}$ is the flux of carbon into the cell, $\varphi_{loss1}$ and $\varphi_{loss2}$ are carbon loss from the cell, and $\varphi_{PCA}$ is carbon that goes through a hypothetical powered carbonic anhydrase (PCA), and $\varphi_{Rubisco}$ is carbon that is fixed by rubisco:

| | |
|---|---|
| $\varphi_{in} = \varphi_{loss1} + \varphi_{PCA}$ | (Eqn. S23) |

| | |
|---|---|
| $\varphi_{PCA} = \varphi_{loss2} + \varphi_{Rubisco}$ | (Eqn. S24) |

We also defined the two loss fractions, $f_1$ and $f_2$:

| | |
|---|---|
| $f_1 = \frac{\varphi_{loss1}}{\varphi_{in}}$ | (Eqn. S25) |

| | |
|---|---|
| $f_1 = \frac{\varphi_{loss2}}{\varphi_{PCA}}$ | (Eqn. S26) |

The isotope relationships and mass balance equations were combined to create the isotope mass balance equations:

| | |
|---|---|
| $\varphi_{in}\delta_{in} = \varphi_{loss1}\delta_{loss1} + \varphi_{PCA}\delta_{PCA}$ | (Eqn. S27) |

| | |
|---|---|
| $\varphi_{PCA}\delta_{PCA} = \varphi_{loss2}\delta_{loss2} + \varphi_{Rubisco}\delta_{Rubisco}$ | (Eqn. S28) |

These set of equations was solved symbolically to arrive at the solution:

| | |
|---|---|
| $\varepsilon_P = \varepsilon_{loss2} - \varepsilon_{loss1} - \varepsilon_{in} + f_1(\varepsilon_{PCA} - \varepsilon_{loss1}) + f_2(\varepsilon_{Rubisco} - \varepsilon_{loss2})$ | (Eqn. S29) |

Equation S29 was solved analytically as described in the section above, except two different *f* vectors were inputted: $f_1$ and $f_2$. See GitHub for code for plotting and solving at https://github.com/reneezwang/ancestral-rubisco-cyano. Full model results are shown in Fig. S10. Figure 4C in the main text shows solutions for *f*, = 0.1, which is denoted as shown in Fig. S10.

In addition, we focused only on $C_i$ uptake as $CO_2$ because we are interested in a model that could achieve more negative $\varepsilon_p$ values ($^{13}$C-depleted biomass), and $HCO_3^-$ uptake (i.e. through bicarbonate pumps like BicA, SbtA, or BCT1 (41) ) would not help us because it would shift all εp values to be ≈8‰ more positive ($^{13}$C-enriched biomass).

Model outputs are discussed in the main text, and we note that our model is *highly* idealized – we tried to modify the traditional model as little as possible to explain our data, which was to achieve $\varepsilon_p$ > $\varepsilon_{Rubisco}$ with physiologic consequences that make sense. We wanted to demonstrate with our simple, proposed model that just slight modifications to the traditional model can start to harmonize our experimental results with model outputs. This may allow for future modeling avenues that can continue to augment our understanding of carbon isotope fractionation within bacterial autotrophs.

**4c. Fitting our data with other models**

We fit our data with three other algal carbon isotope models to see if they could rationalize our results – the Sharkey & Berry model (25), the Erez et al. model (42), and the Eichner et al. model (31).

Sharkey and Berry measured the carbon isotope fractionation of plants and eukaryotic algae, *Chlamydomonas reinhardtii*, grown at varied $pCO_2$ conditions and derived a model for carbon isotope fractionation by algae that accounts for the algal CCM (see Figure 4 and Equation 2 in (25); re-written in Figure S9A). This model accounted for the CCM by taking into account active $C_i$ uptake, and it assumed that all $C_i$ entering the cell was in the form of $HCO_3^-$ and that all $C_i$ lost from the cell is as $CO_2$. They defined the loss of $C_i$ from the cell as the ratio of two relative fluxes, $F_3$ and $F_1$, which are plotted on the x-axis in Figure S9A. We plotted our measured $\varepsilon_P$ values (colored circles) using this model and got $C_i$ leakage values ($F_3/F_1$) that exceeded 1 for all ANC data, and for WT High $CO_2$ data. Leakage values greater than 1 imply that the cell is not fixing any carbon, which is incompatible with our growth curve data (i.e. ANC grew in all conditions, and was therefore fixing carbon).

Erez et al. (42) grew batch cultures of the cyanobacterium *Synechococcus* sp. PCC7942 (the same parent strain used in this study) bubbled with ambient lab air and found $\varepsilon_p$ values up to 33‰, greater than $\varepsilon_{Rubisco}$ values known at the time (28 or 22‰). This result is in contrast to Popp et al. who found using *Synechococcus* sp. CCMP838 that cyanobacterial $\varepsilon_p$ values do not vary with growth rate or changing $CO_2$ concentrations or exceed known cyanobacterial $\varepsilon_{Rubisco}$ values (1). Therefore, Erez et al. also need an additional fractionation factor to explain their data, and presented a model in their Equation 4 that modifies the Sharkey and Berry (25) model by adding a separate compartment for the carboxysome. They also invoke a "CA-like" enzyme that catalyzes the one-way hydration of $CO_2$, which both scavenges $CO_2$ lost from the carboxysome and introduces an additional isotopic fractionation factor since the isotopic fractionation of this reaction is thought to be large (they test 12‰ and 15‰ as potential values). We are interested in the relationship between $\varepsilon_p$ and $C_i$ lost, which is the difference in $C_i$ lost ($F_3$ in their Figure 6) versus $C_i$ uptaken ($F_1$ in their Figure 6). So, we rearranged Equation 4 in Erez et al. using Equation 1 in Erez et al. to derive the equation:

$$\epsilon_P = X\epsilon_{equil} + \epsilon(\frac{F_3}{F_1}) \text{ (Equation S30)}$$

Where *X* is the fraction of $CO_2$ to total $C_i$ uptake (*X*=1 is all $CO_2$, *X*=0 is all $HCO_3^-$). The modification to the Sharkey and Berry model is the addition of this term, *X*. The Erez model was able to largely rationalize ANC $\varepsilon_p$ data ($F_3/F_1<1$), but only if all $C_i$ uptake is $CO_2$ (*X*=1), and it gives extremely high leakage values for the high light condition (0.99 and 0.90) (Figure S9B). In addition, if *X*=1 for WT, then implausible negative values for leakage ($F_3/F_1$) are calculated for three of the four reference condition replicates (-0.04, -0.02, -0.03) and all the high light replicates (-0.003, -0.01). Overall, the Erez model implies that $C_i$ leakage is overall higher for ANC vs. WT. In addition, their model only fits ANC $\varepsilon_p$ values in the unlikely scenario that all $C_i$ uptake by ANC is $CO_2$.

Eichner et al. (31) grew the cyanobacterium *Trichodesmium erythraeum* IMS101 with varied nitrogen sources at varied $pCO_2$ concentrations and compared leakage estimates derived from $\varepsilon_p$ with an independent, non-isotopic method of membrane inlet mass spectrometry (MIMS). We note that they used a diazotrophic cyanobacterium while we did not. Similar to our study, they found that isotopic leakage estimates derived using the Sharkey and Berry model (25) regularly exceeded 1, while MIMS estimates gave more reasonable values (see their Figure 3). Similar to (42), they needed an additional isotopic fractionation factor, so they modified the Sharkey and Berry model by adding a compartment for the carboxysome and called upon the NDH complex specifically, which results in Equation 14 and 15 of their paper, re-written as:

$$\epsilon_P = a_{cyt}\epsilon_{equil} + L_{cyt}(a_{carb}\epsilon_{cyt} + L_{carb}\epsilon_{Rubisco})$$ (Equation S31)

Where the fractionational contribution of $HCO_3^-$ to total $C_i$ uptake into the cytosol or carboxysome is $a_{cyt}$ or $a_{carb}$ respectively ($a$=1 is all $HCO_3^-$); the relative proportion of $C_i$ leaking out of versus entering the cytosol or carboxysome is $L_{cyt}$ or $L_{carb}$ respectively; $\varepsilon_{cyt}$ is the isotopic fractionation of the NDH-$1_4$ complex.

Because of the independent MIMS method used in (31), they were able to independently constrain parameters that we could not (i.e. $a_{cyt}$). Therefore, we use the values they found most likely to explain their results, which is Scenario 5 in their Table 2 ($a_{carb}$=1, $a_{cyt}$=0.8, $\varepsilon_{cyt}$=30) and varied $L_{carb}$ from 0 to 1. They note that although an $\varepsilon_{cyt}$ value less than 30‰ could explain their data if the other parameters were varied, "In a scenario assuming an upper estimate for $\varepsilon_{cyt}$ of +30‰ (scenario 5, Table 2), which is within the range of fractionation measured in other enzymes such as RubisCO, our MIMS-measured data can be reproduced even for the high $pCO_2$ treatment."

Using the Eichner model, we are able to rationalize all of our WT and ANC data, though only for $L_{carb}$ > ≈0.2 for ANC (Figure S9C). This is consistent with their results, which suggests an $L_{carb}$ value of 0.9. However, we note that they invoke the NDH complex for internal $C_i$ recycling, to convert $CO_2$ lost from the carboxysome back to $HCO_3^-$ for re-entry in the carboxysome. We invoke the NDH complex for light-powered $CO_2$ uptake. Regardless, both the Eichner model and ours are able to rationalize $\varepsilon_p$ data by calling an additional fractionation factor that allows $\varepsilon_{Rubisco}$ to exceed $\varepsilon_p$ (i.e. derived leakage values are less than 1).

For all models, we solved analytically for values of $\varepsilon_P$, given the experimentally measured values of $\varepsilon_{Rubisco}$, and inputting values of $f$ ranging from 0 to 1. We then plotted our experimental $\varepsilon_P$ values onto the model output, which gave us a value of $f$. After doing so, we noticed—perhaps unsurprisingly—that ANC $\varepsilon_P$ values could not be plotted onto the model outputs, as described in the main text and shown in Figure 4A. The code for plotting and solving can be found on GitHub at https://github.com/reneezwang/ancestral-rubisco-cyano.

**5. Emplacement of Rubisco into the Carboxysome**

ANC strain growth at ambient $pCO_2$ supports the conclusion that the CCM is functioning properly, as it is well established that CCM deletions / mutations prevent cyanobacterial growth at ambient $CO_2$ (see (43, 44) for review). In addition, the carboxylation rate ($V_C$) for the ancestral rubisco is roughly half that of the extant rubisco (4.72±0.14 $s^{-1}$ vs. 9.78±0.48 $s^{-1}$ respectively), so the CCM has to be working for it to be able to grow at ambient. Consistent with these past results, a recent paper utilizing an ancestral analogue strain, Hurley et al. (2021) (45), deleted the CCM and found that their strain does not grow at $CO_2$ levels of 1, 18, and 30x PAL (present atmospheric levels) but was able to grow at 36 and 107x PAL at pH 7.3-8.1. Therefore, ANC strain growth at ambient $pCO_2$ supports the conclusion that the CCM is functioning correctly.

In addition, Shih et al. (2016) (9) shows rubisco emplacement using fluorescence microscopy with tags for RbcL (rubisco large subunit) and CcmN (carboxysomal subunit) in their Figure 8. Though that strain expresses both the extant and ancestral rubisco RbcS and RbcL sequence, there is no rubisco fluorescence seen external of the carboxysome.

For even further due diligence, we wanted to ensure that the ancestral rubisco emplaces properly into the carboxysome, and that doing the full swap of the extant for the ancestral rubisco sequence does not have any unintended physiologic effects on other aspects of the CCM. We performed two additional analyses: 1) Transmission electron microscopy (TEM) imaging of carboxysomes, and 2) Searching for residues shown to be required for successful rubisco emplacement into the carboxysome.

**5a. Additional TEM Images**

Additional TEM images are shown in Figure S13. Briefly, WT and ANC cells were grown in the reference condition (ambient $pCO_2$, normal light flux) and harvested at mid-log. Cells were sectioned and prepared for TEM imaging with the help of University of California Berkeley Electron Microscopy Lab. See Methods for full sample preparation and sectioning details.

**5b. Reconstructed ancestral rubisco residue analysis**

In Cyanobacteria, rubisco and carbonic anhydrase (CA) proteins are packed tightly within the carboxysome as liquid condensates (43). Successful formation of $\beta$-carboxysomes involves aggregation of rubisco by the scaffolding protein CcmM. It has recently been shown in *Synechococcus elongatus* PCC 7942, the same model organism used in this study, that cysteine residues in the small subunit-like (SSUL) module of CcmM is key for this process, and that disulfide bond formation in the SSUL is required for carboxysome formation *in vivo* (46).

In addition, Wang et al. show that SSUL interacts with rubisco at two interfaces, Interface I and Interface II. The structural features of these two interfaces are shown in Figure 4c and 4d of their manuscript with the contact residues specified (46). We performed an alignment of the WT and reconstructed ancestral rubisco sequence using Clustal Omega (47) (48) and looked for these residues. We found that eight of the ten residues were conserved for Interface I, and all residues were conserved in the ancestral sequence for Interface II (Tables S8 and S9, and Figure S14). This, in addition to the TEM imaging and growth of ANC at ambient $pCO_2$, gives us confidence that substituting the extant rubisco sequence with the reconstructed ancestral sequence does not affect carboxysome function, and that the ancestral rubisco emplaces within the carboxysome.

**5c. Spectroscopy**

In order to compare the pigment composition displayed by wild type versus ANC mutant, we performed room temperature absorbance spectra measurement between 400-800 nm for cultures with similar density (OD730=0.4). WT and ANC strains were grown in the reference condition as stated above (buffered BG-11 media, shaking at 250 rpm, with white cool fluorescent light at 120 µE, 30°C, and bubbled with ambient air (0.04% $CO_2$ (v/v)). WT and ANC cells were collected at mid-log (40 and 80 h, respectively) at OD730=0.4. Samples with OD730 = 0.4 (NanoDrop OneC Microvolume UV-Vis, Thermo Scientific) were prepared as described for electron microscopy (see Methods in main text) and absorbance spectra were measured with a UV–Vis Scanning Spectrophotometer (UV-2101PC, Shimadzu, Japan) in the range of 400-800 nm. Data was normalized to emission at 800 nm.

Results can be seen in Figure S15. Absorbance measurements confirmed the chlorosis phenotype observed for the ANC strain. The WT and ANC strains were normalized to the same optical density at 800nm, however, the ANC strain demonstrated lower relative absorbance values at 620nm where phycocyanin, the major pigment of phycobilisomes is known to absorb.

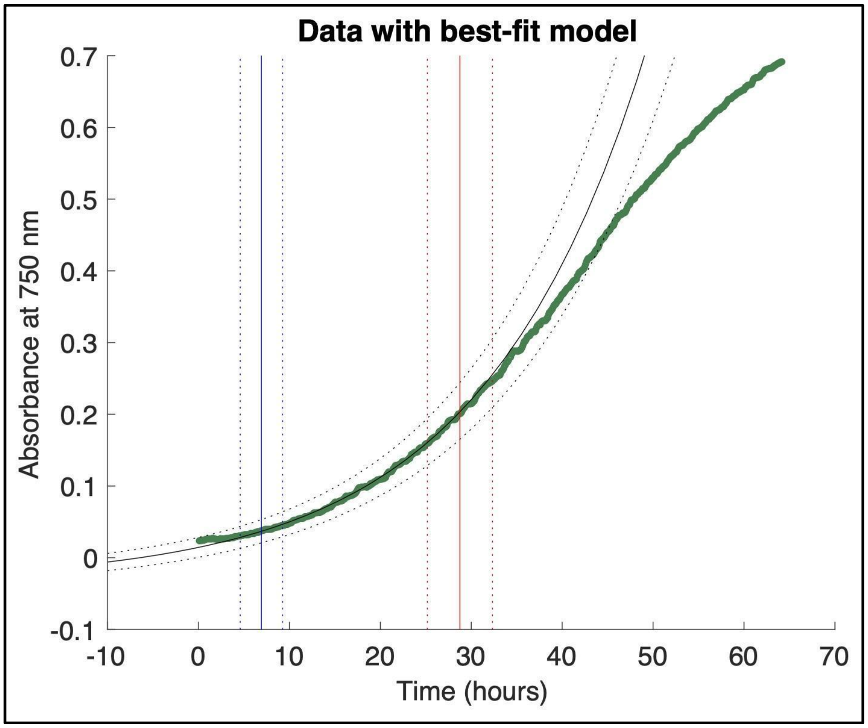

**Figure S1.** Best-fit model for calculating growth constant, *k*, for one growth curve. Black solid and dotted lines indicate best fit for the exponential section of the growth curve. Blue solid and dotted lines indicate best fit left bound. Red solid and dotted lines indicate best fit right bound. Analyses were performed using MATLAB and Statistics Toolbox (vR2020b).

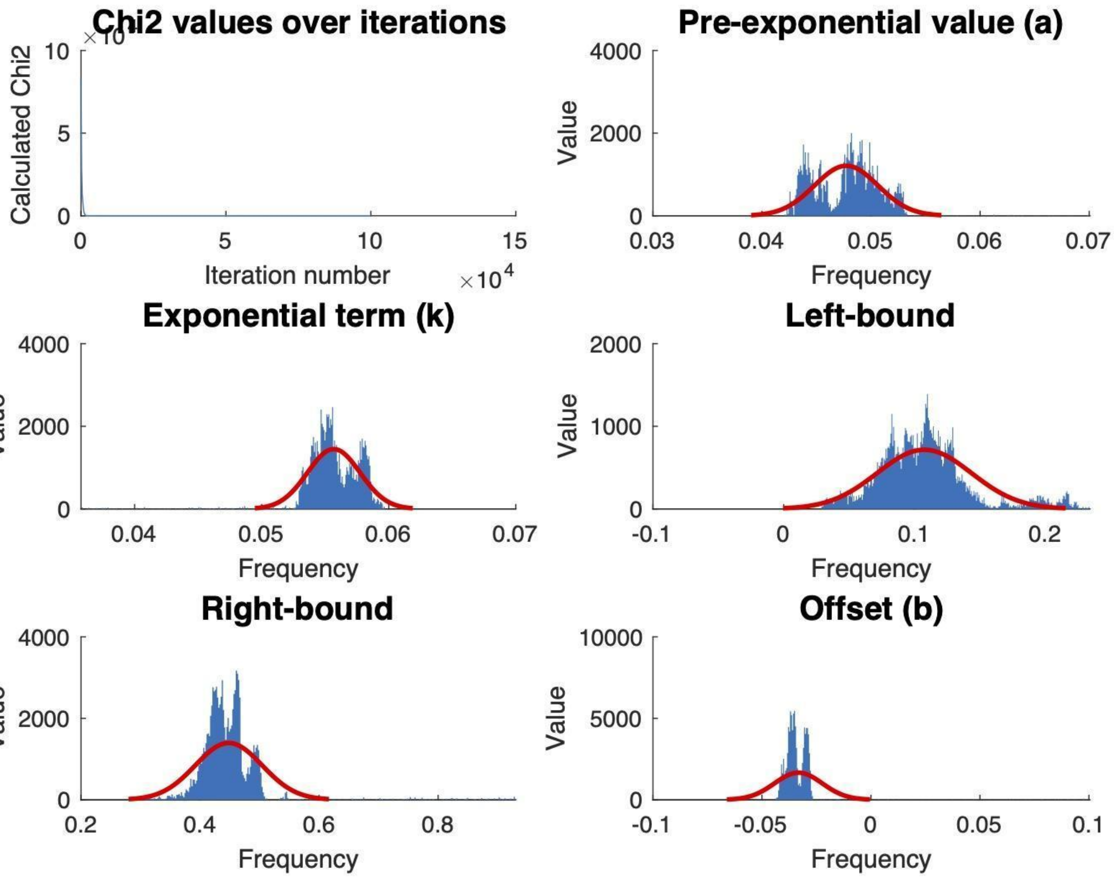

**Figure S2.** Outputs for parameters used in MCMC to calculate the growth constant, *k*. A histogram of each output is shown in blue, and a probability density function fit to the data is shown in red. Analyses were performed using MATLAB and Statistics Toolbox (vR2020b).

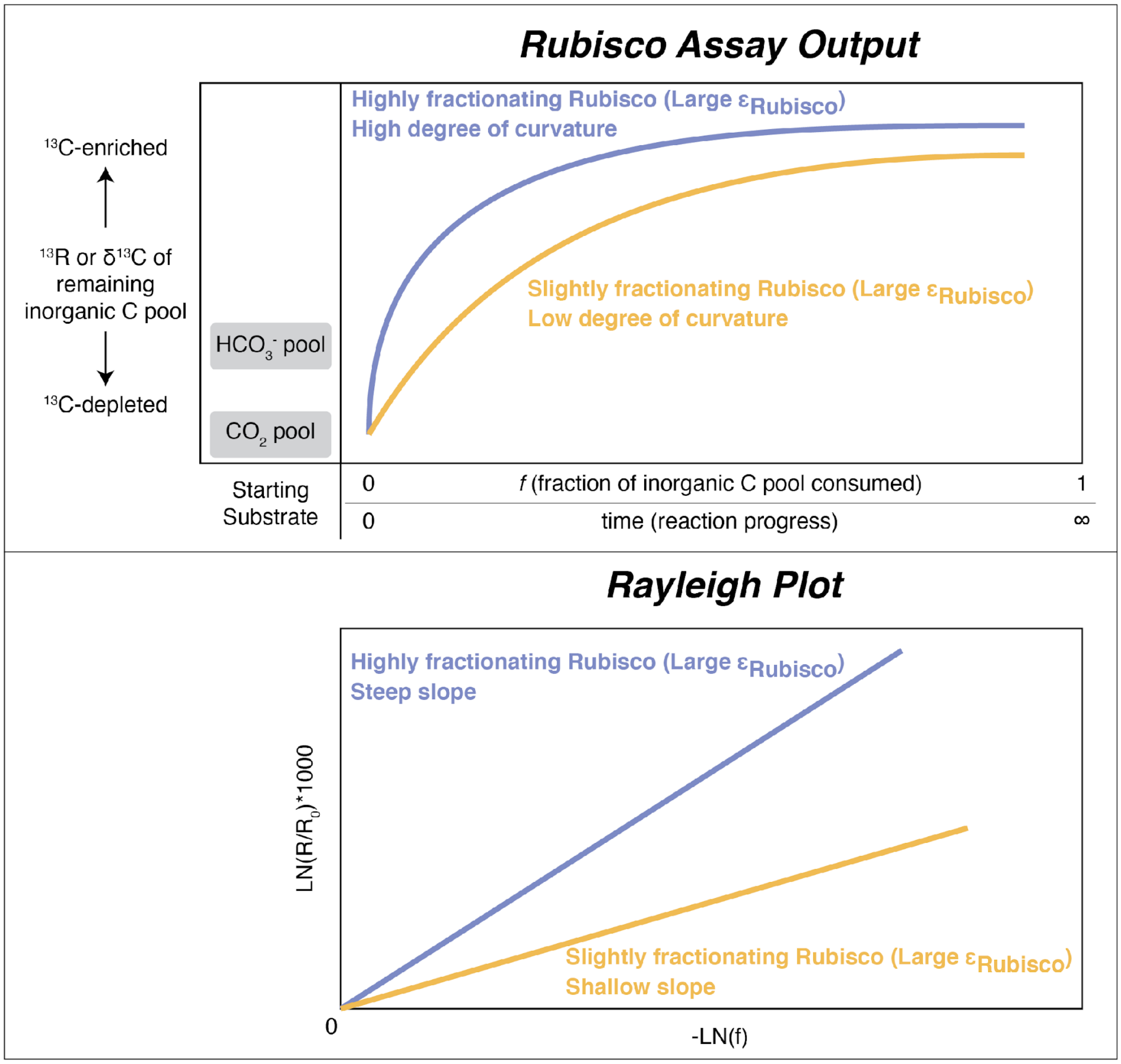

**Figure S3.** Cartoon showing expected results of Rubisco assay for strongly vs. slightly fractionating Rubisco. Top panel shows measured outputs of $\delta^{13}C$ or $^{13}R$ values vs. reaction progress or fraction of inorganic C pool consumed (*f*). Bottom panel shows the log-log version of that plot, which is called a Rayleigh plot. $R/R_0$ is the $^{13}R$ ratio of the sample at a given time point vs. the initial $^{13}R$ ratio of the starting inorganic C pool.

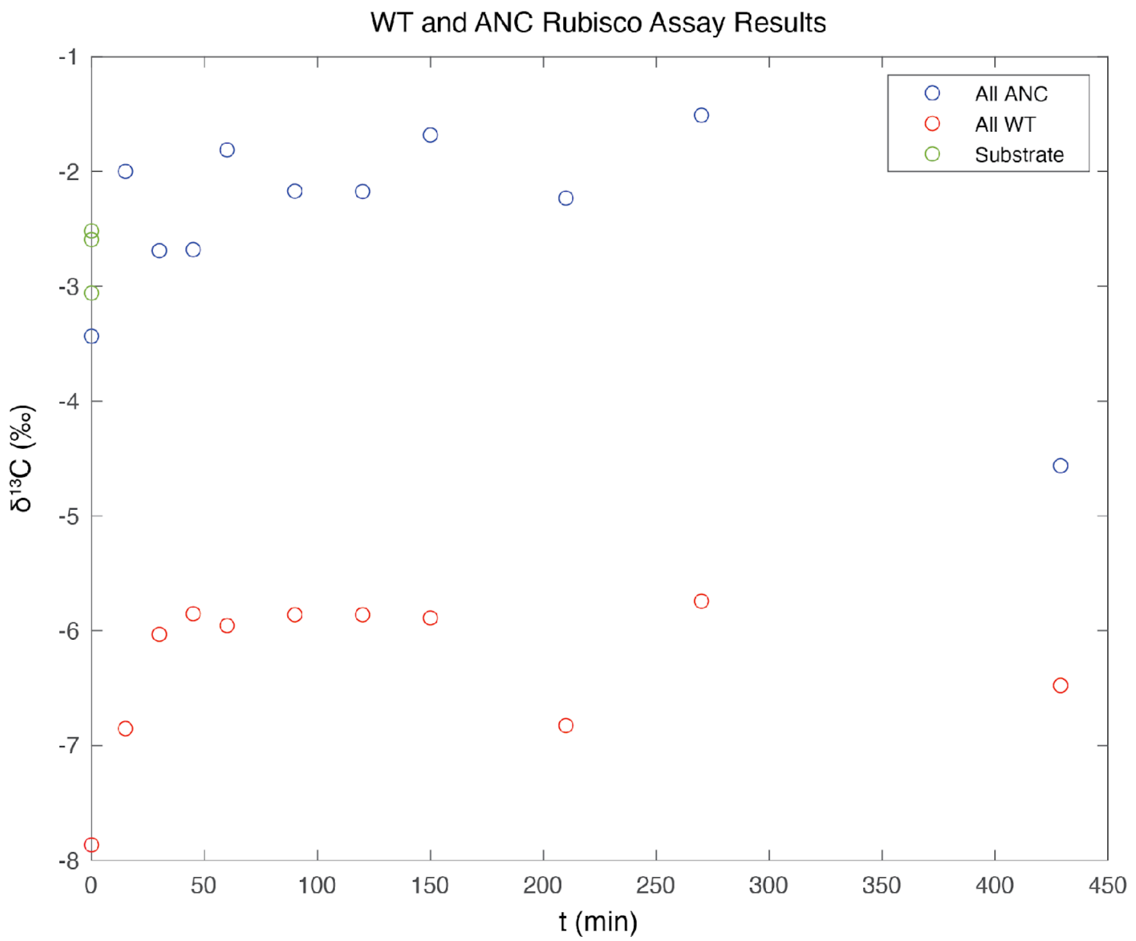

**Figure S4.** Results of WT (in red) and ANC (in blue) Rubisco assays, shown as $\delta^{13}C$ vs. time (minutes). Substrate (in green) indicates acidified $HCO_3^-$ substrate; it is plotted at t=0 for ease of viewing. Analyses were performed using MATLAB and Statistics Toolbox (vR2020b).

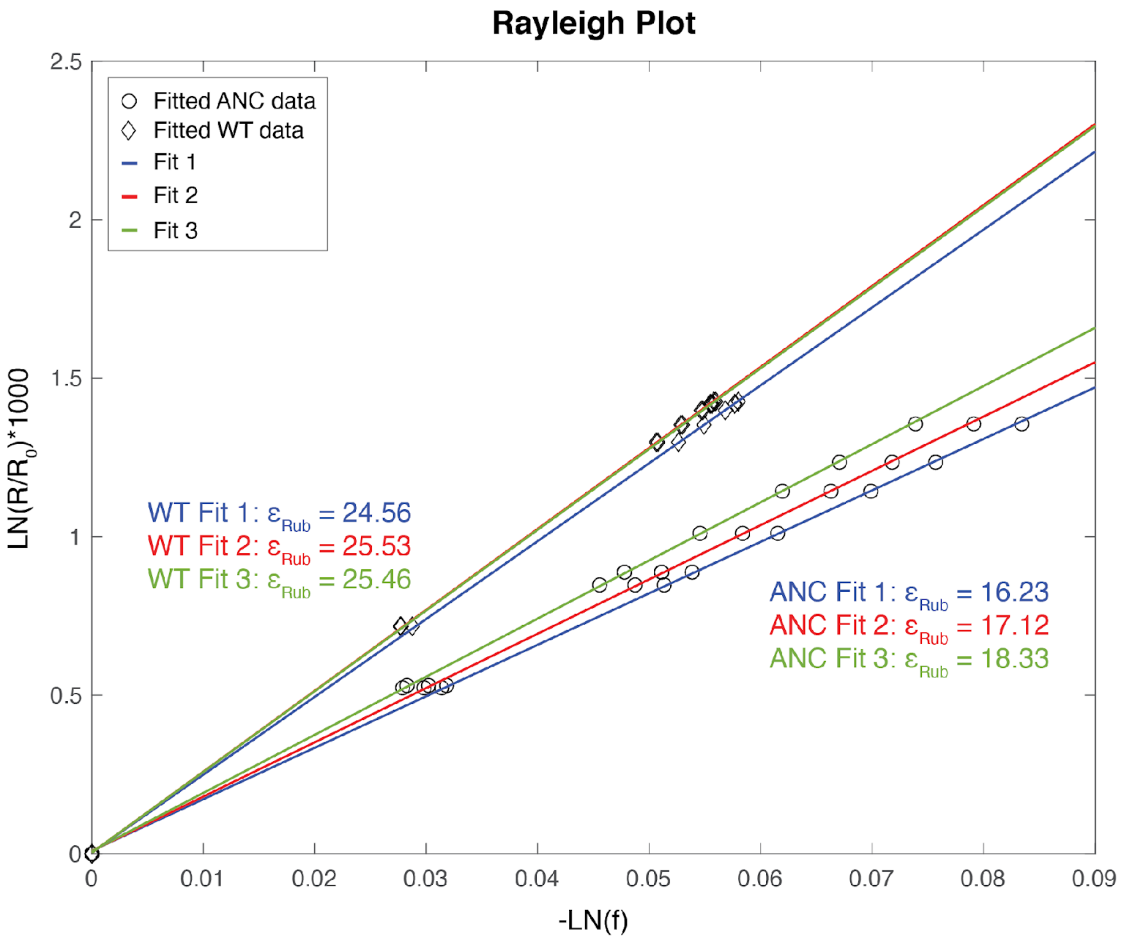

**Figure S5.** Rayleigh plot for WT and ANC Rubisco assays. Analyses were performed using MATLAB and Statistics Toolbox (vR2020b).

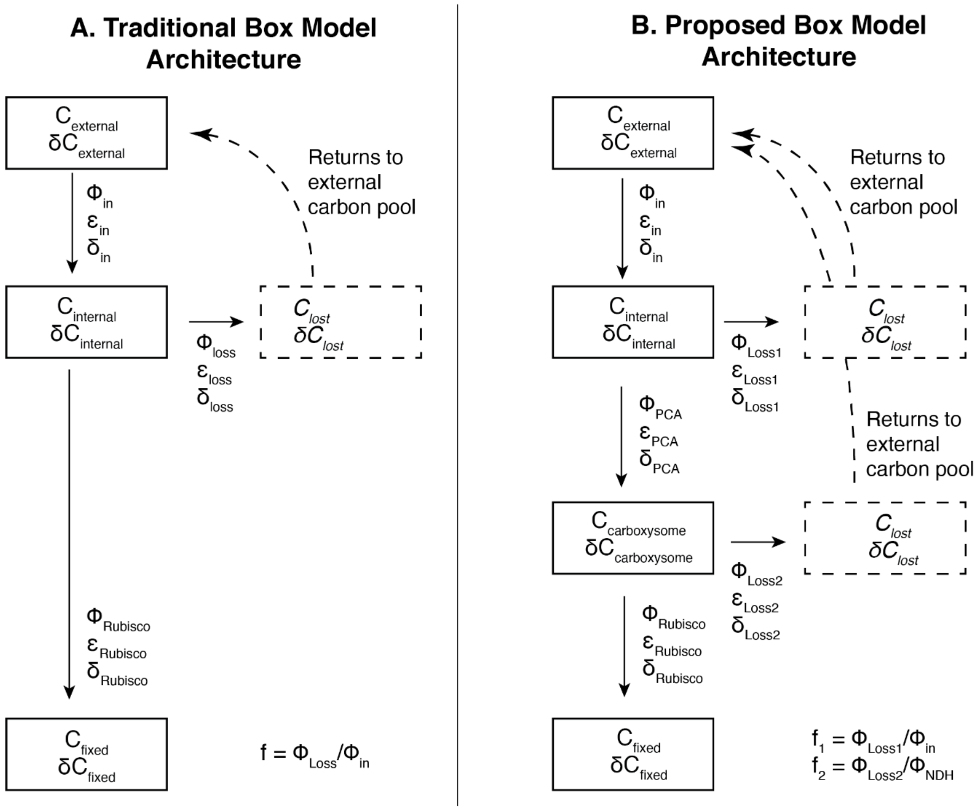

**Figure S6:** Model architecture for: A) The traditional box model, and B) Our proposed box model. PCA = Powered Carbonic Anhydrase.

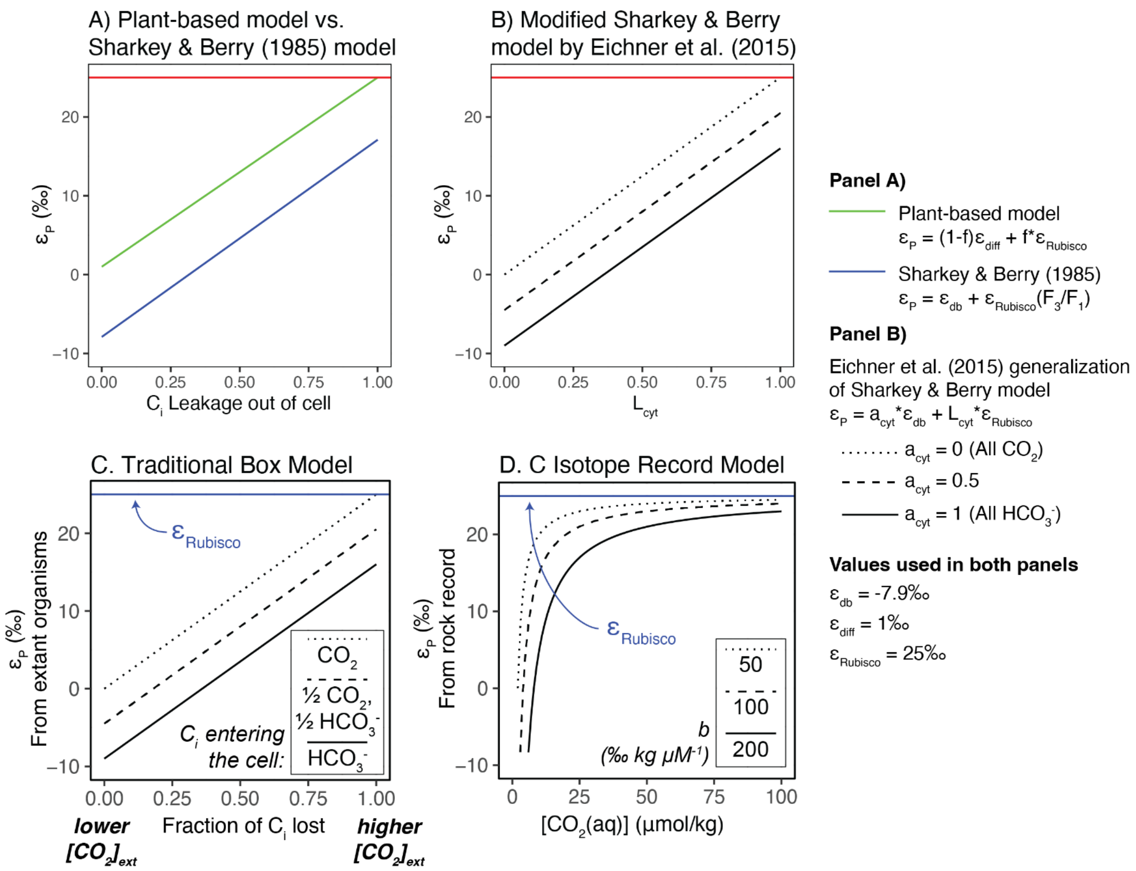

**Figure S7: Variations on the "Traditional Model" and the "C Isotope Record Model."** A) The plant-based model we derived in the supplemental is shown in green, while the model proposed by Sharkey and Berry (1985) for algae generally is shown in blue. Both models have the slope of $\varepsilon_{Rubisco}$ (25‰ is used as an example here). They are offset by the equilibrium fractionation of $CO_2 \leftrightarrow HCO_3^-$, where $HCO_3^-$ is $^{13}C$-enriched relative to $CO_2$ (in this field's reference frame, a more negative isotopic value). The equations for each model are given in the right panel of the figure; for simplicity, we label the x-axis as "$C_i$ leakage out of the cell" because it is named differently in each model (*f* in our derivation; $F_3/F_1$ for Sharkey and Berry (25) Figure 4). B) The Eichner et al. (2015) generalization of the Sharkey and Berry model. Eichner et al. (31) derives a two-compartment cyanobacterial model that can be generalized to the Sharkey and Berry model, as well as the plant-based model. The equation is shown in the right-most panel, and results in a line with a slope of $\varepsilon_{Rubisco}$ and a y-intercept set by the term $a_{cyt}$*$\varepsilon_{db}$ to show if the total $C_i$ uptake is primarily $CO_2$ ($a_{cyt}$=0) or primarily $HCO_3^-$ ($a_{cyt}$=1). When $a_{cyt}$=0, you effectively get the plant-based model in Panel A), and when $a_{cyt}$=1, you get the Sharkey & Berry model in Panel A). Other values used are $\varepsilon_{Rubisco}$ (fractionation of the enzyme rubisco) = 25‰, $\varepsilon_{diff}$ (fractionation of $CO_2$ diffusing into the cell) = 1‰. For $\varepsilon_{db}$, the fractionation of the $CO_2 \leftrightarrow HCO_3^-$ equilibrium, Sharkey & Berry (1985) used a value of -7.9 while Eichner et al. (2015) uses a value of -9. All analyses were performed using R Statistical Software (v4.1.0; R Core Team 2021) and figures were produced using the ggplot2 package (v3.3.6; Wickham, 2016). C) The traditional box model as shown in the main text. $\varepsilon_P$ values are measured from extant organisms in the lab. D) The C Isotope record model. $\varepsilon_P$ values are derived from the rock record. Both C) and D) have an upper limit where $\varepsilon_P = \varepsilon_{Rubisco}$.

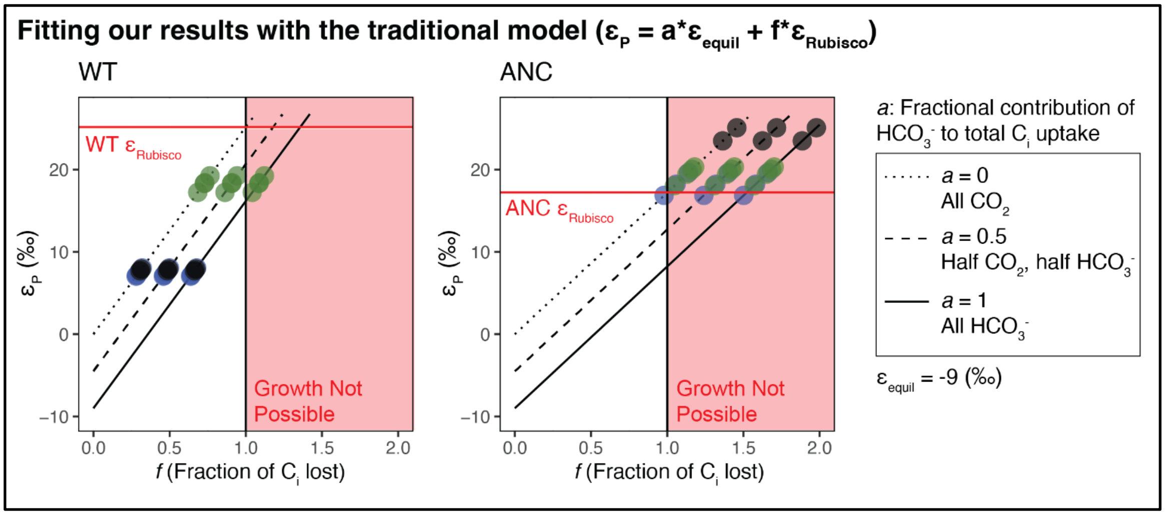

**Figure S8: ANC data cannot be rationalized with the traditional model.** Measured $\varepsilon_P$ values for each strain (circles) were fit with the traditional model at varying $C_i$ uptake compositions (lines). Blue circles indicate reference condition (ambient $pCO_2$ (0.05% (v/v)), standard light flux (120 µE)); Green circles indicate high $CO_2$ condition (5% $pCO_2$ (v/v), 120 µE); Black circles indicate high light condition (0.05% $pCO_2$ (v/v), 500 µE). Dotted lines shows traditional model solution with $C_i$ uptake as 100% $CO_2$; solid line shows $C_i$ uptake as 100% $HCO_3^-$; dashed line shows $C_i$ uptake as 50% $CO_2$, 50% $HCO_3^-$. The $\varepsilon_{Rubisco}$ values used for WT and ANC were 25.18‰ and 17.23‰ respectively. Solid red line indicates where $\varepsilon_P = \varepsilon_{Rubisco}$. We use the same $\varepsilon_{equi}$ value of -9‰ as used in (31). All analyses were performed using R Statistical Software (v4.1.0; R Core Team 2021) and figures were produced using the ggplot2 package (v3.3.6; Wickham, 2016).

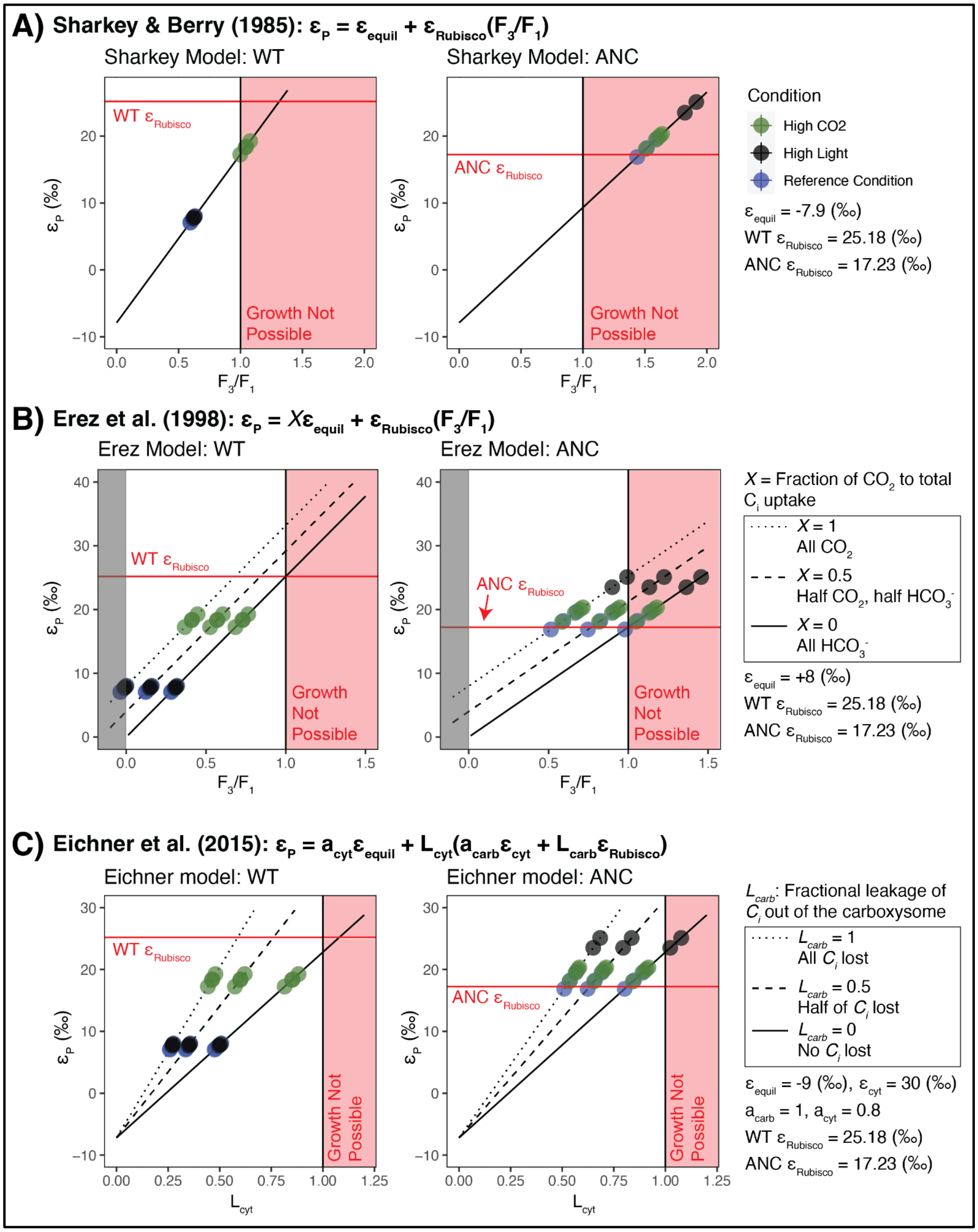

**Figure S9: WT and ANC data fit with other models.** Measured $\varepsilon_P$ values for each strain (circles) were fit with the A) Sharkey and Berry model (25), B) Erez et al. model (42), and C) Eichner et al. model (31). For all models, the $\varepsilon_{Rubisco}$ values used for WT and ANC were 25.18‰ and 17.23‰ respectively, and the solid red line indicates where $\varepsilon_P = \varepsilon_{Rubisco}$. For all models, Blue circles indicate reference condition (ambient $pCO_2$ (0.05% (v/v)), standard light flux (120 µE)); Green circles indicate high $CO_2$ condition (5% $pCO_2$ (v/v), 120 µE); Black circles indicate high light condition (0.05% $pCO_2$ (v/v), 500 µE). For all models, the red-shaded zone indicates leakage values >1. A) We used the same $\varepsilon_{equil}$ value (-7.9‰) used in (25). $F_3/F_1$ indicates leakage of $C_i$ from cell. B) We used the same $\varepsilon_{equil}$ value (+8‰) used in (42). Negative leakage values are shaded in gray. Dotted lines shows solution with $C_i$ uptake as 100% $CO_2$; solid line shows $C_i$ uptake as 100% $HCO_3^-$; dashed line shows $C_i$ uptake as 50% $CO_2$, 50% $HCO_3^-$. C) We use the same $\varepsilon_{equil}$ value (-9‰) used in (31). The values chosen for $\varepsilon_{cyt}$, $a_{carb}$, and $a_{cyt}$ are from Scenario 5 in Table 2 in (31); see text for discussion. Dotted lines shows solution where all $C_i$ taken into the carboxysome leaks out; solid line shows solution where all $C_i$ taken into the carboxysome is fixed by rubisco; dashed line shows where half of $C_i$ taken into the carboxysome is fixed. $L_{cyt}$, on the x-axis, is leakage of $C_i$ from the cell. All analyses were performed using R Statistical Software (v4.1.0; R Core Team 2021) and figures were produced using the ggplot2 package (v3.3.6; Wickham, 2016).

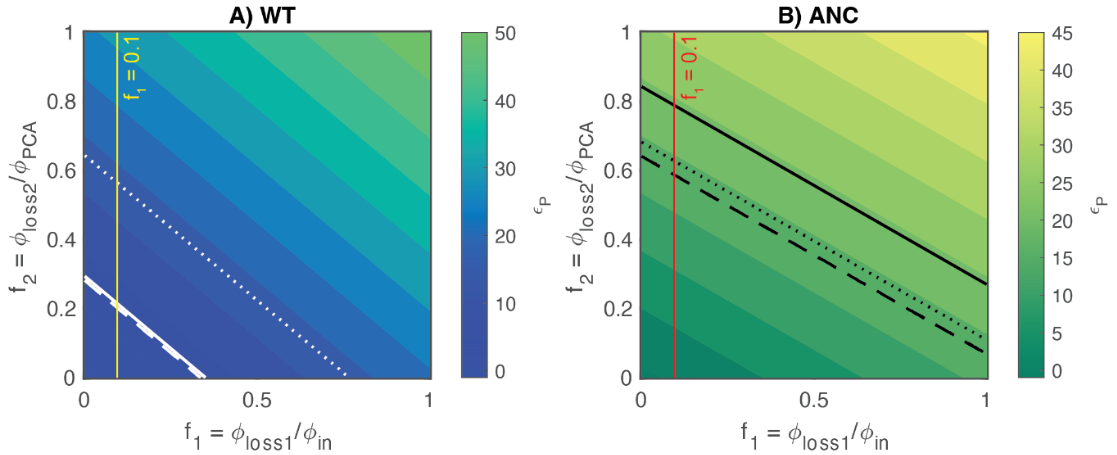

**Figure S10:** Full model outputs for the proposed box model for A) WT and B) ANC. $f_1 = 0.1$ is denoted with either a yellow or red solid line for WT or ANC respectively . Mean experimental $\varepsilon_p$ values for each condition are shown as diagonal lines as follows: 1) Dashed line is the reference condition; 2) Dotted line is the high $CO_2$ condition; 3) Solid line is the high light condition. Analyses were performed using MATLAB and Statistics Toolbox (vR2020b).

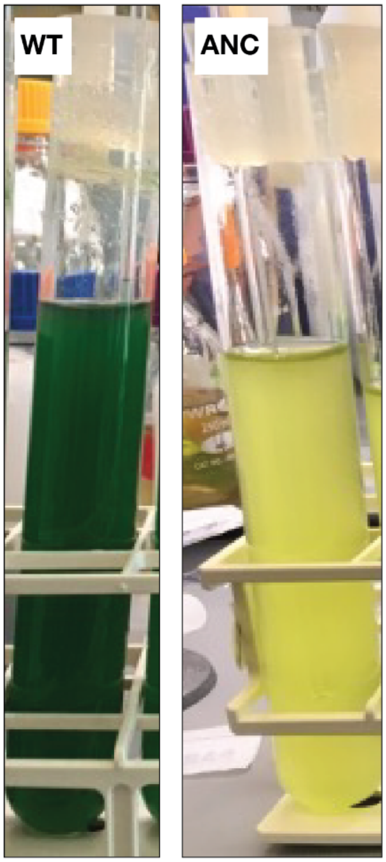

**Fig S11.** Photo showing WT strain (left) and ANC strain (right) at the end of Condition 3 growth conditions. Note yellow-green color indicative of chlorosis.

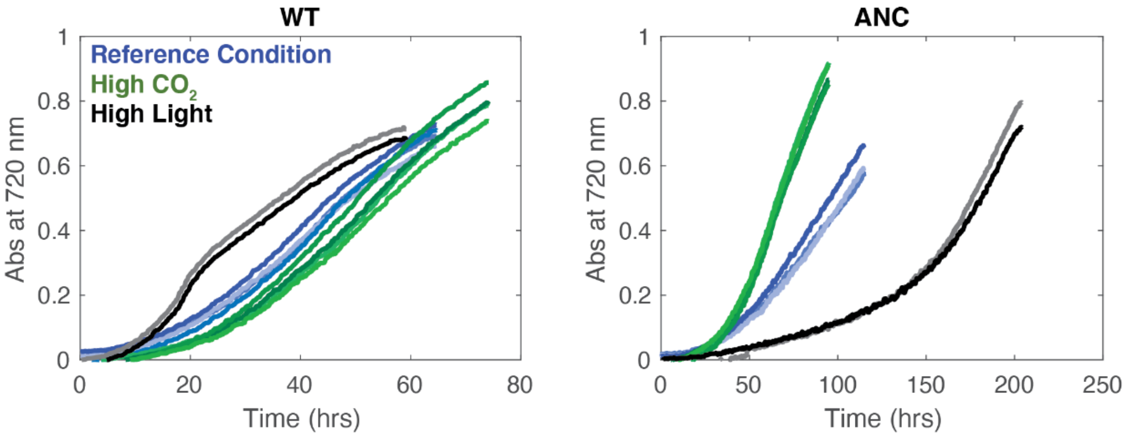

**Fig. S12.** Full growth curves for WT and ANC strains. Analyses were performed using MATLAB and Statistics Toolbox (vR2020b).

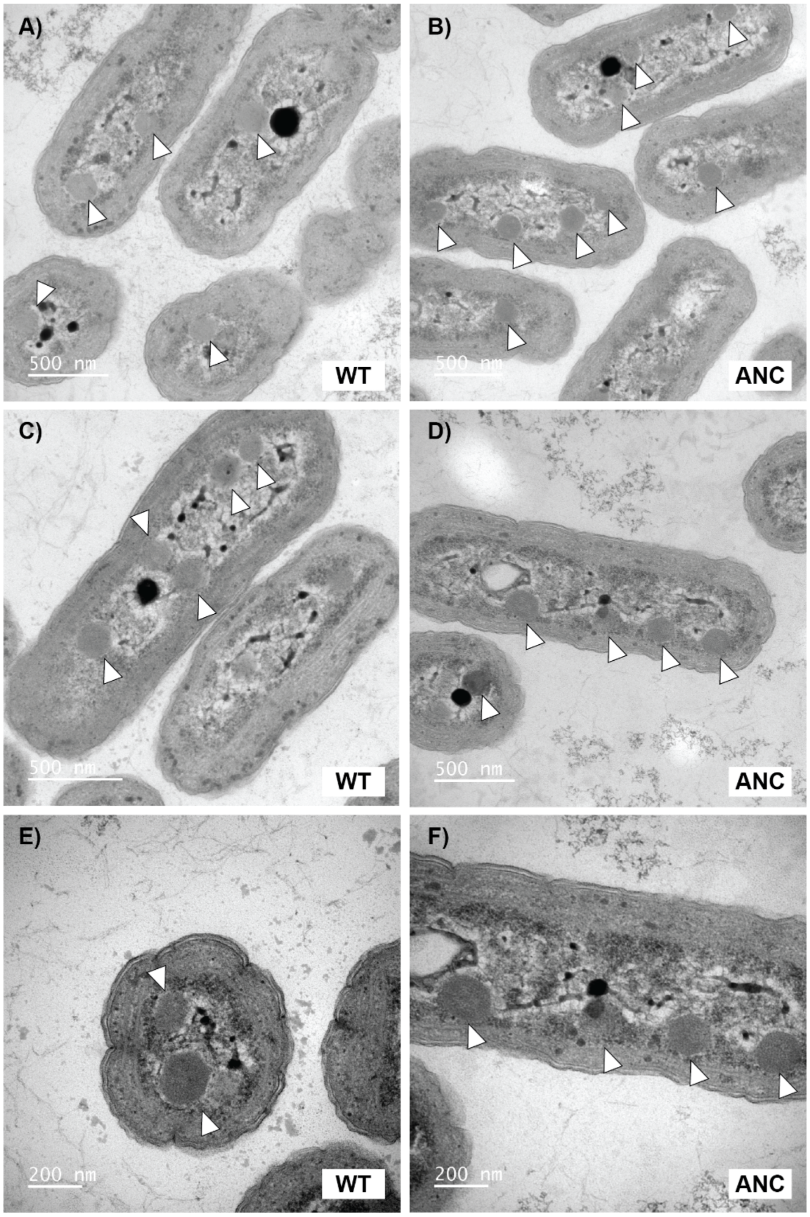

**Figure S13: Additional TEM Images of WT and ANC strains showing carboxysomes and similar cell shape and size.** Transmission electron microscopy (TEM) images show WT (A,C,E) and ANC (B,D,F) strains that were harvested mid-log phase while growing at ambient $pCO_2$ and normal light conditions (see Methods). Both strains show multiple carboxysomes per cell, as indicated by white arrows, and carboxysomes exhibit classic hexagon shape (49). The dark internal bodies are likely polyphosphate bodies (50). WT Image C) is main text Figure 4A; ANC Image D is main text Figure 4B,C.

**Rubisco Large Subunit**

```
WTRbcL     MPKTQSAAGYKAGVKDYKLTYYTPDYTPKDTDLLAAFRFSPQPGVPADEAGAAIAAESST   60
ANCRbcL    MTKTQSAAGYKAGVKDYRLTYYTPDYTPKDTDLLAAFRVTPQPGVPPEEAGAAVAAESST   60
           * ***************:********************.:****** :*****:******

WTRbcL     GTWTTVWTDLLTDMDRYKGKCYHIEPVQGEENSYFAFIAYPLDLFEEGSVTNILTSIVGN   120
ANCRbcL    GTWTTVWTDLLTDMDRYKGRCYHIEPVPGEDNSYFAFIAYPLDLFEEGSVTNILTSIVGN   120
           *******************:******* **:*****************************

WTRbcL     VFGFKAIRSLRLEDIRFPVALVKTFQGPPHGIQVERDLLNKYGRPMLGCTIKPKLGLSAK   180
ANCRbcL    VFGFKALRALRLEDIRFPVAYVKTFQGPPHGIQVERDKLNKYGRPLLGCTIKPKLGLSAK   180
           ******:*:*********** ***************** *******:*************

WTRbcL     NYGRAVYECLRGGLDFTKDDENINSQPFQRWRDRFLFVADAIHKSQAETGEIKGHYLNVT   240
ANCRbcL    NYGRAVYECLRGGLDFTKDDENINSQPFQRWRDRFLFVADAIHKAQAETGEIKGHYLNVT   240
           ********************************************:***************

WTRbcL     APTCEEMMKRAEFAKELGMPIIMHDFLTAGFTANTTLAKWCRDNGVLLHIHRAMHAVIDR   300
ANCRbcL    APTCEEMMKRAEFAKELGMPIIMHDFLTAGFTANTTLAKWCRDNGVLLHIHRAMHAVIDR   300
           ************************************************************

WTRbcL     QRNHGIHFRVLAKCLRLSGGDHLHSGTVVGKLEGDKASTLGFVDLMREDHIEADRSRGVF   360
ANCRbcL    QKNHGIHFRVLAKCLRLSGGDHLHTGTVVGKLEGDRASTLGFVDLLREDYIEADRSRGIF   360
           *:**********************:**********:*********:***:********:*

WTRbcL     FTQDWASMPGVLPVASGGIHVWHMPALVEIFGDDSVLQFGGGTLGHPWGNAPGATANRVA   420
ANCRbcL    FTQDWASMPGVMAVASGGIHVWHMPALVEIFGDDSVLQFGGGTLGHPWGNAPGATANRVA   420
           ***********: ***********************************************

WTRbcL     LEACVQARNEGRDLYREGGDILREAGKWSPELAAALDLWKEIKFEFETMDKL*   472
ANCRbcL    LEACVQARNEGRDLMREGGDILREAAKWSPELAAALELWKEIKFEFETVDKL*   472
           ************** **********.**********:***********:****
```

**Rubisco Small Subunit**

```
WTRbcS     MSMKTLPKERRFETFSYLPPLSDRQIAAQIEYMIEQGFHPLIEFNEHSNPEEFYWTMWKL   60
ANCRbcS    MQVWTPAKNKKYETFSYLPPLSDEQIAKQIQYILSQGWVPCVEFNEDSHPENRYWTMWKL   60
           *.: *  *::::***********.*** **:*::.**: * :****.*:**: *******

WTRbcS     PLFDCKSPQQVLDEVRECRSEYGDCYIRVAGFDNIKQCQTVSFIVHRPGRY*   111
ANCRbcS    PLFGAQDAAQVLSEVQACRKAFPNCYIRVVGFDNVKQCQCMSFIVHRPA*--   109
           ***..:.  ***.**: **. : :*****.****:**** :*******.
```

**Interface I**
Conserved in ANC
Not conserved in ANC
**Interface II**
Conserved in ANC

**Figure S14: Sequence alignment of RbcL and RbcS for the WT, extant rubisco sequence and the reconstructed ancestral rubisco sequence.** Alignment was performed using Clustal Omega (47) (48) for the large and small subunits of the extant, WT rubisco sequence (WTRbcL and WTRbcS respectively), and for the large and small subunits of the reconstructed ancestral rubisco (ANCRbcL and ANCRbcS respectively). Residues that are conserved in ANC in Interface I are shown in green, and residues that are not conserved are shown in red. All ANC residues were conserved for Interface II and are shown in blue. Residue numbering is shown with -3 offset from (46) and black diamonds are placed every ten residues to help with counting. Asterisk (*) indicates fully conserved residue between WT and ANC sequences; colon (:) indicates conservation between amino acids of strongly similar properties as; period (.) indicates conservation between groups of weakly similar properties as below; blank space indicates no conservation. A dash (-) indicates absence of amino acid. For more information on alignment calculations, see (47, 48) for more information.

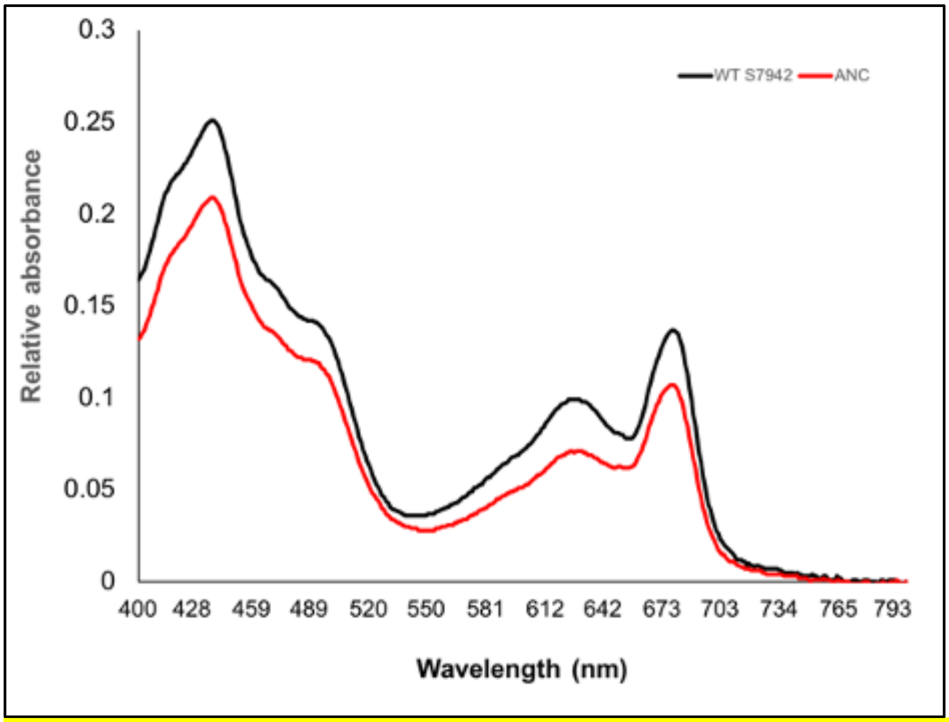

**Figure S15:** Absorption spectra of *Synechococcus elongatus* PCC 7942 wild type (black line) versus ANC mutant (red line). Absorption curves are representative of two replicates and data was normalized to values at 800 nm. Absorbance at 620nm is lower for the ANC strain indicating lower levels of phycocyanin, the major pigment of the phycobilisome, per cell compared to the wild type strain.

| Name | Sequence | Notes |
|---|---|---|

| | | |
|---|---|---|
| pAncRubisco-KanR | tcaccaataaataacgcccggcggcaaccgagcgttctgaacaaatccagatggagt<br>tctgaggtcattactggatctatcaacaggagtccaagcgagctcgatatcaaattacg<br>ccccgccctgccactcatcgcagtactgttgtaattcattaagcattctgccgacatgga<br>agccatcacaaacggcatgatgaacctgaatcgccagcggcatcagcaccttgtcgc<br>cttgcgtataatatttgcccatggtgaaaacgggggcgaagaagttgtccatattggcca<br>cgtttaaatcaaaactggtgaaactcacccagggattggctgaaacgaaaaacatatt<br>ctcaataaaccctttagggaaataggccaggttttcaccgtaacacgccacatcttgcg<br>aatatatgtgtagaaactgccggaaatcgtcgtggtattcactccagagcgatgaaaac<br>gtttcagtttgctcatggaaaacggtgtaacaagggtgaacactatcccatatcaccag<br>ctcaccgtctttcattgccatacgaaattccggatgagcattcatcaggcgggcaagaat<br>gtgaataaaggccggataaaacttgtgcttatttttctttacggtctttaaaaaggccgtaa<br>tatccagctgaacggtctggttataggtacattgagcaactgactgaaatgcctcaaaat<br>gttctttacgatgccattgggatatatcaacggtggtatatccagtgatttttttctccattttag<br>cttccttagctcctgaaaatctcgataactcaaaaaatacgcccggtagtgatcttatttca<br>ttatggtgaaagttggaacctcttacgtgcccgatcaatcatgaccaaaatcccttaacg<br>tgagttttcgttccactgagcgtcagaccccgtagaaaagatcaaaggatcttcttgaga<br>tcctttttttctgcgcgtaatctgctgcttgcaaacaaaaaaaccaccgctaccagcggtg<br>gtttgtttgccggatcaagagctaccaactctttttccgaaggtaactggcttcagcagag<br>cgcagataccaaatactgttcttctagtgtagccgtagttaggccaccacttcaagaact<br>ctgtagcaccgcctacatacctcgctctgctaatcctgttaccagtggctgctgccagtgg<br>cgataagtcgtgtcttaccgggttggactcaagacgatagttaccggataaggcgcag<br>cggtcgggctgaacggggggttcgtgcacacagcccagcttggagcgaacgaccta<br>caccgaactgagatacctacagcgtgagctatgagaaagcgccacgcttcccgaag<br>ggagaaaggcggacaggtatccggtaagcggcagggtcggaacaggagagcgc<br>acgagggagcttccagggggaaacgcctggtatctttatagtcctgtcgggtttcgcca<br>cctctgacttgagcgtcgatttttgtgatgctcgtcaggggggcggagcctatggaaaaa<br>cgccagcaacgcggcctttttacggttcctggccttttgctggccttttgctcacatgttctttc<br>ctgcgttatcccctgattctgtggataaccgtagggcgcgcctgcaggcggccgcgaat<br>tggtcctgtactgcgatcgtgcaaggcacggtttctaatgtgaccgttgcggtcgaagcc<br>gggatgtatgccgctgagcggatcggccagctcaacgcaatcatggtcattcccagac<br>cgctagacgacttgatggacagcttgcctgagccgcagtcggatagcgaagcagccc<br>agccactccaattaccgctgcgggttcgcgaaaaacaaccgctgttggagctaccgg<br>aactcgaacggcagccgatcgcgatcgaagcaccgcgacttttagcagaagagcga<br>cagtctgcgttggaattggctcaagagacaccgctcgccgagcccttagagctcccca<br>atcctcgtgatgatcagtgatggaaaaagcactgtaattcccttggtttttggctgaaagtt<br>tcggactcagtagacctaagtacagagtgatgtcaacgccttcaagctagacgggag<br>gcggcttttgccatggttcagcgatcgctcctcatcttcaataagcagggcatgagccag<br>cgttaagcaaatcaaatcaaatctcgcttctgggcttcaataaatggttccgattgatgat<br>aggttgattcatgaggaatctaaggcttaattctccacaaaagaattaagcgtccgtcgc<br>aacggaatgctccgctggacttgcgctgtgggactgcagctttacaggctccccctgcc<br>agaaatcctgaatcgtcgagcatatctgacatatctctagggagagacgacatgacca<br>aaacccagagcgccgccggttacaaagccggtgtgaaggattatcgcctgacctact<br>ataccccgattacaccccgaaggataccgatctgctggctgcctttcgcgtgaccccg<br>cagcccggtgtgccgcccgaagaggccggcgcggctgtggcggccgaaagcagc<br>acgggcacctggaccaccgtctggaccgatctgctgaccgatatggatcgctacaaa<br>ggccgctgctatcacattgaacccgtgccgggcgaggataacagctactttgccttcat<br>cgcgtatcccctggatctgttcgaagagggcagcgtcaccaacatcctgaccagcatt<br>gtgggcaatgtctttggcttcaaagctctgcgcgccctgcgcctggaagatattcgctttc<br>cggtggcctatgtcaagacctttcaaggcccgccccacggcatccaagtggagcgcg<br>ataaactgaacaagtatggccgccccctgctgggctgcaccattaaaccgaagctgg<br>gcctgagcgccaaaaattacggccgcgctgtctatgaatgcctgcgcggcggcctgg<br>attttaccaaggatgatgagaacatcaatagccagcccttccaacgctggcgcgatcg<br>ctttctgttcgtggcggatgctattcacaaagcccaggcggaaaccggcgagatcaag<br>ggccattacctgaacgtcaccgctcccacctgcgaagagatgatgaaacgcgcgga<br>atttgctaaggagctgggcatgccgatcattatgcacgattttctgaccgcgggcttcacc | Plasmid used to generate β ancestral RuBisCO strain-KanR |

| | |
|---|---|
| | gctaacaccaccctggccaaatggtgccgcgataatggcgtgctgctgcacatccatc<br>gcgccatgcatgcggtcattgatcgccagaaaaatcacggcatccattttcgcgtgctg<br>gctaagtgcctgcgcctgagcggcggcgatcacctgcatacgggtacggtggtgggt<br>aaactggaaggcgatcgcgccagcaccctgggttttgtcgatctgctgcgcgaagatt<br>atattgaggctgatcgcagccgcggcatctttttcacccaagattgggctagcatgccgg<br>gtgtgatggctgtggctagcggcggcatccacgtctggcatatgccggccctggtcgaa<br>atctttggcgatgatagcgtgctgcagtttggtggcggcaccctgggtcacccatggggt<br>aatgctcccggcgccacggctaatcgcgtggctctggaagcctgcgtccaagctcgca<br>atgagggtcgcgatctgatgcgcgagggcggcgatattctgcgcgaagctgcgaagt<br>ggagcccggagctggcggctgccctggaactgtggaaagagatcaagtttgaatttga<br>aaccgtcgataagctgtaaggagcctctgactatcgctggggagtgagcgttgctgc<br>gtaaagctttctccccagcctttcgacttaacctttcaggatttctgaatcatgcaagtgtgg<br>acccccgcgaagaacaagaagtacgaaaccttcagctacctgcccccgctgagcga<br>tgagcagatcgctaagcagatccaatacattctgagccaaggctgggtgccctgcgtc<br>gaatttaacgaggatagccacccggaaaatcgctattggaccatgtggaaactgccg<br>ctgtttggtgctcaggatgcggcccaagtgctgagcgaggtccaagcttgccgcaaag<br>cctttccgaactgctacatccgcgtggtcggcttcgataatgtgaagcagtgccaatgca<br>tgagcttcattgtccatcgcccggcgtaaagcctgatttgtcttgatagctgctcctgccttt<br>gggcaggggctttttctgtctgccattcttgaggaagtaagcttagatcgacctgcaggg<br>gggggggggaaagccacgttgtgtctcaaaatctctgatgttacattgcacaagataa<br>aaatatatcatcatgaacaataaaactgtctgcttacataaacagtaatacaaggggtg<br>ttatgagccatattcaacgggaaacgtcttgctcgaggccgcgattaaattccaacatg<br>gatgctgatttatatgggtataaatgggctcgcgataatgtcgggcaatcaggtgcgac<br>aatctatcgattgtatgggaagcccgatgcgccagagttgtttctgaaacatggcaaag<br>gtagcgttgccaatgatgttacagatgagatggtcagactaaactggctgacggaattt<br>atgcctcttccgaccatcaagcattttatccgtactcctgatgatgcatggttactcaccac<br>tgcgatccccgggaaaacagcattccaggtattagaagaatatcctgattcaggtgaa<br>aatattgttgatgcgctggcagtgttcctgcgccggttgcattcgattcctgtttgtaattgtc<br>cttttaacagcgatcgcgtatttcgactcgctcaggcgcaatcacgaatgaataacggtt<br>tggttgatgcgagtgattttgatgacgagcgtaatggctggcctgttgaacaagtctgga<br>aagaaatgcataagcttttgccattctcaccggattcagtcgtcactcatggtgatttctca<br>cttgataaccttatttttgacgaggggaaattaataggttgtattgatgttggacgagtcgg<br>aatcgcagaccgataccaggatcttgccatcctatggaactgcctcggtgagttttctcct<br>tcattacagaaacggctttttcaaaaatatggtattgataatcctgatatgaataaattgca<br>gtttcatttgatgctcgatgagtttttctaatcagaattggttaattggttgtaacactggcag<br>agcattacgctgacttgacgggacggcggctttgttgaataaatcgaacttttgctgagtt<br>gaaggatcagatcacgcatcttcccgacaacgcagaccgttccgtggcaaagcaaa<br>agttcaaaatcaccaactggtccacctacaacaaagctctcatcaaccgtggctccctc<br>actttctggctggatgatggggcgattcaggcctggtatgagtcagcaacaccttcttca<br>cgaggcagacctcagcgcccccccccccctgcaggtctggcggactctttccctttgct<br>ctacgcccatgaatgcgatcgcagtctcccctgtccagcacgttggagtgattggtggtg<br>gccagttagcttggatgctggcaccagcagcgcaacagttggggatgtcgctgcacgt<br>tcaaacacccaatgatcacgacccagcagtagcgatcgcggatcaaaccgtattagc<br>agcagttgctgacgctgcagcgactgcgaaattggctcaagcctgtgacgtcatcacat<br>tcgaaaatgagtttgttgatctgccggctttgaccgagctggaggaaactggtgtccggtt<br>tcgcccccgtccagcggcgatcgcctccctgctcgacaaacttgatcagcgacaacta<br>ttgactcgtctgggattgccaaccccacgctttttagcgatcgcggcagcaaccgcaac<br>agagtcggagctaacagccttgggctttccggtggtgctgaagcaacgccgccatggc<br>tacgacggcaagggaacacaggttttgcgatcgctagcagaacttcaacaagccttg<br>cagtcttatggcgatacgccactactcctcgaagagttcattccctttgagcaggaattag<br>cggtgatggttgcccgtagtcagagtggggcgatcgcgactttccctgtggttcagaccc<br>atcagcagaatcaggtctgtcgttgggtcgttgctcctgctgccatcccaggcgcgttgc<br>aaaaagccgttgctgcgatcgcccgaaccctcgttgagacggtcgattatgttggcgtc<br>gcgggcattgaactctttcagcagggcgatcgcctctgggtgaacgaaattgcgcccc<br>gcacccacaactcaggacactacagcttggacgcctgccagacttcgcagtttgaac |

| | |
|---|---|
| | agcagttgcgagcgatcgctgatctgcctttgggatcgacagcattgcagtggcccggtgccttaatggttaatctcctcggcttcgaggatcaccgctggatctcctgcaggcggccgcggcgcgcc |

| | | |
|---|---|---|
| pNS2-KanR | cacaccacgtctcacccttcacacaggaaacagaccatgtccagagaccattaatgc<br>agctggcacgacaggtttcccgactggaaagcgggcagtgagcgcaacgcaattaa<br>tgtgagttagctcactcattaggcaccccaggctttacactttatgcttccggctcgtatgtt<br>gtgtggaattgtgagcggataacaatttcacacaggaaacagctatgaccatgattac<br>gccaagcttgcatgcctgcaggtcgactctagaggatccccgggtaccgagctcgaat<br>tcactggccgtcgttttacaacgtcgtgactgggaaaaccctggcgttacccaacttaat<br>cgccttgcagcacatccccctttcgccagctggcgtaatagcgaagaggcccgcacc<br>gatcgcccttcccaacagttgcgcagcctgaatggcgaatggcgcctgatgcggtatttt<br>ctccttacgcatctgtgcggtatttcacaccgcatatggtgcactctcagtacaatctgctc<br>tgatgccgcatagttaagccagccccgacacccgccaacacccgctgacgcgccct<br>gacgggcttgtctgctcccggcatccgcttacagacaagctggtctctagcggttaaga<br>gaagattttcagcctgatacagattaaatcagaacgcagaagcggtctgataaaaca<br>gaatttgcctggcggcagtagcgcggtggtcccacctgaccccatgccgaactcaga<br>agtgaaacgccgtagcgccgatggtagtgtggggtatccccatgcgagagtagggaa<br>ctgccaggcatcaaataaaacgaaaggctcagtcgaaagactgggcctttcgttttatc<br>tgttgtttgtcggtgaacgctctcctgagtaggacaaatccgccgggagcggatttgaac<br>gttgcgaagcaacggcccggagggtggcgggcaggacgcccgccataaactgcca<br>ggcatcaaattaagcagaaggccatcctgacggatggccttttagtaagcttagatcga<br>cctgcagggggggggggggaaagccacgttgtgtctcaaaatctctgatgttacattgca<br>caagataaaaatatatcatcatgaacaataaaactgtctgcttacataaacagtaatac<br>aaggggtgttatgagccatattcaacgggaaacgtcttgctcgaggccgcgattaaatt<br>ccaacatggatgctgatttatatgggtataaatgggctcgcgataatgtcgggcaatca<br>ggtgcgacaatctatcgattgtatgggaagcccgatgcgccagagttgtttctgaaacat<br>ggcaaaggtagcgttgccaatgatgttacagatgagatggtcagactaaactggctga<br>cggaatttatgcctcttccgaccatcaagcattttatccgtactcctgatgatgcatggttac<br>tcaccactgcgatccccgggaaaacagcattccaggtattagaagaatatcctgattca<br>ggtgaaaatattgttgatgcgctggcagtgttcctgcgccggttgcattcgattcctgtttgt<br>aattgtccttttaacagcgatcgcgtatttcgactcgctcaggcgcaatcacgaatgaat<br>aacggtttggttgatgcgagtgattttgatgacgagcgtaatggctggcctgttgaacaa<br>gtctggaaagaaatgcataagcttttgccattctcaccggattcagtcgtcactcatggtg<br>atttctcacttgataaccttatttttgacgaggggaaattaataggttgtattgatgttggacg<br>agtcggaatcgcagaccgataccaggatcttgccatcctatggaactgcctcggtgagt<br>tttctccttcattacagaaacggctttttcaaaaatatggtattgataatcctgatatgaata<br>aattgcagtttcatttgatgctcgatgagtttttctaatcagaattggttaattggttgtaacac<br>tggcagagcattacgctgacttgacgggacggcggctttgttgaataaatcgaacttttg<br>ctgagttgaaggatcagatcacgcatcttcccgacaacgcagaccgttccgtggcaaa<br>gcaaaagttcaaaatcaccaactggtccacctacaacaaagctctcatcaaccgtgg<br>ctccctcactttctggctggatgatggggcgattcaggcctggtatgagtcagcaacacc<br>ttcttcacgaggcagacctcagcgccccccccccccctgcaggtcgatctggtaacccc<br>agcgcggttgctaccaagtagtgacccgcttcgtgatgcaaaatccgctgacgatattc<br>gggcgatcgctgctgaatgccatcgagcagtaacgtggcaccccgcccctgccaagt<br>caccgcatccagactgaacagcaccaagaggctaaaacccaatcccgccggtagc<br>agcggagaactacccagcattggtcccaccaaagctaatgccgtcgtggtaaaaatc<br>gcgatcgccgtcagactcaagcccagttcgctcatgcttcctcatctaggtcacagtcttc<br>ggcgatcgcatcgatctgatgctgcagcaagcgttttccataccggcgatcgcgccgtc<br>gccctttcgctgccgtggcccgcttacgagctcgtttatcgaccacgatcgcatccaaat<br>ccgcgatcgcttcccagtccggcaattcagtctggggcgtccgtttcattaatcctgatca<br>ggcacgaaattgctgtgcgtagtatcgcgcatagcggccagcctctgccaacagcgc<br>atcgtgattgcctgcctcaacaatctggccgcgctccatcaccaagatgcggctggcat<br>tacgaaccgtagccagacggtgagcaatgataaagaccgtccgtccctgcatcaccc<br>gttctagggcctcttgcaccaaggtttcggactcggaatcaagcgccgaagtcgcctca<br>tccagaattaaaatgcgtggatctagccgcgttgctggcgtttttccataggctccgcccc<br>cctgacgagcatcacaaaaatcgacgctcaagtcagaggtggcgaaacccgacag<br>gactataaagataccaggcgtttccccctggaagctccctcgtgcgctctcctgttccga<br>ccctgccgcttaccggatacctgtccgcctttctcccttcgggaagcgtggcgctttctcat | Plasmid used to generate NS2-KanR strain |

| | | |
|---|---|---|
| | agctcacgctgtaggtatctcagttcggtgtaggtcgttcgctccaagctgggctgtgtgc<br>acgaaccccccgttcagcccgaccgctgcgccttatccggtaactatcgtcttgagtcc<br>aacccggtaagacacgacttatcgccactggcagcagccactggtaacaggattagc<br>agagcgaggtatgtaggcggtgctacagagttcttgaagtggtggcctaactacggct<br>acactagaagaacagtatttggtatctgcgctctgctgaagccagttaccttcggaaaa<br>agagttggtagctcttgatccggcaaacaaaccaccgctggtagcggtggtttttttgtttg<br>caagcagcagattacgcgcagaaaaaaaggatctcaagaagatcctttgatctgtcc<br>agcttgtcatctgccggatgaggcaaaaccctgcctacggcgcgattacatcgtccca<br>gcgcgatcgctcttactgttgatggctcgtgcttaaaaacaatgcaaacttcaccgtttca<br>gctggtgattttcgactgtgatggtgtgcttgttgatagcggaacgcatcactaatcgcgtc<br>tttgcagacatgctcaatgaactgggtctgttggtgactttggatgacatgtttgagcagttt<br>gtgggtcattccatggctgactgtctcaaactaattgagcgacggttaggcaatcctcca<br>ccccctgactttgttcagcactatcaacgccgtacccgtatcgcgttagaaacgcatcta<br>caagccgttcctggggttgaagaggctttggatgctcttgaattgccctactgtgttgcgtc<br>cagtggtgatcatcaaaagatgcgaaccacactgagcctgacgaagctctggccacg<br>atttgagggacgaatcttcagcgtgactgaagtacctcgcggcaagccatttcccgatg<br>tctttttgttggccgccgatcgcttcggggttaatcctacggcctgcgctgtgatcgaagac<br>accccccttgggagtagcggcaggcgtggcggcaggaatgcaagtgtttggctacgcg<br>ggttccatgcccgcttggcgtctgcaagaagccggtgcccatctcatttttgacgatatgc<br>gactgctgcccagtctgctccaatcgtcgccaaaagataactccacagcattgcccaat<br>ccctaacccctgctcgcgccgcaactacacactaaaccgttcctgcgcgatcgctctta<br>ctgttgatggctcgtgcttaaaaacaatgcaaccctaaccgtttcagctggtgattttcgg<br>acgatttggcttacagggataactgagagtcaacagcctctgtccgtcattgcacaccc<br>atccatgcactggggacttgactcatgctgaatcacatttcccttgtccattgggcgagag<br>gggaggggaatcttctggactcttcactaagcggcgatcgcaggttcttctacccaagc<br>agtggcgatcgcttgattgcagtcttcaatgctggcctctgcagccatcgccgccacca<br>aagcatcgtaggcgggacgttgttgctccagtaaagtcttcgcccgtaacaatccccag<br>cgactgcgtaaatccgcttcggcaggattgcgatcgagttgccgccacagttgtttccac<br>tgggcgcgatcgtcagctccccccttccacgttgccgtagaccagttgctctgccgctgca<br>ccggccatcaacacctgacaccactgttccagcgatcgctgactgagttgcccctgtgc<br>ggcttcggcttctagcgcagctgcttggaactgcacaccccccgcgaccaggttgtccttg<br>gcgcagcgcttcccacgctgagagggtgtagcccgtcacgagcgcttacagacaag<br>ctgtgaccgcctccgggagctgcatgtgtcagaggttttcaccgtcatcaccgaaacgc<br>gcgaggcagcagatcaattcgcgcgcgaaggcgaagcggcatgcatttacgttgac<br>accatcgaatggtgcaaaacctttcgcggtatggcatgatagcgcccggaagagagt<br>caattcagggttggctgagacgtggtgtg | |
| Primer RJN610 | agggcatgagccagcgttaa | Anneals upstream of RbcLS locus |
| Primer RJN611 | ggtggtgttggcggtgaaac | Anneals to WT RbcL locus |

| | | |
|---|---|---|
| Primer RJN612 | cacgcgaaaatggatgccg | Anneals to mutant ancestral RbcL |
| Primer RJN613 | gcaatcccagacgagtcaatagtt | Anneals downstream of RbcLS locus |

**Table S1.** List of primers and plasmids used in this study. The NS2-KanR strain is referred to as 'WT' in this study, while the β ancestral rubisco strain-KanR is referred to as 'ANC.'

| Strain | Replicate | Condition | Growth constant (*k*) (1/hr) | Doubling Time (hrs) |
|---|---|---|---|---|
| WT | 1 | Reference Condition | 0.0557 ± 0.0021 | 12.4 ± 0.5 |
| WT | 2 | Reference Condition | 0.0563 ± 0.0026 | 12.3 ± 0.6 |
| WT | 3 | Reference Condition | 0.0521 ± 0.0012 | 13.3 ± 0.3 |
| WT | 4 | Reference Condition | 0.0687 ± 0.0052 | 10.1 ± 0.8 |
| ANC | 1 | Reference Condition | 0.0342 ± 0.0020 | 20.3 ± 1.2 |
| ANC | 2 | Reference Condition | 0.0313 ± 0.0046 | 22.1 ± 3.3 |
| ANC | 3 | Reference Condition | 0.0348 ± 0.0029 | 19.9 ± 1.7 |
| WT | 1 | High $CO_2$ | 0.0535 ± 0.0029 | 13.0 ± 0.7 |
| WT | 2 | High $CO_2$ | 0.0606 ± 0.0037 | 11.4 ± 0.7 |
| WT | 3 | High $CO_2$ | 0.0618 ± 0.0025 | 11.2 ± 0.5 |
| WT | 4 | High $CO_2$ | 0.0598 ± 0.0060 | 11.6 ± 1.2 |
| ANC | 1 | High $CO_2$ | 0.0553 ± 0.0045 | 12.5 ± 1.0 |
| ANC | 2 | High $CO_2$ | 0.0614 ± 0.0026 | 11.3 ± 0.5 |
| ANC | 3 | High $CO_2$ | 0.0591 ± 0.0069 | 11.7 ± 1.4 |
| ANC | 4 | High $CO_2$ | 0.0553 ± 0.0102 | 12.5 ± 2.3 |
| WT | 1 | High Light | 0.1980 ± 0.0188 | 3.5 ± 0.3 |
| WT | 2 | High Light | 0.1874 ± 0.0144 | 3.7 ± 0.3 |
| ANC | 1 | High Light | 0.0165 ± 0.0015 | 42.0 ± 3.8 |
| ANC | 2 | High Light | 0.0125 ± 0.0019 | 55.5 ± 8.4 |

**Table S2: Fitted growth constants and doubling times for growth curves.** Outputs from MCMC approach for fitting exponential phase of growth phase (avg. ± s.d.). Doubling time was calculated as ln(2)/*k*.

| Strain | Rep. | Condition | $\delta^{13}C$ of $CO_2$ (‰) | $\delta^{13}C$ of bulk cells (‰) | $\varepsilon_P$ ($CO_2$/bio) (‰) |
|---|---|---|---|---|---|
| WT | 1 | Reference Condition | -12.455 ± 0.005 | -19.371 ± 0.043 | 7.053 ± 0.045 |
| WT | 2 | Reference Condition | -12.455 ± 0.005 | -19.850 ± 0.046 | 7.544 ± 0.048 |
| WT | 3 | Reference Condition | -12.455 ± 0.005 | -19.480 ± 0.053 | 7.165 ± 0.055 |
| WT | 4 | Reference Condition | -12.455 ± 0.005 | -20.343 ± 0.087 | 8.052 ± 0.090 |
| ANC | 1 | Reference Condition | -12.455 ± 0.005 | -31.482 ± 0.088 | 19.646 ± 0.093 |
| ANC | 2 | Reference Condition | -12.455 ± 0.005 | -30.129 ± 0.089 | 18.223 ± 0.094 |
| ANC | 3 | Reference Condition | -12.455 ± 0.005 | -28.841 ± 0.102 | 16.873 ± 0.107 |
| WT | 1 | High $CO_2$ | -36.839 ± 0.021 | -54.247 ± 0.298 | 18.407 ± 0.322 |
| WT | 2 | High $CO_2$ | -36.839 ± 0.021 | -54.162 ± 0.097 | 18.315 ± 0.108 |
| WT | 3 | High $CO_2$ | -36.839 ± 0.021 | -55.037 ± 0.572 | 19.258 ± 0.618 |
| WT | 4 | High $CO_2$ | -36.839 ± 0.021 | -53.160 ± 0.133 | 17.237 ± 0.146 |
| ANC | 1 | High $CO_2$ | -36.839 ± 0.021 | -53.924 ± 1.002 | 18.059 ± 1.079 |
| ANC | 2 | High $CO_2$ | -36.839 ± 0.021 | -55.750 ± 1.382 | 20.027 ± 1.494 |
| ANC | 3 | High $CO_2$ | -36.839 ± 0.021 | -56.029 ± 1.307 | 20.329 ± 1.413 |
| ANC | 4 | High $CO_2$ | -36.839 ± 0.021 | -55.216 ± 1.605 | 19.451 ± 1.732 |
| WT | 1 | High Light | -12.455 ± 0.005 | -20.213 ± 0.102 | 7.918 ± 0.105 |
| WT | 2 | High Light | -12.455 ± 0.005 | -20.007 ± 0.132 | 7.706 ± 0.136 |
| ANC | 1 | High Light | -12.455 ± 0.005 | -36.632 ± 0.082 | 25.097 ± 0.088 |
| ANC | 2 | High Light | -12.455 ± 0.005 | -35.131 ± 0.073 | 23.501 ± 0.077 |

**Table S3: Measured carbon isotope values ($\delta^{13}C$) and calculated $\varepsilon_P$ values.** Values (avg. ± s.e.) are reported relative to VPDB.

| Strain | Condition | $CO_2$ Concentration (%) | Light intensity ($\mu$E) | $\varepsilon_P$ ($CO_2$/bio) (‰) |
|---|---|---|---|---|
| WT | Reference Condition | 0.04 | 120 | 7.453 ± 0.124 |
| ANC | Reference Condition | 0.04 | 120 | 18.247 ± 0.170 |
| WT | High $CO_2$ | 5 | 120 | 18.304 ± 0.720 |
| ANC | High $CO_2$ | 5 | 120 | 19.467 ± 2.897 |
| WT | High Light | 0.04 | 500 | 7.812 ± 0.172 |
| ANC | High Light | 0.04 | 500 | 24.299 ± 0.117 |

**Table S4: $\varepsilon_P$ values used for Figure 2 in main text.** Values (avg. ± s.e.) are reported relative to VPDB.

| ID | Rep | time (min) | Avg δ13C | Std dev δ13C | Std err δ13C | Avg R | Std dev R | Std err R |
|---|---|---|---|---|---|---|---|---|
| Sub | 1 | 0 | -3.06 | 0.19 | 0.06 | 0.0111628 | 1.46E-06 | 4.60E-07 |
| Sub | 2 | 0 | -2.59 | 0.09 | 0.03 | 0.0111664 | 7.21E-07 | 2.28E-07 |
| Sub | 3 | 0 | -2.52 | 0.12 | 0.04 | 0.0111670 | 9.15E-07 | 2.89E-07 |
| ANC | 1 | 0 | -3.43 | 0.36 | 0.11 | 0.0111598 | 2.83E-06 | 8.96E-07 |
| ANC | 2 | 15 | -2.00 | 0.33 | 0.10 | 0.0111711 | 2.60E-06 | 8.22E-07 |
| ANC | 3 | 30 | -2.69 | 0.22 | 0.07 | 0.0111657 | 1.72E-06 | 5.43E-07 |
| ANC | 4 | 45 | -2.68 | 0.19 | 0.06 | 0.0111658 | 1.51E-06 | 4.79E-07 |
| ANC | 5 | 60 | -1.81 | 0.31 | 0.10 | 0.0111726 | 2.46E-06 | 7.78E-07 |
| ANC | 6 | 90 | -2.17 | 0.28 | 0.09 | 0.0111697 | 2.21E-06 | 6.97E-07 |
| ANC | 7 | 120 | -2.17 | 0.19 | 0.06 | 0.0111697 | 1.53E-06 | 4.85E-07 |
| ANC | 8 | 150 | -1.68 | 0.30 | 0.09 | 0.0111736 | 2.33E-06 | 7.38E-07 |
| ANC | 9 | 210 | -2.23 | 0.22 | 0.07 | 0.0111693 | 1.75E-06 | 5.53E-07 |
| ANC | 10 | 270 | -1.51 | 0.29 | 0.09 | 0.0111750 | 2.25E-06 | 7.11E-07 |
| ANC | 11 | 429 | -4.56 | 0.09 | 0.03 | 0.0111509 | 6.64E-07 | 2.10E-07 |
| WT | 1 | 0 | -7.87 | 0.23 | 0.07 | 0.0111249 | 1.80E-06 | 5.68E-07 |
| WT | 2 | 15 | -6.85 | 0.16 | 0.05 | 0.0111329 | 1.28E-06 | 4.04E-07 |
| WT | 3 | 30 | -6.03 | 0.27 | 0.08 | 0.0111394 | 2.12E-06 | 6.70E-07 |
| WT | 4 | 45 | -5.85 | 0.23 | 0.07 | 0.0111408 | 1.82E-06 | 5.75E-07 |
| WT | 5 | 60 | -5.96 | 0.28 | 0.09 | 0.0111400 | 2.23E-06 | 7.05E-07 |
| WT | 6 | 90 | -5.86 | 0.22 | 0.07 | 0.0111407 | 1.78E-06 | 5.62E-07 |
| WT | 7 | 120 | -5.86 | 0.23 | 0.07 | 0.0111407 | 1.82E-06 | 5.76E-07 |
| WT | 8 | 150 | -5.89 | 0.20 | 0.06 | 0.0111405 | 1.56E-06 | 4.94E-07 |
| WT | 9 | 210 | -6.83 | 0.13 | 0.04 | 0.0111331 | 1.04E-06 | 3.30E-07 |
| WT | 10 | 270 | -5.74 | 0.23 | 0.07 | 0.0111417 | 1.84E-06 | 5.82E-07 |
| WT | 11 | 429 | -6.48 | 0.14 | 0.04 | 0.0111359 | 1.10E-06 | 3.49E-07 |

**Table S5.** Results of the WT and ANC Rubisco assays. Avg $\delta^{13}C$ refers to the average of 10 measurement repetitions. Standard deviation (Std dev) and standard error (Std err) are calculated as described.

| | | Model Fits | | | Goodness of Fit | | | | |
|---|---|---|---|---|---|---|---|---|---|
| Strain | Fit | a | b | c | sse | rsquare | dfe | adjrsquare | rmse |
| ANC | 1 | -1.03E-05 | 0.66312 | 0.01117 | 1.19E-10 | 0.34286 | 7 | 0.15511 | 4.12E-06 |
| ANC | 2 | -1.03E-05 | 0.07952 | 0.011171 | 8.57E-11 | 0.52600 | 7 | 0.39058 | 3.50E-06 |
| ANC | 3 | -9.97E-06 | 0.03655 | 0.011171 | 1.08E-10 | 0.40244 | 7 | 0.23171 | 3.93E-06 |
| WT | 1 | -1.54E-05 | 0.68629 | 0.01114 | 2.72E-11 | 0.88130 | 5 | 0.83381 | 2.33E-06 |
| WT | 2 | -1.62E-05 | 0.04995 | 0.011141 | 7.11E-12 | 0.96901 | 5 | 0.95662 | 1.19E-06 |
| WT | 3 | -1.63E-05 | 0.05983 | 0.011141 | 5.78E-12 | 0.97484 | 5 | 0.96477 | 1.07E-06 |

**Table S6.** Model fits for the general model y = a*EXP(-b*x)+c. sse = Sum of Squares Due to Error or summed square of residuals. rsquare = R-Square value. dfe = Residual Degrees of Freedom. adjrsquare = Degrees of Freedom Adjusted R-Square. rmse = Root Mean Squared Error.

| Strain | Fit | m | 95% ci | 95% ci | b | 95% ci | 95% ci |
|---|---|---|---|---|---|---|---|
| ANC | 1 | 16.23 | 16.05 | 16.42 | 0.009900 | -0.000475 | 0.020275 |
| ANC | 2 | 17.12 | 16.94 | 17.30 | 0.009391 | -0.000460 | 0.019243 |
| ANC | 3 | 18.33 | 18.15 | 18.52 | 0.008776 | -0.000441 | 0.017994 |
| WT | 1 | 24.56 | 24.39 | 24.72 | 0.004023 | -0.004336 | 0.012382 |
| WT | 2 | 25.53 | 25.36 | 25.70 | 0.003873 | -0.004178 | 0.011923 |
| WT | 3 | 25.46 | 25.29 | 25.62 | 0.003884 | -0.004189 | 0.011956 |

**Table S7.** Fit results of Rayleigh curve. 95% ci = 95% Confidence Interval. m and b are the constants in the model y = m*x+b.

| **Interface I Amino Acids** | | | |
|---|---|---|---|
| Rubisco subunit | Wang et al. (2019) reported residue number | Residue number with offset | Present in reconstructed ancestral rubisco? |
| RbcL | Asp76 / D76 | Asp73 / D73 | Yes |
| RbcL | Arg79 / R79 | Arg76 / R76 | Yes |
| RbcL | Glu351 / E351 | Glu348 / E348 | Yes |
| RbcL | His353 / H353 | His350 / H350 | No |
| RbcL | Glu355 / E355 | Glu352 / E352 | Yes |
| RbcS | Gln36 / Q36 | *N/A* | Yes |
| RbcS | Gly37 / G37 | *N/A* | Yes |
| RbcS | Asp93 / D93 | *N/A* | Yes |
| RbcS | Asn94 / N94 | *N/A* | Yes |

| | | | |
|---|---|---|---|
| RbcS | Ile95 / I95 | *N/A* | No |

**Table S8: Contact residues between RbcL, RbcS, and SSUL at Interface I in *Synechococcus elongatus* PCC 7942.** Interface I involves both the large (RbcL) and small (RbcS) subunits of rubisco. Numbered amino acids are taken from Figure 4c of Wang et al. (Wang et al. 2019) There is an offset of -3 between the numbering of Wang et al. and our WT sequence for RbcL. There is no offset for RbcS. We first converted the reported residue number to the offset number before looking for the residue in our sequence.

| **Interface II Amino Acids** | | | |
|---|---|---|---|
| Rubisco subunit | Wang et al. (2019) reported residue number | Residue number with offset | Present in reconstructed ancestral rubisco? |
| RbcL | Tyr29 / Y29 | Tyr26 / Y26 | Yes |
| RbcL | Thr30 / T30 | Thr27 / T27 | Yes |
| RbcL | Pro31 / P31 | Pro28 / P28 | Yes |
| RbcL | Lys32 / K32 | Lys29 / K29 | Yes |
| RbcL | Tyr85 / Y85 | Tyr82 / Y82 | Yes |
| RbcL | His86 / H86 | His83 / H83 | Yes |

**Table S9: Contact residues between RbcL and SSUL at Interface II in *Synechococcus elongatus* PCC 7942.** Interface II only involves the large (RbcL) subunit of rubisco. Numbered amino acids are taken from Figure 4c of Wang et al. (Wang et al. 2019) There is an offset of -3 between the numbering of Wang et al. and our WT sequence for RbcL. We first converted the reported residue number to the offset number before looking for the residue in our sequence.
